# Rapid detection of *Staphylococcus aureus* and *Streptococcus pneumoniae* by real-time analysis of volatile metabolites

**DOI:** 10.1101/2022.03.16.484604

**Authors:** Alejandro Gómez-Mejia, Kim Arnold, Julian Bär, Kapil Dev Singh, Thomas C. Scheier, Silvio D. Brugger, Annelies S. Zinkernagel, Pablo Sinues

**Affiliations:** Department of Infectious Diseases and Hospital Epidemiology, University Hospital Zurich, University of Zürich, Zurich, Switzerland; University Children’s Hospital Basel (UKBB), 4056 Basel, Switzerland; Department of Biomedical Engineering, University of Basel, 4123 Allschwil, Switzerland

**Keywords:** bacterial infections, clinical diagnostics, real-time mass spectrometry, SESI-HRMS, *Staphylococcus aureus*, *Streptococcus pneumoniae*

## Abstract

Rapid detection of pathogenic bacteria is needed for rapid diagnostics allowing adequate and timely treatment. In this study, we aimed to evaluate the technical feasibility of Secondary Electro-Spray Ionization-High Resolution Mass Spectrometry (SESI-HRMS) as a diagnostic tool for rapid detection of bacterial infections and compare its performance with the current standard of diagnostics. We compared the time required to confirm growth of the pathogenic bacteria *Staphylococcus aureus* and *Streptococcus pneumoniae* by conventional detection by culture and MAL-DI-TOF vs. detection of specific volatile organic compounds (VOCs) produced by these human pathobionts. SESI-HRMS could consistently detect VOCs produced by *S. aureus* or *S. pneumoniae* on blood agar plates within minutes, allowing to positively identify bacteria within hours. Unique *S. aureus* and *S. pneumoniae* features were detected already at bacterial densities as low as ∼10^3^ colony forming units. Rich mass spectral fingerprints allowed for the distinction of these two bacteria on a species and even strain level. To give an incentive towards clinical application of this technology, further analyzed 17 clinical samples previously diagnosed by conventional methods. We predominantly obtained a separation of samples which showed growth (i.e. presence of living bacteria) compared to samples with no bacterial growth (i.e. presence of dead bacteria). We conclude that SESI-HRMS allows rapid identification of unique bacterial features. Further development of real-time analysis of clinical samples by SESI-HRMS will shorten the time required for microbiological diagnosis with a high level of confidence and sensitivity and should help to improve patient’s tailored treatment.

**IMPORTANCE:** A timely identification of a pathogenic bacteria causing the infection is of pivotal importance for the initiation of an adequate antimicrobial therapy. In this regard, different technologies have been developed with the aim to achieve a highly reliable, specific, and overall fast identification of pathogenic bacteria. However, conventional diagnostic techniques still require long preprocessing times (hours to days) to acquire enough biological material for an accurate identification of the pathogen. Therefore, in this work, we aimed to further shorten the detection time of current gold standards for microbiological diagnostics by providing a system capable of a fast, sensitive and specific discrimination of different pathogenic bacteria. This system relies on the real-time mass spectrometric detection of volatile organic compounds (VOCs) produced by a given organism during its growth, potentially leading to a significant shortening of the time required to obtain a positive reliable diagnostic.

## INTRODUCTION

A prompt and accurate identification of the causative pathogens of a bacterial infection is essential for providing patients with adequate treatments to reduce mortality and to prevent antibiotic resistance (1-4). Bacterial infections caused by the human pathogenic bacteria *Staphylococcus aureus* and *Streptococcus pneumoniae* remain highly prevalent (5-8). Despite the development and availability of antibiotics, mortality remains high, reaching 20 % for *S. aureus* associated endocarditis and more than one million deaths of children below 5 years of age by *S. pneumoniae* (5, 7).

Currently, state-of-the-art bacterial identification methods consist of different strategies: in addition to conventional growth based diagnostics, molecular methods (16S-rRNA, whole genome sequencing and antigen detection) and Matrix Assisted Laser Desorption/Ionization-Time of Flight (MALDI-TOF) mass spectrometry, are the current culture based gold standards for identification of bacteria (9-12). Both methods portray complementary properties. Molecular methods allow for detection of single bacterial components such as pneumococcal antigen or DNA directly in a clinical sample. This translates into a similar diagnostic tool comparable to MALDI-TOF, at the cost of not being able to differentiate between live and dead bacteria or to present limitations to identify antibiotic susceptibilities (13). On the other hand, the rich peptide fingerprints detected by MALDI-TOF, allow for a very high degree of specificity, covering a vast range of pathogens. However, this gold-standard method still bears limitations. In order to be able to identify a bacterium, a minimum concentration of bacteria have to be grown for a defined time to allow an accurate identification of the sample (9, 10, 14-16). Such prerequisite introduces additional time for diagnosis of approximately 16 to 24 hours until the bacteria have grown sufficiently (17-19).

Additional experimental methods are being developed to overcome these limitations of the techniques currently accepted in microbiological diagnostics. One such an approach is to detect volatile metabolites (i.e. volatile organic compounds aka VOCs) released by microorganisms as they grow (20, 21). By doing so, the limitations of molecular methods such as time to positive detection and distinguishing between dead and alive bacteria is overcome. In addition, similarly to the specificity provided by the peptide profiles captured by MALDI-TOF, the unique metabolism developed over millions of years of evolution of bacteria provides an opportunity to render a high specificity. This strategy has been usually pursued by different mass spectrometric variants. The workhorse of VOCs analysis released by bacteria has been gas chromatography-mass spectrometry (GC-MS), which allows detecting very complex gas mixtures (22). However, the method is also cumbersome, as it requires lengthy sample preparation, limiting the possibilities of providing quicker results than MALDI-TOF-based analyses. Alternative real-time mass-spectrometric techniques such as selected ion flow tube-mass spectrometry (SIFT-MS) (23-26) and Secondary Electro-Spray Ionization-High Resolution Mass Spectrometry (SESI-HRMS) (27-37) can provide analyses of VOCs on the fly, thus potentially shortening time-to-results.

Despite the encouraging results and evidence accumulated over decades of analysis of VOCs released by pathogens, such analytical strategy did not make it to transition from a research level to a commercially available solution. Some of the reasons include that it is unclear yet whether the sensitivity of such methods is enough to detect bacterial growth during very early stages of bacterial replication, hence potentially accelerating positive results to just a few hours. Another remaining question is whether at such low VOCs concentration levels, the selectivity is enough to enable species differentiation. In this work, we addressed these open questions using SESI-HRMS, which features limits of detection for VOC features as low a part-per-trillion (38). Additionally, SESI-HRMS allows for the simultaneous detection of hundreds of VOCs from microorganisms because of the high resolution of the mass analyzer (39). To do so, we conducted quantitative measurements of two different *S. aureus* and *S. pneumoniae* strains. Parallel image analysis and gold-standard culture and MALDI-TOF measurements were used to benchmark the technology. We culminated the study providing proof-of-principle on the feasibility of this sample preparation-free approach to ‘sniff-out’ a variety of clinical samples.

## RESULTS

### Sensitivity measurements: quantifying the detection limit of colony forming units (CFUs)

To evaluate the sensitivity of SESI-HRMS for the detection of bacterial VOC features, we measured a low number of CFUs (140 – 2000 CFUs per plate) of *S. aureus* and *S. pneumoniae* strains over ∼ 15 hours with SESI-HRMS. In parallel, time-lapse (TL) images were acquired to record bacterial growth over 15 h and up to 38 h for certain replicates (Fig 1 and Fig S1). As an example, figure 1A shows a representation of a time trace for a mass spectral feature observed in *S. aureus* Cowan1 during SESI-HRMS measurement at a mass-to-charge ratio (*m/z)* 144.0476. Complementary TL pixel intensities for each replicate are presented in figure 1B, accompanied by control measurements together with TL image row examples at 15 h, 24 h and 38 h of measurement. No bacteria were visually detected by the TL system by the end of SESI-HRMS acquisition at 15 h. In contrast, *m/z* 144.0476 in *S. aureus* Cowan1 was detected by SESI-HRMS in all four replicates within the first minutes of growth/measurement even if CFU numbers are as low as 140 CFUs (Fig 1A). To control for bacterial growth, the plates were further analyzed for time lapse beyond the 15 h analysis with SESI-HRMS, confirming the appearance of colonies after 24 h of growth (Fig 1B). In general, growth was detected for all bacterial strains under low CFU condition as well as under high CFU condition (Fig S1).

**FIG 1.**
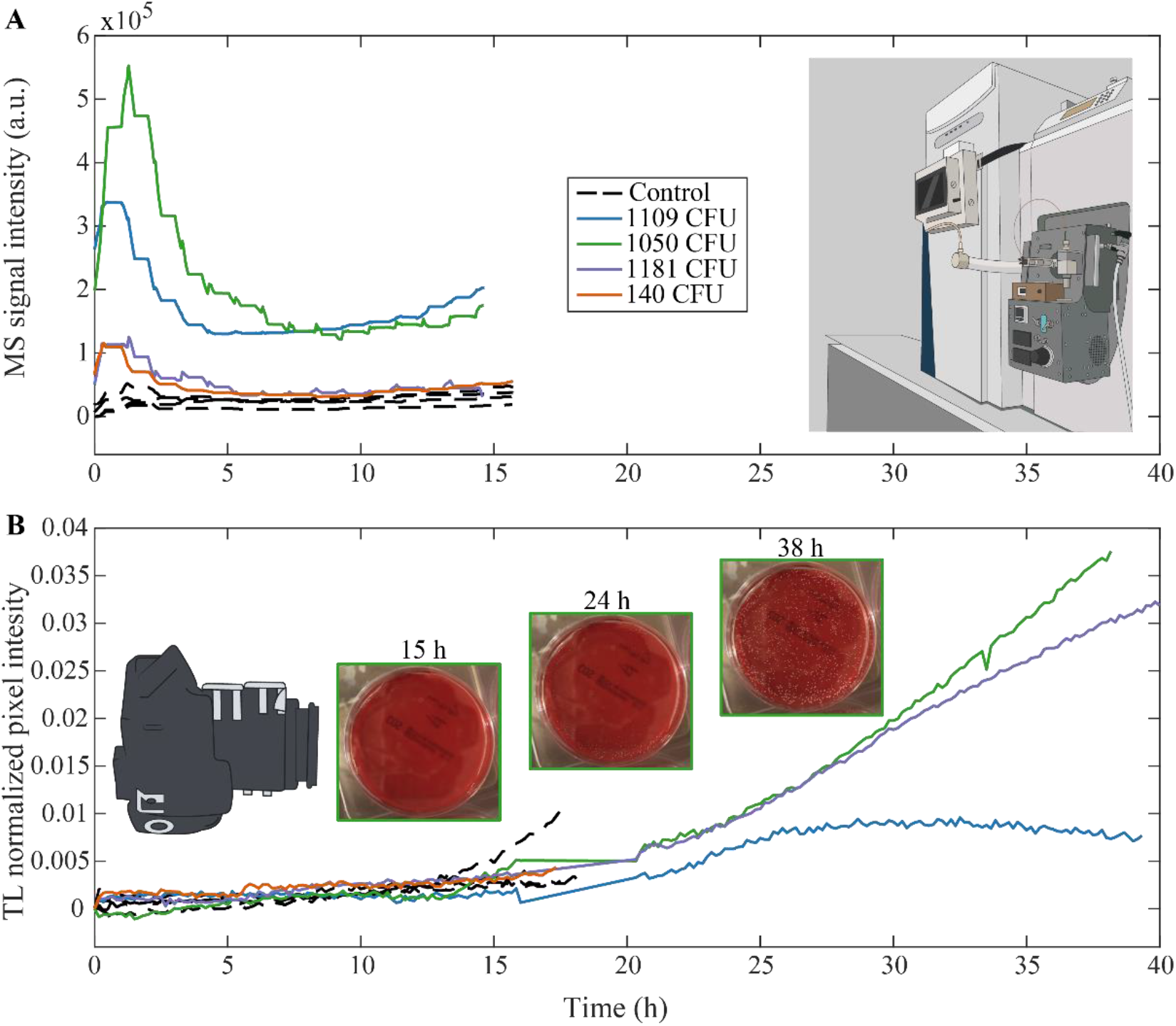
Detection of features in headspace of growing bacterial culture by SESI-HRMS vs. visual monitoring by TL camera. (A) Example of a feature time trace at *m/z* 144.0476 observed in *S. aureus* Cowan1 by SESI-HRMS over 15 h of measurement. (B) Corresponding TL normalized pixel intensity of the measured replicates along with pictures showing visually captured growth at 15 h, 24 h and 38 h after the bacteria were put on the plate. The four colored lines represent four biological replicates and the four dashed black lines represent the four control replicates. The colors indicate how many CFUs were put on the blood agar plate for each replicate measured. MS=Mass spectrometer, CFU = Colony forming unit, TL = Time lapse.

### Selectivity measurements: Real-time detection of unique features on species and strain level

After confirming the detection of features under a low number of CFUs, we investigated whether these detected features were attributable to the *S. aureus* strains JE2 and Cowan1 or the *S. pneumoniae* strains D39 and TIGR4. We were able to assign a total of 392 features to the two bacterial species or their respective strains as summarized in table S1. We found that out of 392 features, 51 features were *S. aureus*-specific (JE2 and Cowan1), 302 features were unique to *S. aureus* JE2 and 15 features unique to *S. aureus* Cowan1. Moreover, 18 features were identified as *S. pneumoniae*-specific (D39 and TIGR4), five features were unique to *S. pneumoniae* D39 and one feature was unique to *S. pneumoniae* TIGR4.

Since the features listed in table S1 identified for low CFUs (in the order of thousands) were not always present in all biological replicates, we aimed to evaluate the use of SESI-HRMS with high – density (billions) CFUs cultures to increase the signal strength and achieve a better reproducibility among the biological replicates. Table S2 summarizes the total of 1,269 features detected under high density conditions. Out of the total of 1,269 features, 19 features were detected in both *S. aureus* strains JE2 and Cowan1. Three specific features were only found in *S. aureus* Cowan1 and 26 features in *S. aureus* JE2. For *S. pneumoniae*, 15 specific features were present in both strains D39 and TIGR4. When both *S. pneumoniae* strains were evaluated separately, 1,206 features were unique to *S. pneumoniae* D39 and no unique features were identified for TIGR4.

Out of the 26 features assigned to the strain *S. aureus* JE2, one representative feature at *m/z* 104.1069 is shown for all biological replicates and the controls (Fig 2A). The signal of this particular feature was more abundant in all four *S. aureus* JE2 replicates with a signal intensity of ∼ 4×10^5^ (a.u.) and nearly absent among all other strains and controls (Fig 2A). A similar profile was shown for additional nine features depicted in the heat maps of S. *aureus* JE2 (Fig 2A). They start to be detectable at ∼ 5 h with increasing abundance towards the end of the measurement at 15 h. In contrast, the features remained at the baseline level for the rest of strains and for the control. Another example of *S. aureus* species specific time profiles is shown in figure 2B. Out of 19 features detected in both *S. aureus* strains JE2 and Cowan1, a relevant feature at *m/z* 101.0608 is shown for all biological replicates and controls (Fig 2B). Very similar to figure 2A, this particular feature started to increase towards the end of the measurement and was only present in both *S. aureus* strains with a high signal intensity and nearly not detected in the other strains and controls. Figure S2 shows the heatmaps and/or representative specific features identified for the remaining species and strains. Unique features were assigned to each strain and species with the exception of *S. pneumoniae* TIGR4 for which no unique features were assigned, albeit its growth was confirmed by TL (Fig S1). Nevertheless, some features were present in *S. pneumoniae* TIGR4 whereas they were absent in the control (Fig S2E).

**FIG 2.**
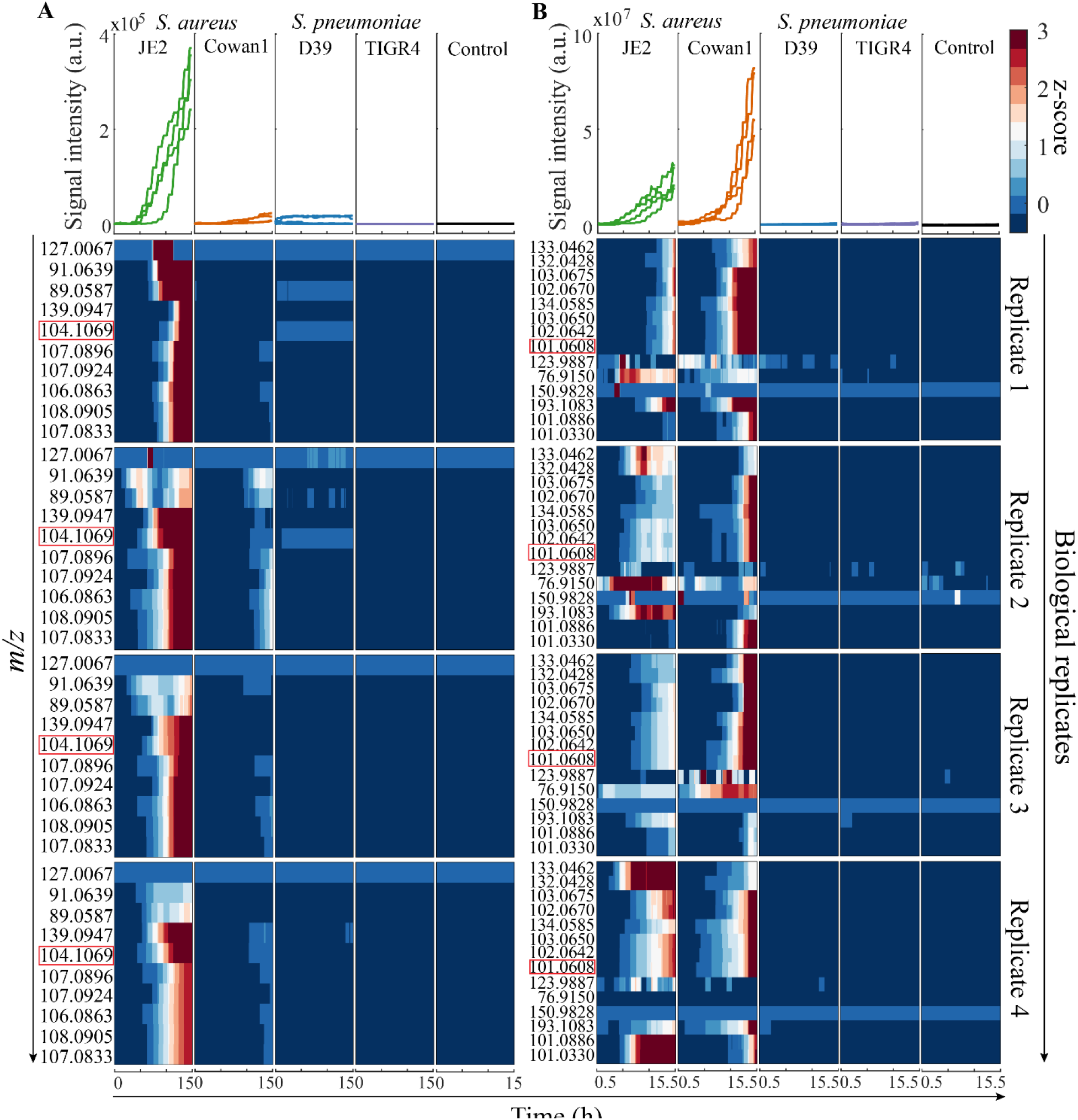
Specific time – dependent features detected during bacterial growth. (A) Example time trace of the positive ion at *m/z* 104.1069 (framed in red) unique to *S. aureus* JE2 is shown on top of the heatmaps consisting of total ten features (positive ions) unique to *S. aureus* JE2. (B) Example time trace of the negative ion at *m/z* 101.0608 (framed in red) unique to the species *S. aureus* (i.e. present in both JE2 and Cowan1) is shown on top of the heatmaps consisting of total 14 features (negative ions) unique to species *S. aureus*. Real-time evolution of all features is shown over 15 hours of measurement by SESI-HRMS for all four investigated strains (n=4 biological replicates) and controls (n=4). The color bar indicates the z-score values of absolute signal intensity for each feature, from low (dark blue) to increased signal intensity (dark red).

Furthermore, we also compared the overlap of features detected using low density culture (Table S1) versus using a high-saturated growth plate (Table S2). Only six species *S. pneumoniae* specific features were found under both conditions (Fig S3).

### Diverging metabolic trajectories of bacterial strains

Next, we investigated the evolution over time of the features produced by the different bacteria under investigation as they grew. Given the large number of features detected, we visualized our multivariate dataset using Principal Component Analysis (PCA) and dendrogram trees to obtain clusters of the different bacterial strains at different stages of bacterial growth (Fig 3, Fig S4 and Fig S5). An initial separation of *S. pneumoniae* D39 from the other strains became apparent after 15 min of measurement (Fig S4). At the time point 7 h, a separation was noted for the *S. aureus* JE2 replicates. The *S. aureus* Cowan1 group started to drift apart from the controls at 10 h after growth. The best discrimination of the different groups was observed at 12 h as shown in the PCA and dendrogram tree in figure 3. Except of *S. pneumoniae* TIGR4, all strains could be distinguished from the controls very clearly. Furthermore, it is well visible how Euclidean distance shortens as a function of growth time (Fig S5).

**FIG 3.**
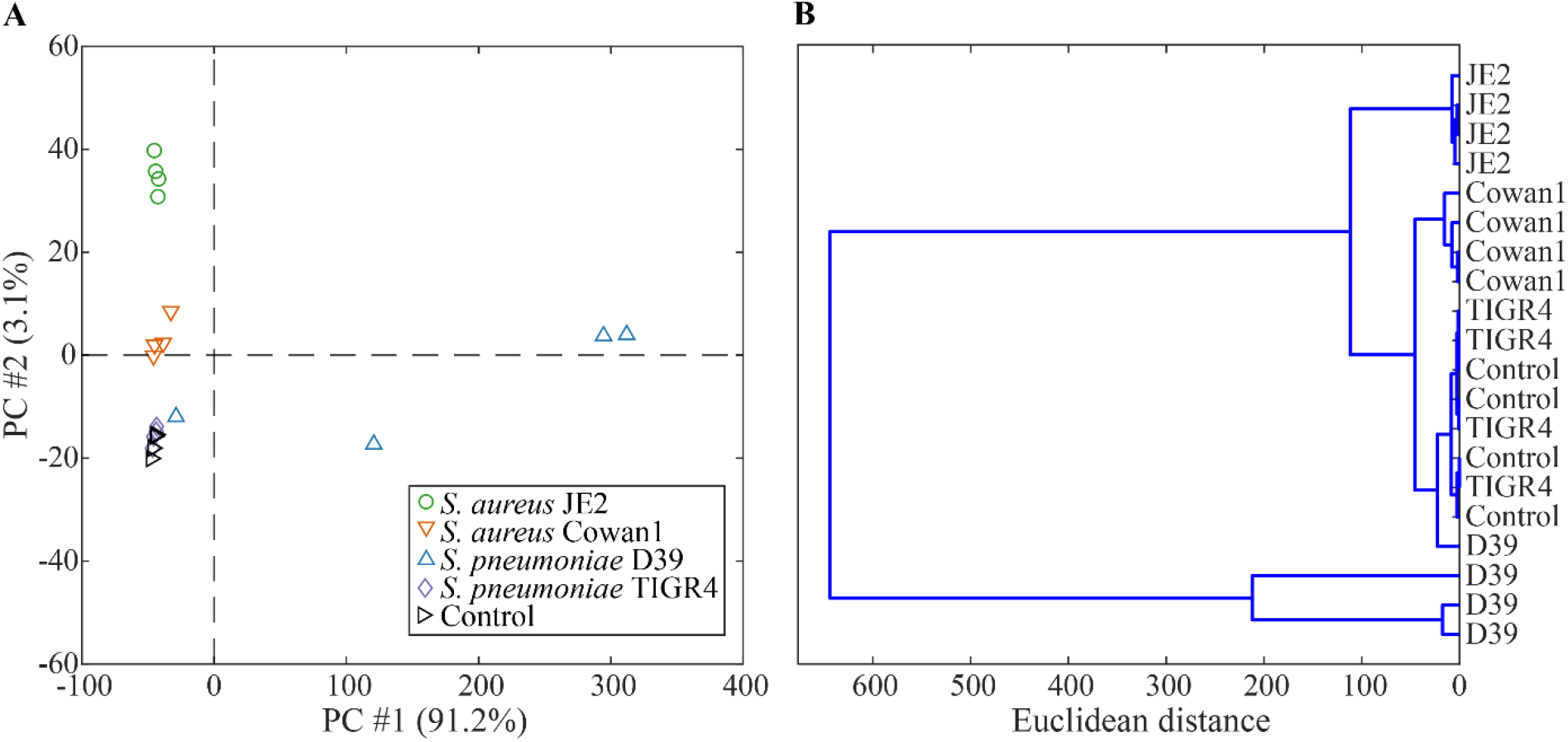
PCA score plot and dendrogram of PCA scores explaining 95% of variance illustrated at time point 12 h after the start of measurement. (A) PCA score plot of 1235 strain specific features (positive and negative ions) at 12 h identified for high CFU cultures. (B) Dendrogram showing the detailed hierarchical relationship between bacterial species and strains at time point 12 hours.

### Measurement of clinical patient samples by SESI-HRMS

After assessing the quantitative and qualitative capabilities of our real-time analysis system to detect bacterial growth in enriched bacterial cultures, we tested the feasibility of such an approach for the direct analysis of a heterogeneous set of 17 clinical samples from 13 different patients derived from various origins including heart valves, skin, deep tissue, as well as foreign bodies such as pacemakers (Table 1). All clinical samples were initially analyzed by routine diagnostics and the etiological agents identified by MALDI-TOF. Most samples came from patients which underwent antibiotic therapy prior to SESI-HRMS measurement. Hence, for 10 out of the 17 clinical samples, no bacterial growth was detected by conventional growth on agar plates. For the remaining seven samples for which bacterial growth was detected, four were *S. aureus* positive and three grew *S. epidermidis* at the time of measurement by SESI-HRMS (Table 1). This is a typical problem encountered in clinics rendering the current culture based microbiological diagnostics inefficient. Indeed, our clinical information confirmed that 11 out of 13 patients from this study were previously treated with different doses of antibiotics, and no bacterial growth could be detected at the sampling time for eight out of 13 of these patients (Table 1).

**Table 1.**
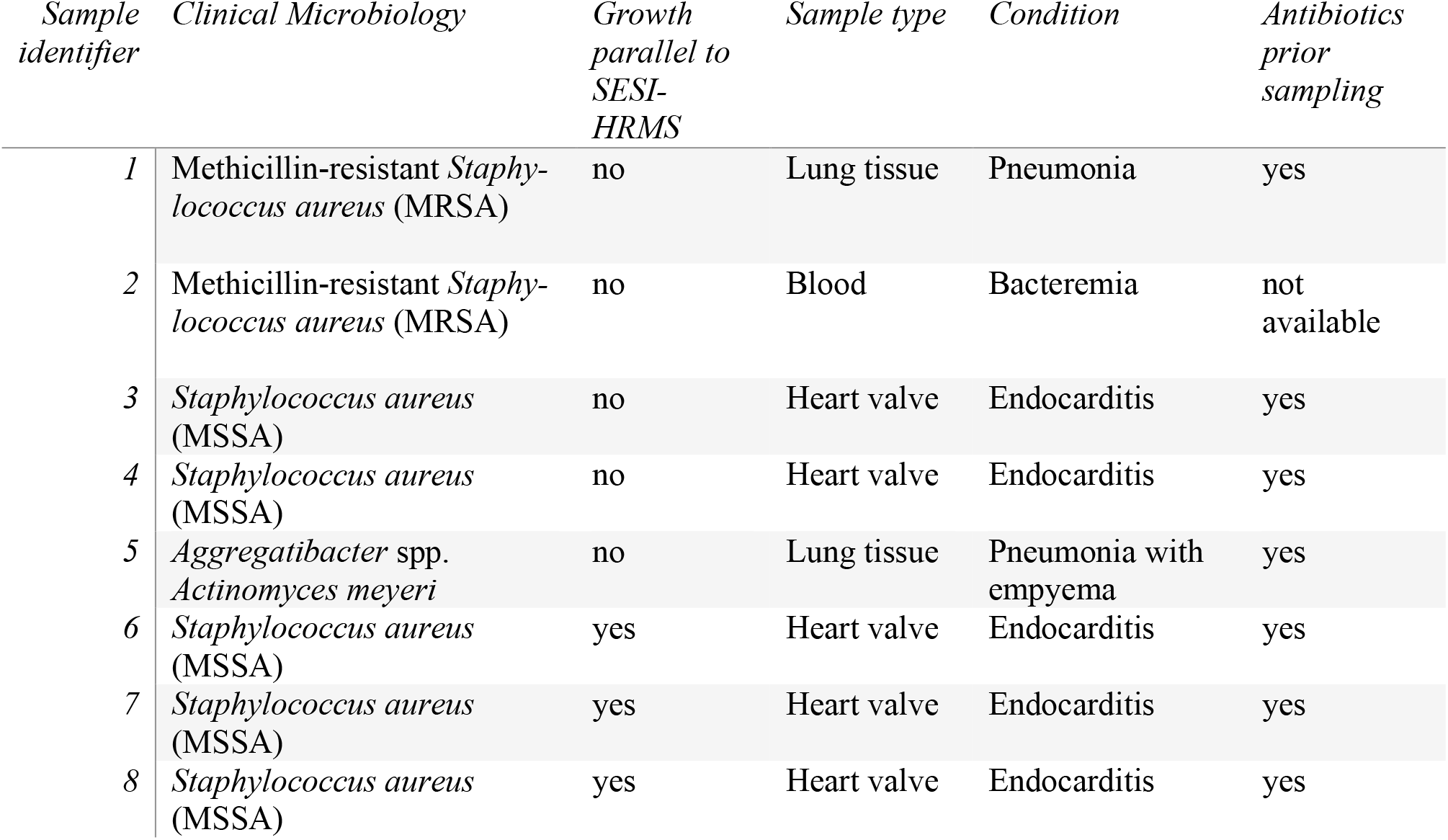
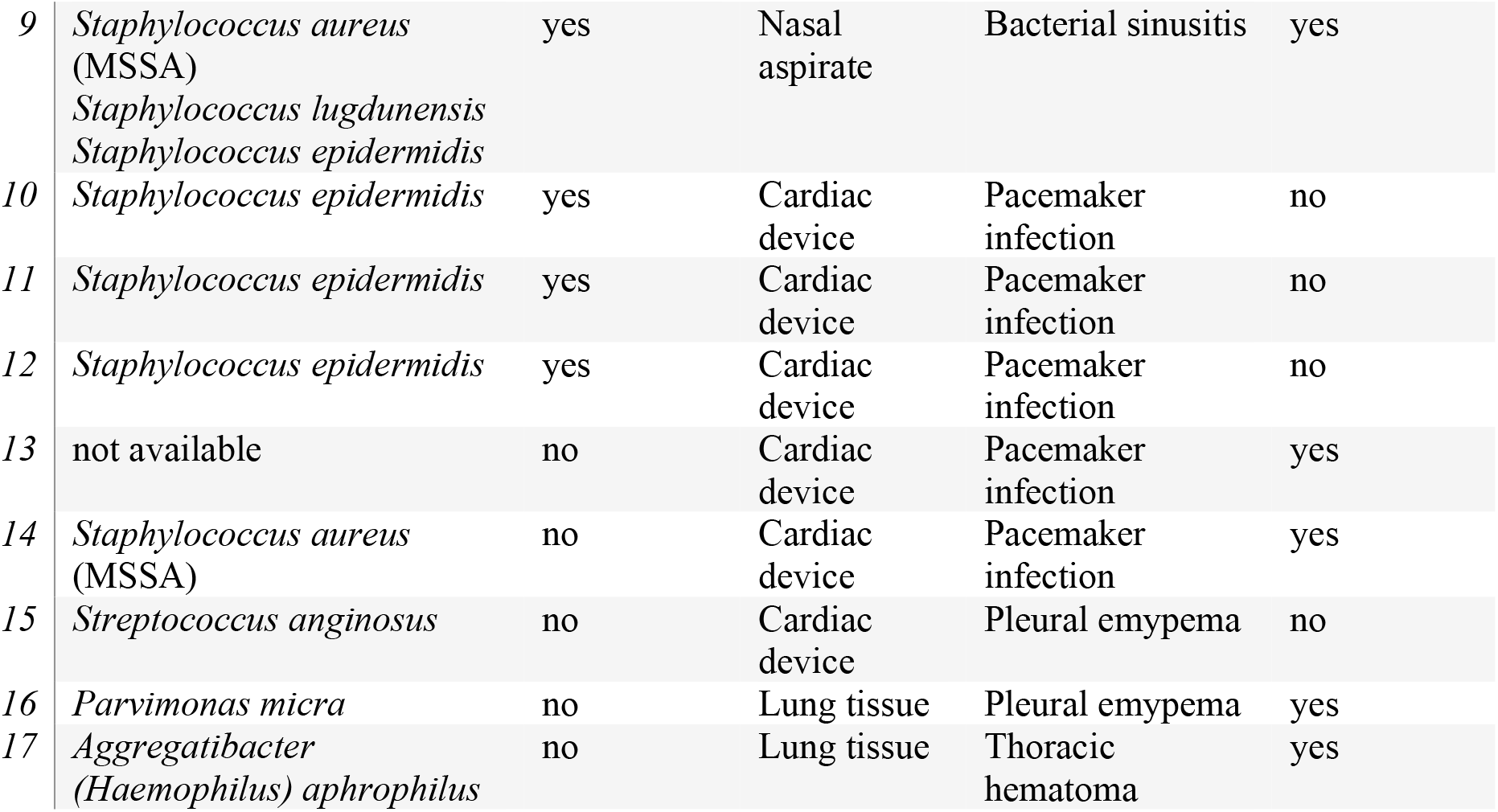
Clinical characteristics of patient samples analyzed by SESI-HRMS. Description of clinical microbiology, detected growth by SESI-HRMS, sample type, clinical condition and whether there was antibiotic treatment before analysis by SESI-HRMS.

Despite these challenges, all samples obtained from patients were subjected to a targeted analysis whereby the specific features previously identified under high-density conditions were extracted from the clinical dataset. To then visualize this highly complex dataset, t-SNE analysis was performed (Fig 4). A clear cluster in the middle of the t-SNE space consisting of clinical samples from *S. aureus* (methicillin-susceptible, MSSA) and *S. epidermidis* infections are visible. For these samples, growth of bacteria was confirmed *a posteriori* (Table 1). In addition, two clinical *S. aureus* samples (MRSA, further growth not confirmed) clustered at the bottom-right of the tSNE space.

**FIG 4.**
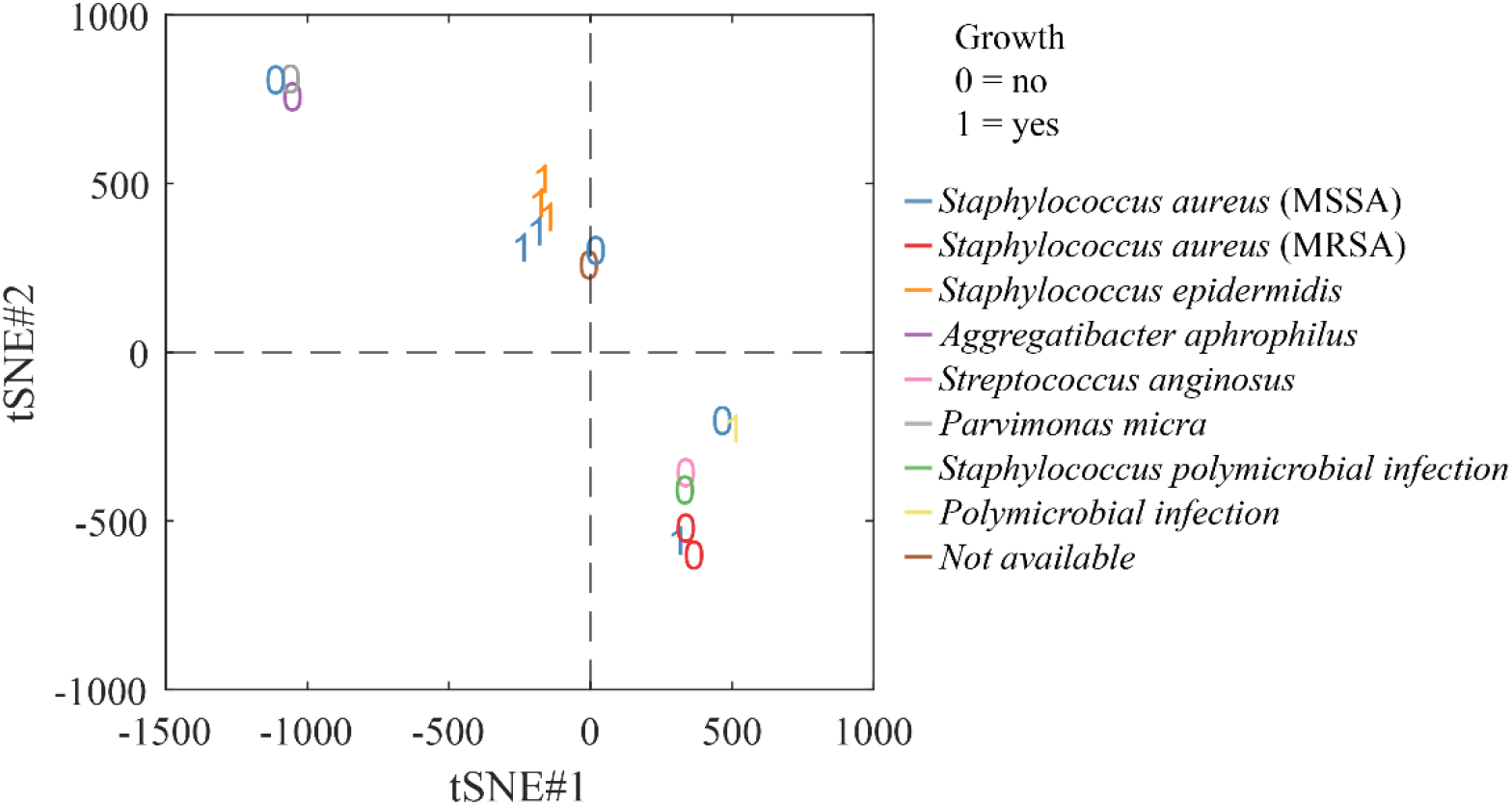
t-SNE analysis of clinical patient samples. Samples for which growth of the bacteria could be confirmed in parallel to SESI-HRMS measurement are represented as 1 = yes, whereas samples where no growth was observed are represented as 0 = no. The different colors indicate the causative bacterial strains responsible for the infection obtained in the sample withdrawn from patients. For details regarding the samples refer to table 1. MSSA = Methicillin-susceptible Staphylococcus aureus, MRSA = Methicillin-resistant Staphylococcus aureus.

## DISCUSSION

In this study we showed that SESI-HRMS detects in real-time unique features of the human pathogens *S. aureus* and *S. pneumoniae* within minutes of growth on an agar plate and of distinct strains. A high level of sensitivity for cultures with less than 1000 CFUs was achieved with detectable features, allowing for a clear differentiation between these important two human bacterial pathogens, even within strains. Since bacterial numbers are often low in patient samples, especially if the patients already is undergoing antimicrobial therapy, this is of great importance for future diagnostics.

For any diagnostic method to be of clinical use, it requires to be sensitive and specific enough to enable meaningful further clinical decisions such as an accurate antibiotic treatment. A third dimension of crucial importance in diagnostics of suspected bacterial infection is time-to-response. A perfect diagnostic method should be sensitive enough to detect a positive sample during early phases of infection, selective enough to distinguish different species or strains and should be fast and require little-to-no sample preparation. Currently, state-of-the-art DNA-based diagnostic methods for bacterial identification require just a single bacterial component to provide a positive response (18). However, limitations include that no differentiation between live and dead bacteria or a limited identification of antibiotic susceptibilities are possible. These limitations can seriously affect its clinical usefulness (10). On the other side, peptide profile identification by mass spectrometric methods can overcome these two noted limitations of DNA-based methods. However, this comes at the expense of requiring relatively lengthy pre-growing steps to enable active and sufficient bacterial cells to be detected by the MALDI-TOF system (17, 40, 41).

The proposed mass spectrometric method lies somewhere in between these two techniques, hence overcoming some of their limitations. On the one hand, detectable features accumulate in the headspace of the specimen only if the bacteria replicate, hence are alive. On the other hand, the data presented in here regarding the sensitivity of the SESI-HRMS suggests that bacterial loads in the order of 10^3^ CFUs are enough to be detected within one hour, well before sophisticated image analysis methods detect any indication of macroscopic bacterial growth which is the current gold standard in diagnostics. Also importantly, the proposed approach analyzes features in real-time, hence enables monitoring the blood agar plates directly as the bacteria grow. This provides a large automation potential as one can easily envisage multiplexing multiple dishes whereby an automatic valve would switch across samples to monitor their growth every few minutes.

Regarding the selectivity required to discriminate different pathogens, the high resolution of Orbitrap mass analyzers enables the separation of typically thousands of ions in the *m/z* range of 50-500, where most of the features from VOCs lie (42). Such resolving power renders a very high specificity potential when it comes to distinguish specific metabolic patterns stemming from different microorganisms. In our case, hundreds of features were found to be specific for each species. For the first time, we showed here that such level of specificity can go down to the strain level, reinforcing the notion that metabolomics is very well suited to capture such subtle heterogeneity as it provides a downstream read-out of genetic plus environmental factors (43, 44). The different environments between low- and high-CFU conditions may well explain the rather diverse metabolic signature observed under both conditions, leading to a modest overlap in the features detected under such different conditions.

Thus, overall SESI-HRMS identification of bacterial species by the present design suggests a high level of sensitivity and specificity, whereby already after 6 minutes, some strains, like *S. pneumoniae* D39, differ substantially from negative controls. The best separation was observed after 12 hours, which can be significantly quicker than the current MALDI-TOF identification procedures.

Proof-of-principle of the applicability of the method in a more realistic clinical context was also achieved by measuring clinical samples, including patient tissue and foreign material (e.g. pacemakers). This is one key advantage of this technique, as it requires no sample preparation, and therefore it is suitable for any solid or liquid specimen with a total analysis time of five-ten minutes to fingerprint the samples. In this study, 17 clinical samples from different origin were measured by SESI-HRMS. Following the standard procedure after a bacterial infection is diagnosed, those patients received antibiotics aiming to clear the infection. This often interferes with diagnostics because bacteria do not grow anymore after antibiotic challenge as was the case at the time of sampling and analysis by SESI-HRMS.

Four out of seven samples with an ongoing *S. aureus* infection and positive bacterial growth at the time of the analysis, clustered together in the t-SNE space (Fig 4). These results were obtained despite a low bacterial load in the clinical samples. While these results should be interpreted with caution, they clearly suggest that the proposed methodology for diagnostics in bacterial infections can be used with unprocessed clinical material. Similar to the peptide libraries used in MALDI-TOF bacterial analysis, further clinical work should be devoted to construct mass spectral libraries of VOCs, combined with modern classification algorithms to enable this technology and complement current state-of-the-art diagnostics.

This study also comes with limitations that need to be addressed, such as the use of only Grampositive bacteria. In addition, this study didn’t evaluate the identification of *Streptococcus mitis* or other difficult to identify bacteria. However, given the ability of this technique to identify unique features between two different strains of the same species, we believe this approach will allow for a more accurate determination of difficult-to-identify bacteria. Additionally, a machinelearning algorithm will need to be constructed using a much larger library of bacteria species and strains in a follow up study to validate our findings at a much larger scale including both Grampositive and Gram-negative bacteria of clinical interest.

## CONCLUSIONS

In this study, we tested the concept of exploiting the fact that bacteria produce complex volatile metabolic mixtures as they proliferate. SESI-HRMS features a high gas-phase species’ sensitivity, a great selectivity driven by the high-resolution of the mass analyzer. This was accomplished in real-time, without any sample preparation. These characteristics allowed for the first time to monitor the kinetic profiles of hundreds of metabolic species emitted by *S. aureus* and *S. pneumoniae* as they grew on agar plates. These hundreds of features rendered highly specific signatures, which enabled distinguishing the samples even at the strain level. Finally, we scaled-up the concept to test the feasibility of evaluating clinical samples retrieved from patients with bacterial infections. The results showed that such samples can be fingerprinted within five minutes. Characteristic metabolic patterns emerged, suggesting the potential of such an approach to complement current diagnostic methods. Further studies are required to construct VOC libraries of such specimens retrieved from patients to further test the clinical utility of this method.

## MATERIALS AND METHODS

### Bacterial Growth

*S. aureus* (JE2 and Cowan1) and *S. pneumoniae* (D39 and TIGR4) were initially cultivated axenically from glycerol stocks on Columbia agar plates with 5% sheep blood (BioMéreux) at 37°C and 5% CO_2_ for ∼15 to 16 hours (Fig 5A). Two different sets of experiments were performed with each strain. The first experiment consisted of plating a high-density culture (i.e. high CFUs) by performing a subculture on a fresh blood agar plate directly from the overnight plate. In the second experiment, the initial overnight culture in agar was resuspended in PBS and diluted to obtain a low number of CFUs (i.e. low CFUs) ranging from ∼140 to ∼2000 CFUs per plate.

**FIG 5.**
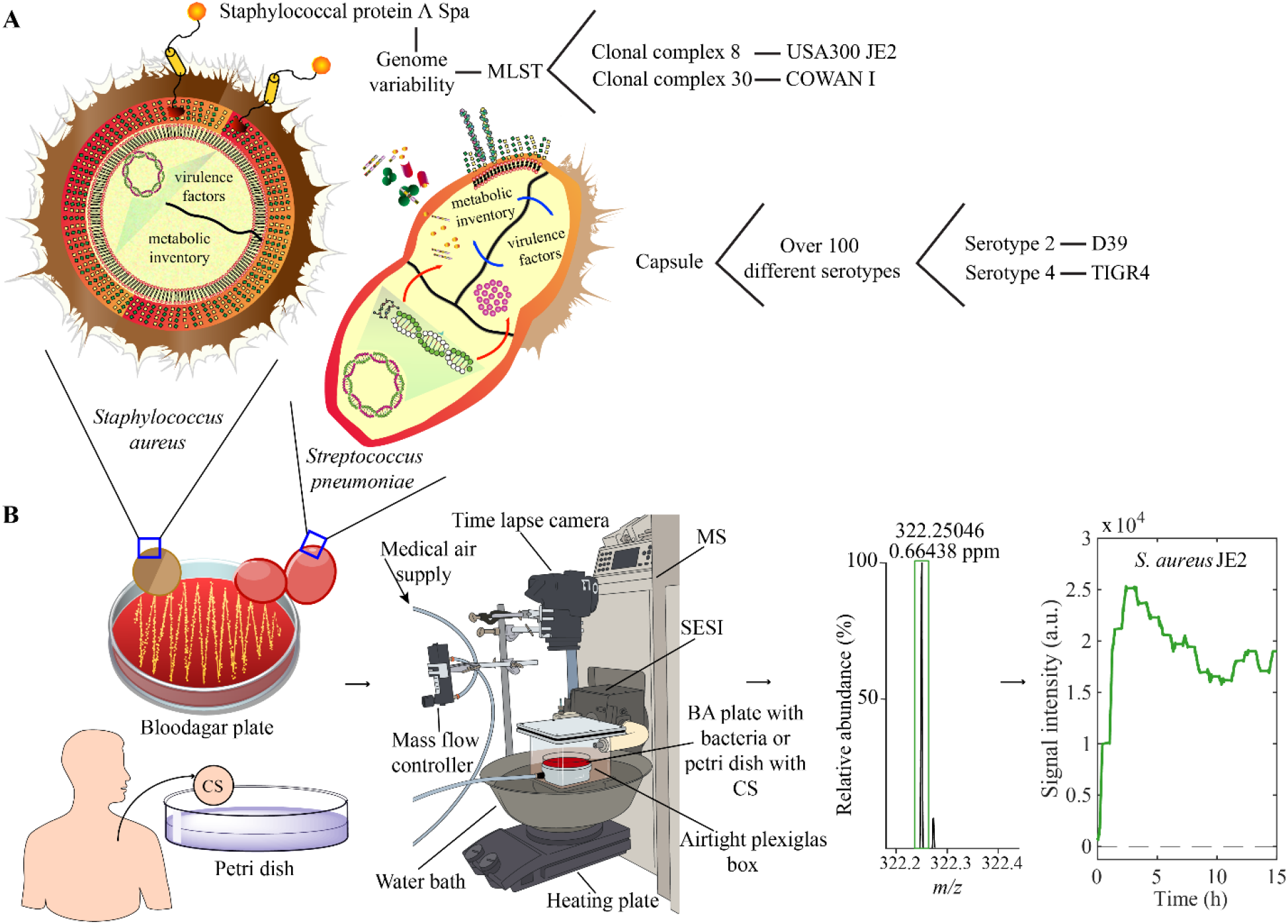
Experimental setup designed for the detection of features in the headspace of bacterial cultures by SESI-HRMS. (A) Genomic difference between the two species *S. aureus* and *S. pneumoniae* and their respective strains. (B) From left to right, either a strain of *S. aureus* (JE2 or Cowan1) or a strain of *S. pneumoniae* (D39 or TIGR4) on a blood agar plate or a clinical sample on a petri dish was placed in an airtight plexiglass box coupled to the SESI-HRMS. A TL camera was directly placed above the box for visualization of bacterial growth in parallel to the SESIHRMS measurement. The setup ensured a closed environment where the features present in the headspace of the samples were guided through a medical grade air flow of 0.5 L/min towards the SESI where they were ionized and then separated according to their mass to charge ratio (*m/z*) in the MS, resulting in real-time traces over ∼15 h (bacterial plates) and five minutes (clinical samples). BA = Blood agar, CS = Clinical sample, MS = Mass spectrometer, SESI = Secondary Electro-Spray Ionization.

### Sampling of Clinical Patient Material

Clinical samples were obtained from patients with bacterial infections requiring surgery. The description of the different clinical samples used in this study such as origin, identified bacterial pathogen, and antibiotic treatment prior to sampling is depicted in table 1. The processing of the patient material depended on the characteristics of the sample. To verify the growth of bacteria from these samples, the original sample was divided into two parts, one for measurement by SE-SI-HRMS (Fig 5B) and one for colony plating and quantification. In short, samples such as skin, heart valves and soft tissues were processed by disrupting the tissue using a tissue lyser (Qiagen TissueLyzer, vibration frequency of 30/s for 10 minutes). For foreign material such as pacemakers, the processing involved an initial sonication step of 5 minutes using a sonicator bath (Ultrasonic bath XUBA3, Grant) in sterile PBS. Following these two initial steps, the remaining processing protocol was the same for all samples. The resulting suspension was washed twice with sterile PBS to remove traces of antibiotics from the sample and a final step with sterile milliQ water was used to lyse the eukaryotic cells. The sample was then serial diluted in sterile milliQ water, plated on blood agar plate and incubated at 37 °C. In total, 17 clinical samples were used for analysis.

### Sample Measurement with SESI-HRMS

The experimental set-up consisted of a custom-made plexiglass box with an airtight closing mechanism which was directly connected to an ion source (Super SESI, FIT, Spain) coupled to a high-resolution mass spectrometer (Exactive Plus, Thermo Fisher Scientific, Germany) (Fig 5B). The sample (bacteria plate or clinical patient material) was placed inside the plexiglass box which was heated at 37 °C in a water bath. A mass flow controller was coupled to the box on the opposite side of the SESI-HRMS via PTFE - tubes and ensured a constant medical grade air supply through the system at a flow rate of 0.5 L/min and carried the VOCs emitted by the bacterial cultures or from the clinical material towards the SESI-HRMS. Mass spectral analysis of bacteria plates was conducted over a period of ∼15 hours and for 5 min in case of clinical samples. An automated switch system (Auto Click Typer version 2.0) allowed to alter between positive and negative ionization mode every 30 minutes when measurements were conducted for ∼15h. In addition, a high-resolution camera was placed above the box and was triggered as described in the “bacterial plate imaging” section of the methods. In total, 36 measurements were performed whereby for both conditions (high-density and low CFUs) a total of 16 measurements were conducted each (n=4 biological replicates per strain) along with measurements of empty blood agar plates which served as control measures (n=4) for both CFU conditions.

To generate the electrospray in the SESI, a 20-µm ID TaperTip silica capillary emitter (New Objective, USA) and a solution of 0.1% formic acid in water were used. The pressure of the SESI solvent environment was set to 1.3 bar. Temperature of the ionization chamber and the sampling line was set to 90 °C and 130 °C respectively. The voltage of the electrospray was set to 3.7 KV in positive and 3 KV in negative ionization mode. The sheath gas flow rate was set to 10, capillary temperature was 320 °C and S-lens RF level 50.0.

Mass spectra were acquired via Thermo Exactive Plus Tune software (version 2.9) in full scan mode (scan range 50 – 500 *m/z*, polarity positive or negative, microscan number 10, ACG target 10^6^, maximum injection time 50 ms) at a resolving power of 140000 at *m/z* 200. The system was calibrated on a regular basis before the measurements externally and internally by using common background contaminants as lock masses in the respective polarity (45, 46).

### Bacterial Plate Imaging

Simultaneously as the plate was analyzed for the production of VOCs by SESI-HRMS, a Time lapse (TL) imaging experiment was performed using a high-resolution camera (Cannon EOS 1200D reflex) triggered every 10 minutes by an Arduino Uno board (Arduino) to capture images of the plate inside the box to visually document bacterial growth (47). To verify the growth of the bacteria on the plate, TL measurement was conducted until ∼ 40 h for specific replicates.

### SESI-HRMS Data Analysis

Data analysis was performed using MATLAB (version 2021b, MathWorks Inc., USA). Raw mass spectra files were accessed via inhouse C# console apps based on Thermo Fisher Scientific’s RawFileReader (version 5.0.0.38). MATLAB functions, *mspeaks* and *ksdensity* were applied to extract the final list of features. As a result, a data matrix of total 571 × 3460 (files x mass spectral features) in positive mode and x 1129 mass spectral features in negative mode was obtained. Specific m/z peaks had to be excluded due noisy interferences from the mass spectrometer. Signal intensity time traces of all features were then computed and smoothed (moving mean; span = 300) for visualization purposes. The mean Area Under the Curve (mAUC) of the time traces (n=4 replicates per strain) was calculated by interpolating the data every 0.01h. To identify features, unique to a particular bacterial strain, two criteria were defined; first criteria kept only features with a log2 fold change (FC) ≥ 2 in mAUC of a particular strain compared to the mAUC of the control and 2^nd^ criteria required a log2 FC≥ 4 in mAUC of the particular strain compared to the averaged mAUC of the other investigated strains. Furthermore, species specific time traces were identified, meaning that they had to be present in both *S. pneumoniae* strains (D39 and TIGR4) or both *S. aureus* strains (JE2 and Cowan1). Therefore three criteria were defined: first criteria kept only features with a log2 FC ≥ 2 in mAUC in both strains of a particular species compared to the mAUC of the control; 2^nd^ criteria defined to only consider the features further if within one species the respective strains had at least a mAUC of 30% of the mAUC of the other strain under the respective species to avoid features to be selected which tended to be rather present in one strain and not in both; 3^rd^ criteria considered only features which showed a log2 FC≥ 4 in average mAUC of the two strains of one species compared to the average mAUC of the two strains of the other species and vice versa. In a next step the time traces of features unique for the different strains and species were auto scaled (z-score), subjected to a hierarchical cluster tree (Ward method; Euclidean distance) and visualized as heat maps showing the evolution of the features over time. Principal Component Analysis (PCA) of 5^th^ – root transformed data matrix and a hierarchical binary cluster tree (Ward Method; Euclidean distance) were used to visually discriminate the different bacterial strains at distinctive time points over ∼15 hours. Clinical samples were analyzed using a targeted approach, where unique positive and negative time traces of features previously identified for the different strains under high density CFU condition were directly extracted from the clinical samples raw data using in-house C# console app based on RawFileReader which resulted in a data matrix of 17 × 1269 (samples x mass spectral features). We then performed t-distributed stochastic neighbor embedding (t-SNE) to visualize this highly complex and exploratory dataset. For all features, molecular formulae were assigned based on accurate mass by using the “seven golden rules” (48), considering the elements C, H, N, O, P and S and the adducts [M + H], [M - H_2_O + H], [M + NH_4_], [M - NH_3_ + H], [M + Na] in positive mode and [M - H], [M – Na], [M - H_2_O - H], [M + NH_3_ - H], [M - NH_4_] in negative ionization mode.

### Image Data Analysis

Simultaneously to the SESI-HRMS measurement, bacterial growth on agar was verified and quantified by TL imaging. The acquired TL images were analyzed with a custom extension for ColTapp (47) to quantify the bacterial growth as changes in pixel intensity over time. As the visual growth pattern of *S. aureus* and *S. pneumoniae* are distinct, different image analysis pipelines were utilized. For *S. aureus*, images were transformed to grayscale by selecting the green channel of the RGB image and subsequent top-hat filtering was performed to reduce lighting heterogeneity. For *S. pneumoniae*, the RGB images were transformed to YIQ color space and subset to the I channel only. A gaussian filter was applied to the first image of the TL series to define a background which then was subtracted from each other image in the TL series.

The following steps were the same for both species: the corrected grayscale images were subset to include only the area within plate boundaries. Then, min-max scaling with [0, 0.7] as range was applied to the pixel intensities. Finally, the sum of all pixel intensities was divided by the sum of pixels to derive a normalized intensity value per time point of a TL image series.

## Ethical Statement

For this study, samples from patients with vascular graft /endovascular infections, infective endocarditis, bone and prosthetic joint infections and any other infections were collected under the framework of the Vascular Graft Cohort study (VASGRA; KEK-2012-0583), the Endovascular and Cardiac Valve Infection Registry (ENVALVE; BASEC 2017-01140), the Prosthetic Joint Infection Cohort (Balgrist, BASEC 2017-01458), and BacVivo (BASEC 2017-02225), respectively. The study was approved by the local ethics committee of the Canton of Zurich, Switzerland

## Acknowledgements

We thank Prof. Dr. Sven Hammerschmidt from the Center for Functional Genomics of Microbes in Greifswald, Germany, for providing both *Streptococcus pneumoniae* strains D39 and TIGR4 (49, 50). We acknowledge the work from the medical personal from the University Hospital Zurich which facilitated the acquisition of the clinical samples and thank the patients for participating.

This work was supported by a grant from Foundation Botnar (Switzerland) and the Swiss National Science Foundation No. 320030_173168 and PCEGP3_181300 to PS, the Swiss National Science Foundation grants nbr. 31003A_176252 to ASZ, as well as by the Clinical Research Priority Program of the University of Zurich Precision Medicine for Bacterial Infections to ASZ and SDB. This work is part of the Zurich Exhalomics project under the umbrella of University Medicine Zurich/Hochschulmedizin Zürich.

## Author Contributions

Conception and Design, AGM, KA, PS and ASZ; Clinical Sample Collection and Processing, AGM, TCS, SDB and ASZ. Data Analysis and interpretation, AGM, KA, JB, KDS, PS and ASZ. Manuscript Writing – Original Draft, AGM, KA, PS and ASZ. Writing, Review &Editing, AGM, KA, JB, KDS, TCS, SDB, PS and ASZ. All authors read and approved the final manuscript.

## Conflicts of interest

PS is cofounder of Deep Breath Initiative A.G. (Switzerland), which develops breath-based diagnostic tools. KDS is consultant for Deep Breath Initiative A.G. (Switzerland).

## FIGURE LEGENDS SUPPLEMENTAL MATERIAL

**FIG S1** TL of bacterial strains and blood agar controls under low and high CFU conditions. (A) TL normalized pixel intensity of the control replicates. The control measures between the two bacterial species (*S. aureus* and *S. pneumoniae*) looked different since a different analysis protocol was used for the evaluation. (B) TL normalized pixel intensity illustrated for the two *S. pneumoniae* strains. (C) TL normalized pixel intensity illustrated for the two *S. aureus* strains. The color legend represents the biological replicates (n=4) measured for each bacterial strain and control along with the number of CFUs used under defined conditions for each biological replicate. TL = Time lapse, CFU = Colony forming unit.

**FIG S2** Specific time – dependent features detected during bacterial growth. (A) Example time trace of the positive ion at *m/z* 201.0433 unique to *S. pneumoniae* D39 is shown on top of the heatmaps consisting of total 1178 features (positive ions) specific to *S. pneumoniae* D39. (B) Example time trace of the negative ion at *m/z* 89.9913 unique to *S. pneumoniae* D39 is shown on top of the heatmaps consisting of total 28 features (negative ions) specific to *S. pneumoniae* D39. (C) Example time trace of the negative ion at *m/z* 209.1032 unique to *S. aureus* JE2 is shown on top of the heatmaps consisting of total 16 features (negative ions) specific to *S. aureus* JE2. (D) Example time trace of the positive ion at *m/z* 135.121 unique to *S. aureus* (JE2 and Cowan1) is shown on top of the heatmaps consisting of total five *m/z-*features (positive ions) specific to *S. aureus* (JE2 and Cowan1). (E) Example time trace of the negative ion at *m/z* 82.0298 unique to *S. pneumoniae* (D39 and TIGR4) is shown on top of the heatmaps consisting of total 14 features (negative ions) specific to *S. pneumoniae*. (F) Time trace of the only positive ion at *m/z* 235.0757 unique to *S. pneumoniae*. (G) Example time trace of the positive ion at *m/z* 179.1042 unique to *S. aureus* Cowan1. Real-time evolution of all features is shown over 15 hours of measurement by SESI-HRMS for all four investigated strains (n=4 biological replicates) and controls (n=4). The color bar indicates the z-score values of absolute MS signal intensity for each feature, from low (dark blue) to increased signal intensity (dark red).

**FIG S3** Overlapping features unique to *S. pneumoniae* (i.e. present in both D39 and TIGR4) under low CFU and high CFU condition. (A) Venn diagram of overlapping features detected for low and high CFU bacterial cultures. (B) Representative time trace of the positive ion at *m/z* 385.2214 observed in *S. pneumoniae* (D39 and TIGR4) for low and high-density cultures. CFU = Colony forming unit.

**FIG S4** PCA score plots over time obtained for high density bacterial cultures. Score plots of 1190 strain specific features (positive ions) for time points 0.1 hour and 0.25 hour. Score plots of 1235 strain specific features (positive and negative ions) at distinct time points over the 15 hours measurement period.

**FIG S5** Dendrogram trees over time obtained for high density bacterial cultures. Cluster analysis of samples considering 1190 strain specific features (positive ions) for time points 0.1 hour and 0.25 hour. Cluster analysis of samples using 1235 strain specific features (positive and negative ions) at distinct time points during the 15 hours measurement period.

## TABLE DESCRIPTIONS SUPPLEMENTAL MATERIAL

**Table S1** Unique features assigned to *S. aureus* and *S. pneumoniae* in positive and negative ionization mode under low CFU condition.

**Table S2** Unique features assigned to the different strains of *S. aureus* and *S. pneumoniae* in positive and negative ionization mode under high CFU condition.

