## Supplemental figures and tables for "Rapid detection of *Staphylococcus aureus* and *Streptococcus pneumoniae* by real-time analysis of volatile metabolites"

**Running title: Early detection of bacteria by SESI-HRMS**

[#]

[#]

### FIGURES

#### FIG S1

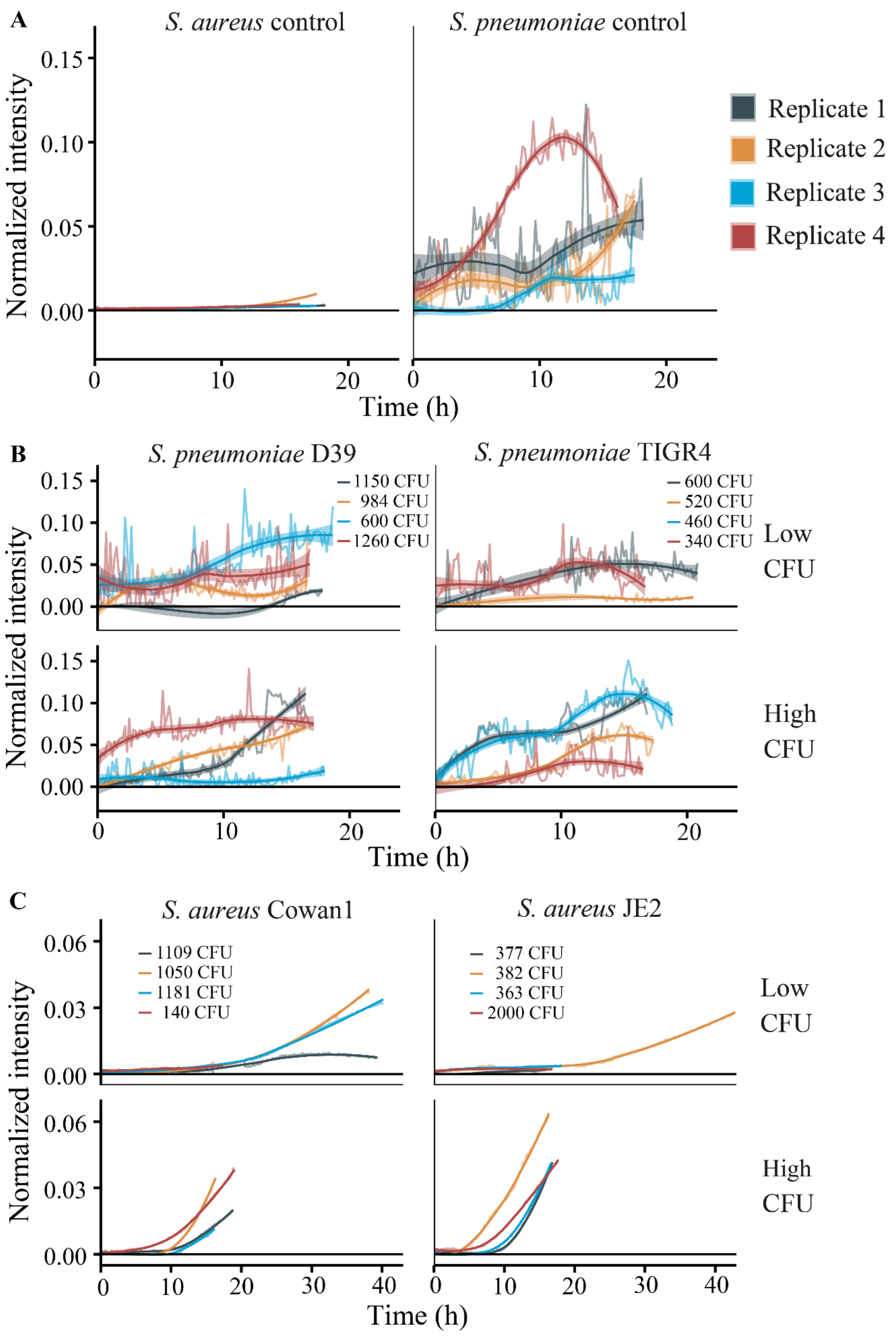

**FIG S2**

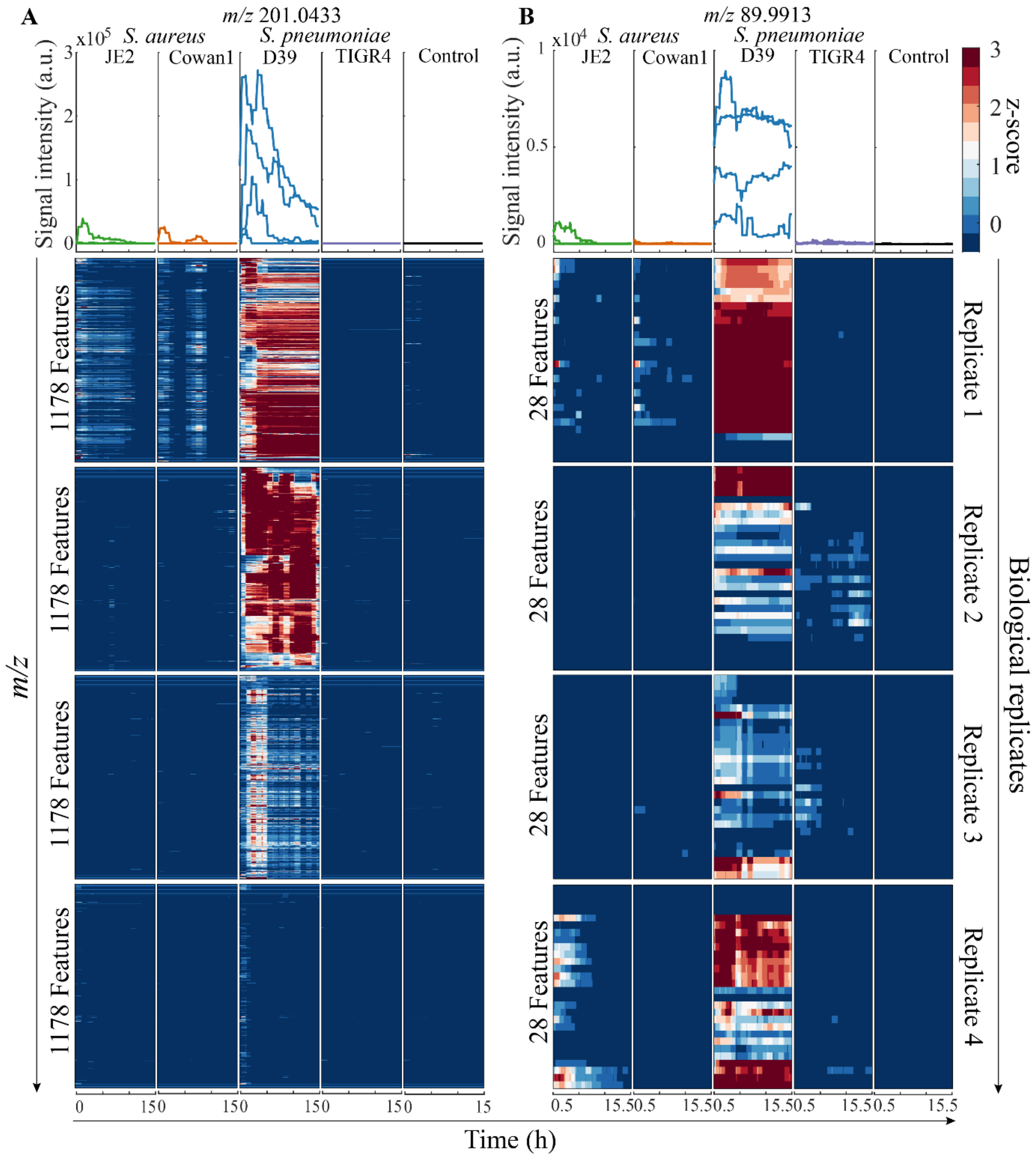
A
m/z 201.0433
x10^5
S. aureus
S. pneumoniae
JE2
Cowan1
D39
TIGR4
Control
Signal intensity (a.u.)
1178 Features
m/z
B
m/z 89.9913
x10^4
28 Features
Replicate 1
Replicate 2
Replicate 3
Replicate 4
Biological replicates
z-score
Time (h)

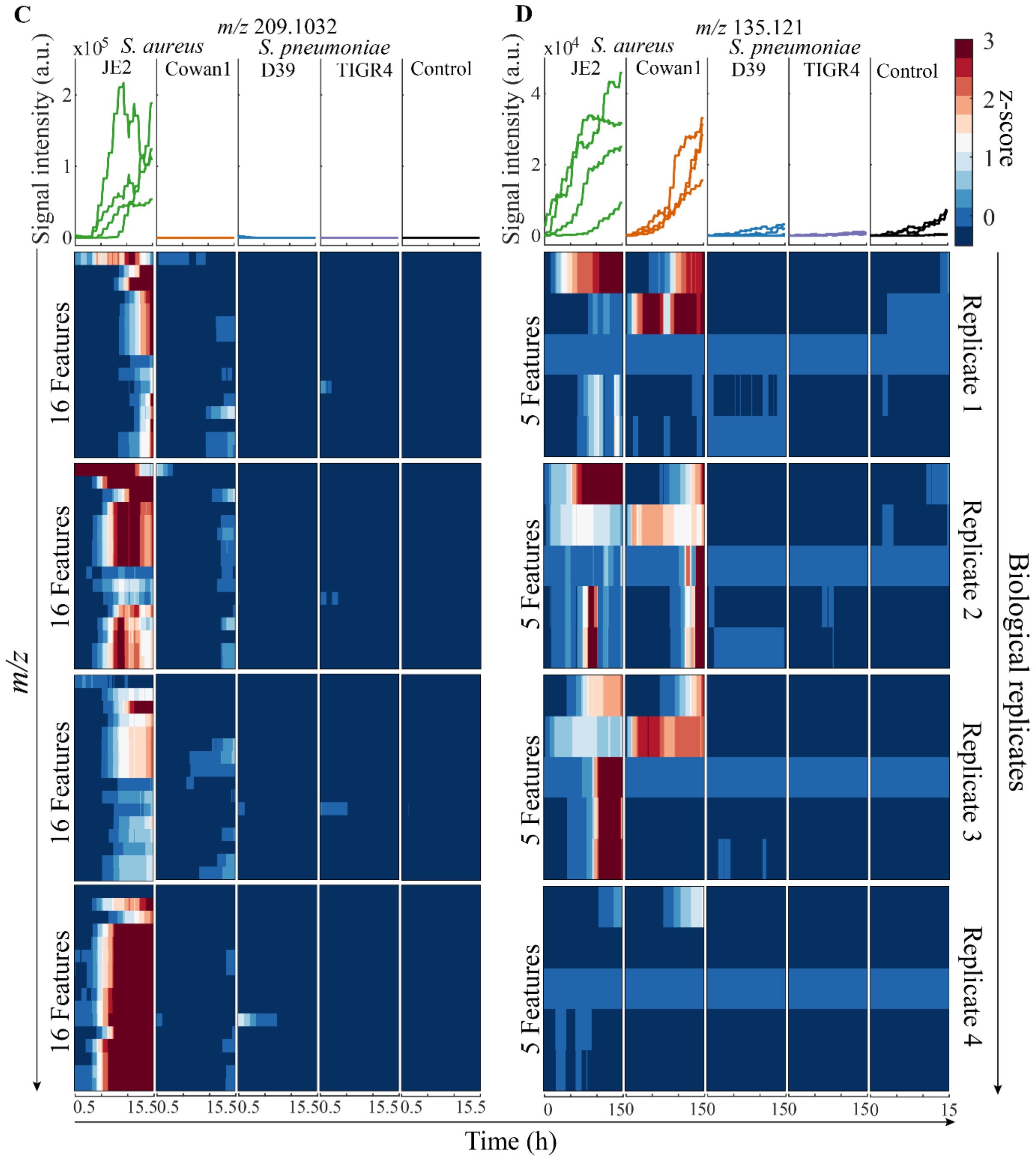
C
m/z 209.1032
x10^5
S. aureus
S. pneumoniae
JE2
Cowan1
D39
TIGR4
Control
Signal intensity (a.u.)
16 Features
m/z
0.5 15.5
D
m/z 135.121
x10^4
5 Features
Replicate 1
Replicate 2
Replicate 3
Replicate 4
Biological replicates
z-score
Time (h)

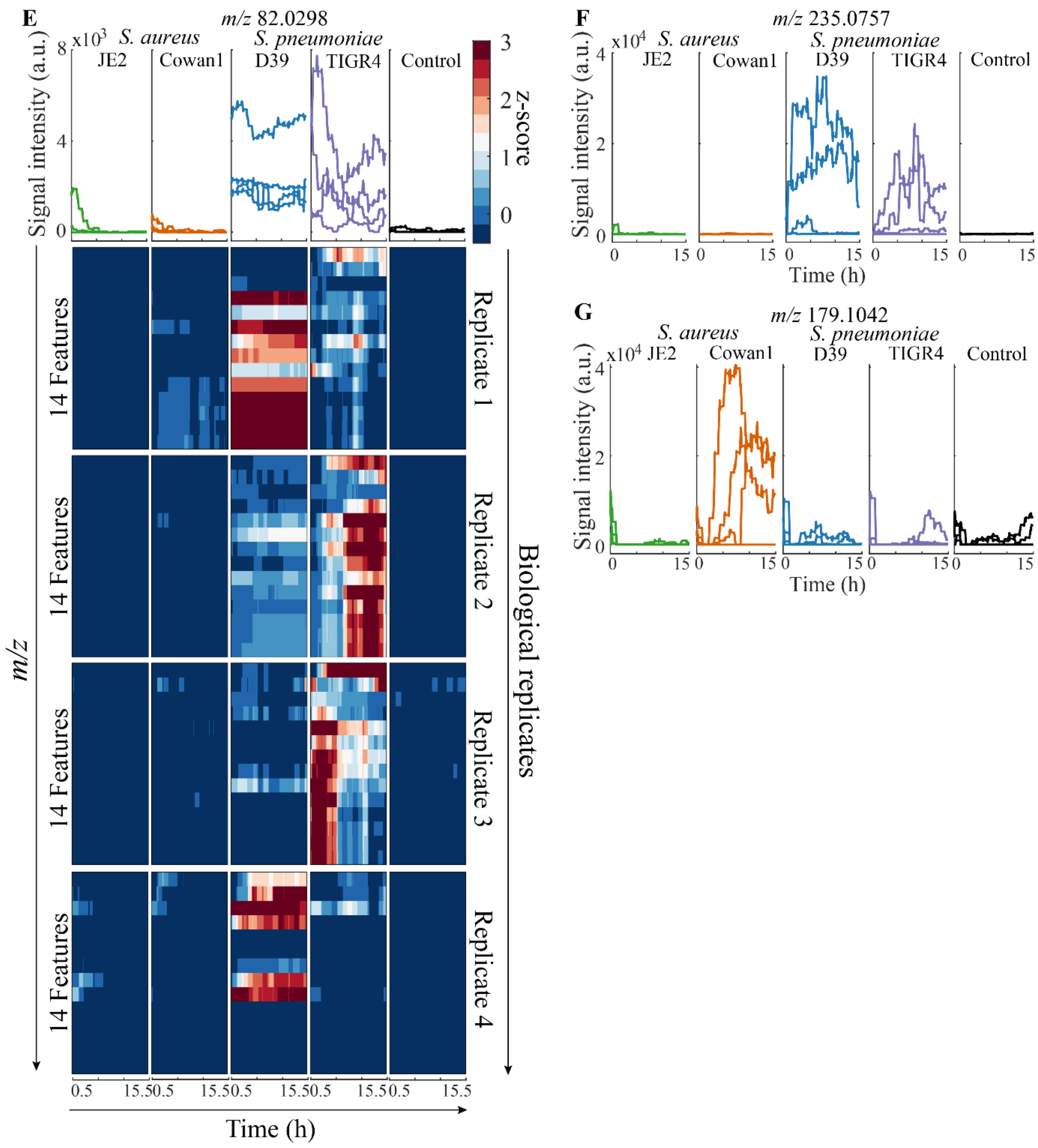
E
m/z 82.0298
x10^3
S. aureus
S. pneumoniae
JE2
Cowan1
D39
TIGR4
Control
Signal intensity (a.u.)
z-score
14 Features
Replicate 1
Replicate 2
Replicate 3
Replicate 4
Biological replicates
m/z
Time (h)
F
m/z 235.0757
x10^4
G
m/z 179.1042

**FIG S3**

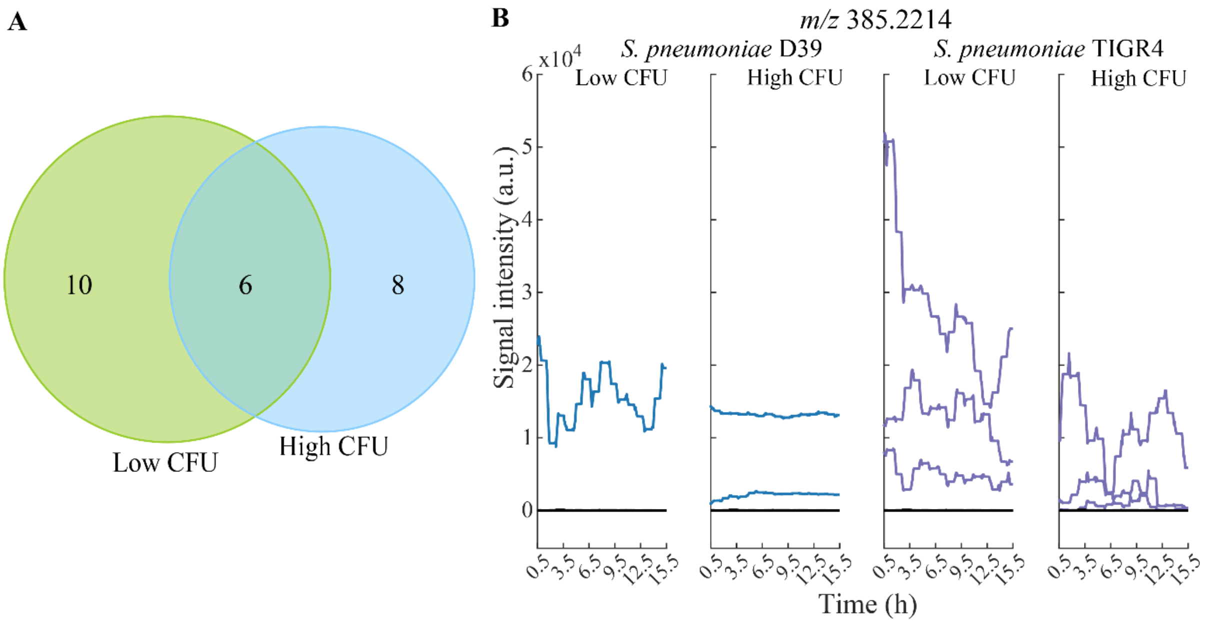
A
10
6
8
Low CFU
High CFU
B
m/z 385.2214
S. pneumoniae D39
S. pneumoniae TIGR4
x10^4
Low CFU
High CFU
Low CFU
High CFU
Signal intensity (a.u.)
0
1
2
3
4
5
6
0.5
3.5
6.5
9.5
12.5
15.5
Time (h)

### FIG S4

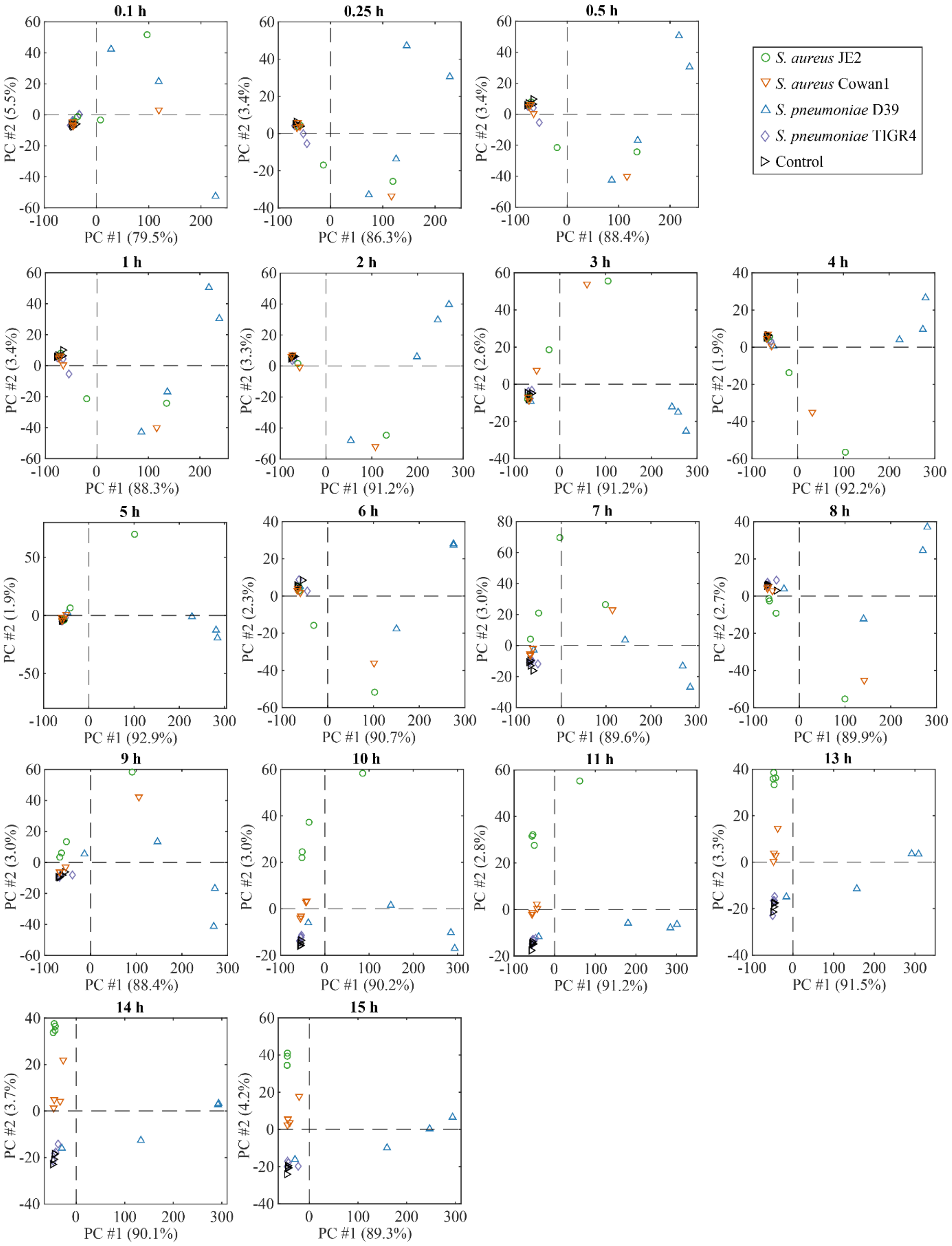
0.1 h
PC #1 (79.5%)
PC #2 (5.5%)
0.25 h
PC #1 (86.3%)
PC #2 (3.4%)
0.5 h
PC #1 (88.4%)
PC #2 (3.4%)
S. aureus JE2
S. aureus Cowan1
S. pneumoniae D39
S. pneumoniae TIGR4
Control
1 h
PC #1 (88.3%)
PC #2 (3.4%)
2 h
PC #1 (91.2%)
PC #2 (3.3%)
3 h
PC #1 (91.2%)
PC #2 (2.6%)
4 h
PC #1 (92.2%)
PC #2 (1.9%)
5 h
PC #1 (92.9%)
PC #2 (1.9%)
6 h
PC #1 (90.7%)
PC #2 (2.3%)
7 h
PC #1 (89.6%)
PC #2 (3.0%)
8 h
PC #1 (89.9%)
PC #2 (2.7%)
9 h
PC #1 (88.4%)
PC #2 (3.0%)
10 h
PC #1 (90.2%)
PC #2 (3.0%)
11 h
PC #1 (91.2%)
PC #2 (2.8%)
13 h
PC #1 (91.5%)
PC #2 (3.3%)
14 h
PC #1 (90.1%)
PC #2 (3.7%)
15 h
PC #1 (89.3%)
PC #2 (4.2%)

### FIG S5

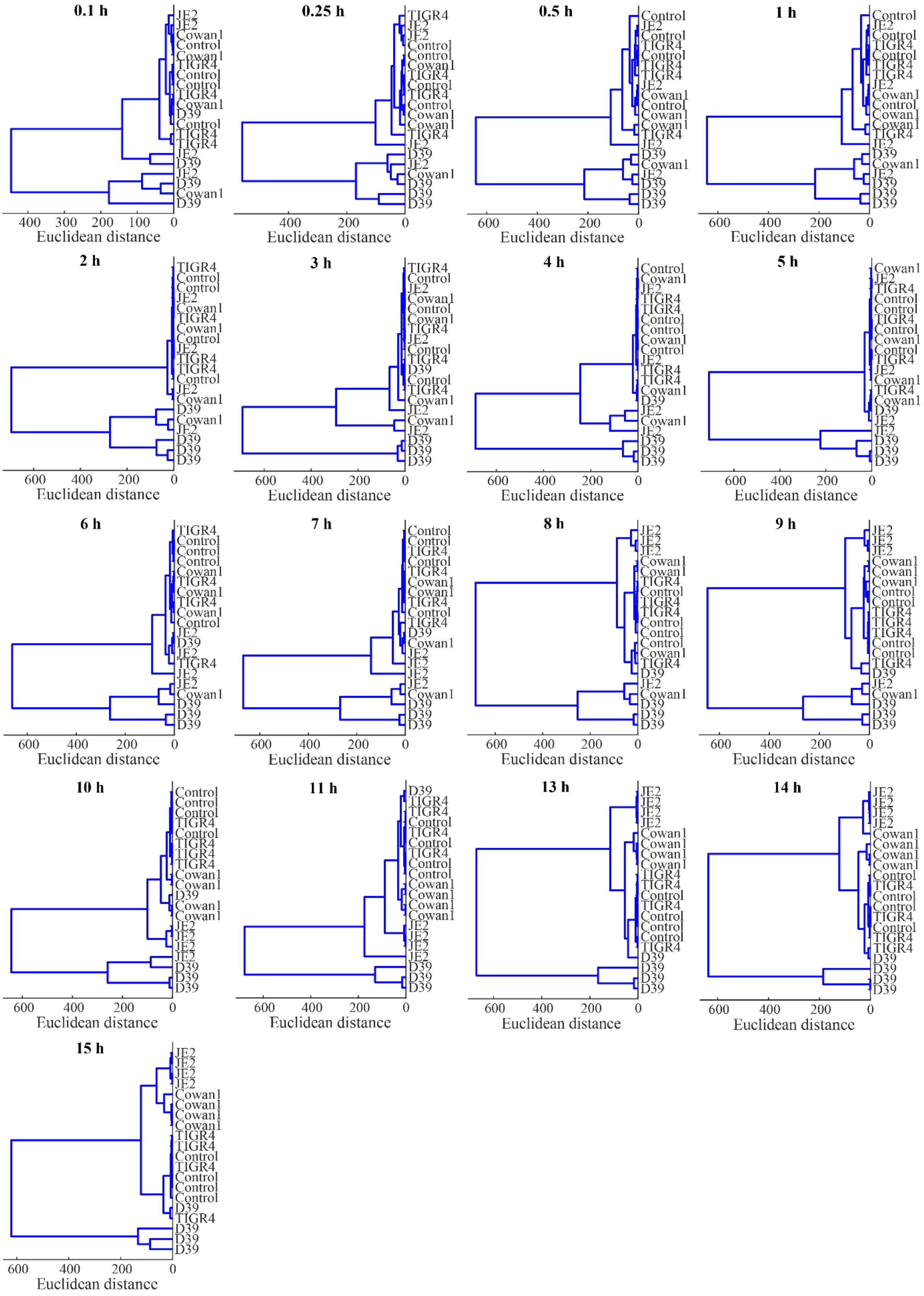
0.1 h
JE2
JE2
Cowan1
Control
Cowan1
TIGR4
Control
Control
TIGR4
Cowan1
D39
Control
TIGR4
TIGR4
JE2
D39
JE2
D39
Cowan1
D39
400 300 200 100 0
Euclidean distance
0.25 h
TIGR4
JE2
JE2
Control
Control
Cowan1
TIGR4
Control
TIGR4
Control
Cowan1
Cowan1
TIGR4
JE2
D39
JE2
Cowan1
D39
D39
D39
400 200 0
Euclidean distance
0.5 h
Control
JE2
Control
TIGR4
Control
TIGR4
TIGR4
JE2
Cowan1
Control
Cowan1
Cowan1
TIGR4
JE2
D39
Cowan1
JE2
D39
D39
D39
600 400 200 0
Euclidean distance
1 h
Control
JE2
Control
TIGR4
Control
TIGR4
TIGR4
JE2
Cowan1
Control
Cowan1
Cowan1
TIGR4
JE2
D39
Cowan1
JE2
D39
D39
D39
600 400 200 0
Euclidean distance
2 h
TIGR4
Control
Control
JE2
Cowan1
TIGR4
Cowan1
Control
JE2
TIGR4
TIGR4
Control
JE2
Cowan1
D39
Cowan1
JE2
D39
D39
D39
600 400 200 0
Euclidean distance
3 h
TIGR4
Control
JE2
Cowan1
Control
Cowan1
TIGR4
JE2
Control
TIGR4
D39
Control
TIGR4
Cowan1
JE2
Cowan1
JE2
D39
D39
D39
600 400 200 0
Euclidean distance
4 h
Control
Cowan1
JE2
TIGR4
TIGR4
Control
Control
Cowan1
Control
JE2
TIGR4
TIGR4
Cowan1
D39
JE2
Cowan1
JE2
D39
D39
D39
600 400 200 0
Euclidean distance
5 h
Cowan1
JE2
TIGR4
Control
Control
TIGR4
Control
Cowan1
Control
TIGR4
JE2
Cowan1
TIGR4
Cowan1
D39
JE2
JE2
D39
D39
D39
600 400 200 0
Euclidean distance
6 h
TIGR4
Control
Control
Control
Cowan1
TIGR4
Cowan1
TIGR4
Cowan1
Control
JE2
D39
JE2
TIGR4
JE2
JE2
Cowan1
D39
D39
D39
600 400 200 0
Euclidean distance
7 h
Control
Control
TIGR4
Control
TIGR4
Cowan1
Cowan1
TIGR4
Control
TIGR4
D39
Cowan1
JE2
JE2
JE2
JE2
Cowan1
D39
D39
D39
600 400 200 0
Euclidean distance
8 h
JE2
JE2
JE2
Cowan1
Cowan1
TIGR4
Control
TIGR4
TIGR4
Control
Control
Control
Cowan1
TIGR4
D39
JE2
Cowan1
D39
D39
D39
600 400 200 0
Euclidean distance
9 h
JE2
JE2
JE2
Cowan1
Cowan1
Cowan1
Control
Control
TIGR4
TIGR4
TIGR4
Control
Control
TIGR4
D39
JE2
Cowan1
D39
D39
D39
600 400 200 0
Euclidean distance
10 h
Control
Control
Control
TIGR4
Control
TIGR4
TIGR4
TIGR4
Cowan1
Cowan1
D39
Cowan1
Cowan1
JE2
JE2
JE2
JE2
D39
D39
D39
600 400 200 0
Euclidean distance
11 h
D39
TIGR4
TIGR4
Control
TIGR4
Control
TIGR4
Control
Control
Cowan1
Cowan1
Cowan1
Cowan1
JE2
JE2
JE2
JE2
D39
D39
D39
600 400 200 0
Euclidean distance
13 h
JE2
JE2
JE2
JE2
Cowan1
Cowan1
Cowan1
Cowan1
TIGR4
TIGR4
Control
TIGR4
Control
Control
Control
TIGR4
D39
D39
D39
D39
600 400 200 0
Euclidean distance
14 h
JE2
JE2
JE2
JE2
Cowan1
Cowan1
Cowan1
Cowan1
Control
TIGR4
Control
Control
TIGR4
Control
TIGR4
TIGR4
D39
D39
D39
D39
600 400 200 0
Euclidean distance
15 h
JE2
JE2
JE2
JE2
Cowan1
Cowan1
Cowan1
Cowan1
TIGR4
TIGR4
Control
TIGR4
Control
Control
Control
D39
TIGR4
D39
D39
D39
600 400 200 0
Euclidean distance

### TABLES

### Table S1

| | | | | | *S. pneumoniae* | | *S. aureus* | |
|---|---|---|---|---|---|---|---|---|
| *m/z* | Molecular formula[a] | Adduct | Ion mode | Mass error (ppm) | D39 | TIGR4 | JE2 | Cowan1 |
| 56.99994 | NA | NA | + | NA | | | x | |
| 57.00405 | NA | NA | + | NA | | | x | |
| 57.99834 | NA | NA | + | NA | | | x | |
| 58.96441 | NA | NA | - | NA | | | | x |
| 58.96477 | NA | NA | - | NA | | | | x |
| 59.52106 | NA | NA | + | NA | x | | | |
| 59.99688 | NA | NA | + | NA | | | x | |
| 60.98584 | H2ON2S | [M-H2O+H]+ | + | 4.420 | | | x | |
| 60.98633 | NA | NA | + | NA | | | x | |
| 61.00421 | H3O2N3 | [M+NH3-H]- | - | -1.825 | x | | | |
| 61.99417 | NA | NA | + | NA | | | x | |
| 69.99937 | C4H6S | [M-NH3+H]+ | + | -4.398 | | | x | |
| 74.01998 | NA | NA | + | NA | | | x | |
| 74.95278 | NA | NA | + | NA | | | x | |
| 75.00968 | NA | NA | + | NA | | | x | |
| 76.01749 | NA | NA | + | NA | | | x | |
| 77.01276 | NA | NA | + | NA | | | x | |
| 78.01297 | NA | NA | + | NA | | | x | |
| 78.99698 | C8 | [M-H2O+H]+ | + | 2.792 | | | x | |
| 80.00853 | NA | NA | + | NA | | | x | |
| 83.02531 | NA | NA | + | NA | | | x | |
| 84.01499 | C5H8S | [M-NH3+H]+ | + | -4.082 | | | x | |
| 85.52444 | NA | NA | + | NA | | | x | |
| 86.01803 | NA | NA | + | NA | | | x | x |
| 86.96266 | NA | NA | + | NA | | | x | |
| 86.96996 | NA | NA | - | NA | | | | x |
| 87.00429 | NA | NA | - | NA | | | | x |
| 88.01658 | NA | NA | - | NA | | | | x |
| 88.98124 | NA | NA | + | NA | | | x | |
| 89.01996 | NA | NA | - | NA | | | | x |
| 90.97661 | NA | NA | + | NA | | | x | |
| 91.03892 | C3H9O3N | [M-NH3+H]+ | + | -0.471 | | x | | |
| 92.03061 | NA | NA | + | NA | | | x | |
| 92.53226 | NA | NA | + | NA | | | x | |
| 93.09097 | C4H12O2 | [M+H]+ | + | -0.389 | x | x | | |
| 93.52197 | NA | NA | + | NA | | | x | x |
| 94.02712 | NA | NA | + | NA | | | x | |
| 94.02809 | NA | NA | + | NA | | | x | x |
| 95.03954 | NA | NA | + | NA | | | x | |
| 95.0467 | C4H8O | [M+Na]+ | + | -0.498 | | | x | |
| 95.96791 | NA | NA | + | NA | | | x | |
| 95.97429 | NA | NA | + | NA | | | x | x |
| 96.02355 | C8H3N | [M-H2O+H]+ | + | 2.557 | | | x | |
| 96.95 | NA | NA | + | NA | | | x | x |

| | | | | | | | | |
|---|---|---|---|---|---|---|---|---|
| 97.00761 | C8H2O | [M-H2O+H]+ | + | 2.924 | | | x | |
| 97.02065 | C5H7OP | [M-H2O+H]+ | + | 4.270 | | | x | |
| 97.026 | C3H6O2 | [M+Na]+ | + | -0.007 | | | x | |
| 97.97251 | H3ONS2 | [M+H]+ | + | -3.832 | | | x | x |
| 98.02783 | C9H6 | [M-NH3+H]+ | + | 1.336 | | | x | |
| 98.9753 | NA | NA | + | NA | | | x | |
| 99.01183 | NA | NA | + | NA | | | x | |
| 100.06301 | NA | NA | + | NA | | | | x |
| 100.93202 | OP2 | [M+Na]+ | + | 4.633 | | | x | |
| 101.00751 | NA | NA | + | NA | | | x | |
| 103.06142 | NA | NA | + | NA | | | x | |
| 104.03058 | NA | NA | + | NA | | | x | |
| 105.00006 | NA | NA | - | NA | | | x | |
| 105.0423 | C4H6N2 | [M+Na]+ | + | -0.235 | | | x | |
| 105.50907 | NA | NA | + | NA | | | x | |
| 106.00794 | C9HN | [M-H2O+H]+ | + | 2.675 | | | x | |
| 106.96641 | NA | NA | + | NA | | | x | |
| 106.99196 | C9O | [M-H2O+H]+ | + | 2.689 | | | x | |
| 107.02538 | NA | NA | + | NA | | | x | |
| 107.95045 | NA | NA | + | NA | | | x | |
| 108.02106 | NA | NA | + | NA | x | x | | |
| 108.042 | C4H7ON | [M+Na]+ | + | 0.178 | | | x | |
| 108.98763 | NA | NA | + | NA | | | x | |
| 109.02287 | NA | NA | + | NA | | | x | |
| 109.02605 | C4H6O2 | [M+Na]+ | + | 0.575 | | | x | |
| 111.03846 | NA | NA | + | NA | | | x | |
| 111.04169 | C4H8O2 | [M+Na]+ | + | 0.449 | | | x | |
| 111.05289 | C3H8ON2 | [M+Na]+ | + | 0.069 | | | x | |
| 112.04503 | NA | NA | + | NA | | | x | |
| 112.04626 | C7H12S | [M-NH3+H]+ | + | -3.422 | | | x | |
| 113.05012 | C8H6N2 | [M-H2O+H]+ | + | 2.384 | | | x | |
| 113.05724 | C4H10O2 | [M+Na]+ | + | -0.672 | | | x | |
| 113.94298 | NA | NA | - | NA | x | x | | |
| 114.0342 | H6N2P2 | [M+NH4]+ | + | -2.569 | | | x | |
| 114.04378 | C6H10O3 | [M-NH3+H]+ | + | 0.451 | | | x | x |
| 114.51439 | NA | NA | + | NA | | | x | |
| 114.98662 | NA | NA | - | NA | x | x | | |
| 115.03654 | C3H8O3 | [M+Na]+ | + | -0.273 | | | x | |
| 115.96338 | NA | NA | + | NA | | | x | |
| 116.94738 | NA | NA | + | NA | | | x | |
| 116.97617 | NA | NA | + | NA | | | x | |
| 117.0409 | NA | NA | + | NA | | | x | |
| 117.09212 | C6H15O2N | [M+NH3-H]- | - | 0.131 | x | | | |
| 118.10562 | NA | NA | + | NA | | | | x |
| 118.98961 | H6N2P2 | [M+Na]+ | + | -2.420 | | | x | |
| 119.0542 | H8O2N4 | [M+Na]+ | + | 2.639 | | | x | |
| 119.94322 | C6O2S | [M-NH3+H]+ | + | 4.358 | | | x | |
| 120.02364 | C10H3N | [M-H2O+H]+ | + | 2.766 | | | x | |
| 120.95314 | NA | NA | + | NA | | | x | |
| 121.0075 | C10H2O | [M-H2O+H]+ | + | 1.619 | | | x | |
| 122.00767 | NA | NA | + | NA | | | x | |
| 122.01994 | NA | NA | + | NA | | | x | |
| 122.05764 | C5H9ON | [M+Na]+ | + | 0.051 | | | x | |
| 122.50284 | NA | NA | + | NA | | | x | |

| | | | | | | | |
|---|---|---|---|---|---|---|---|
| 122.98797 | C9HON | [M+NH3-H]- | - | 2.388 | | | x |
| 123.00384 | NA | NA | + | NA | | x | |
| 123.02325 | C10H4O | [M-H2O+H]+ | + | 2.310 | | x | |
| 123.03439 | C9H4N2 | [M-H2O+H]+ | + | 1.642 | | x | |
| 123.0609 | NA | NA | + | NA | | x | |
| 123.07807 | C6H12O | [M+Na]+ | + | 0.341 | | x | |
| 123.51965 | NA | NA | + | NA | | x | |
| 124.01857 | C9H3ON | [M-H2O+H]+ | + | 2.797 | | x | |
| 124.0548 | C10H7N | [M-H2O+H]+ | + | 1.694 | | x | |
| 125.00248 | C9H2O2 | [M-H2O+H]+ | + | 2.034 | | x | |
| 125.05737 | C5H10O2 | [M+Na]+ | + | 0.681 | | x | |
| 126.01408 | NA | NA | + | NA | | x | |
| 127.00667 | NA | NA | + | NA | | x | |
| 129.98116 | C8H2O3 | [M-NH3+H]+ | + | 0.264 | | x | |
| 132.55952 | NA | NA | + | NA | | x | x |
| 133.04619 | NA | NA | - | NA | x | | |
| 134.00294 | C10HON | [M-H2O+H]+ | + | 2.744 | | x | |
| 134.03919 | C11H5N | [M-H2O+H]+ | + | 1.847 | | x | |
| 134.5299 | NA | NA | + | NA | | x | |
| 134.98679 | C10O2 | [M-H2O+H]+ | + | 1.637 | | x | |
| 135.00504 | C5H4O3 | [M+Na]+ | + | -2.010 | | x | x |
| 135.0231 | C11H4O | [M-H2O+H]+ | + | 1.141 | | x | |
| 135.04156 | C6H8O2 | [M+Na]+ | + | -0.808 | | x | |
| 135.99013 | NA | NA | + | NA | | x | |
| 136.01863 | C10H3ON | [M-H2O+H]+ | + | 2.970 | | x | |
| 136.03486 | NA | NA | + | NA | | x | |
| 136.03676 | C5H7O2N | [M+Na]+ | + | -1.234 | x | | |
| 136.04626 | C9H12S | [M-NH3+H]+ | + | -2.882 | | x | |
| 136.05483 | C11H7N | [M-H2O+H]+ | + | 1.758 | | x | |
| 136.07326 | C6H11ON | [M+Na]+ | + | -0.220 | | x | |
| 136.99109 | C4H10S3 | [M-H2O+H]+ | + | -0.546 | | x | |
| 137.00203 | C10H2O2 | [M-H2O+H]+ | + | -1.047 | | x | |
| 137.0388 | C11H6O | [M-H2O+H]+ | + | 1.450 | | x | |
| 137.07261 | C5H15O3P | [M-H2O+H]+ | + | 0.115 | | x | |
| 138.03409 | C10H5ON | [M-H2O+H]+ | + | 1.705 | | x | |
| 138.04072 | NA | NA | - | NA | | | x |
| 138.04533 | C9H5N3 | [M-H2O+H]+ | + | 1.748 | | x | |
| 138.05047 | NA | NA | + | NA | | x | |
| 138.96367 | C9OS | [M-H2O+H]+ | + | -0.175 | | x | |
| 139.01804 | C10H4O2 | [M-H2O+H]+ | + | 1.274 | | x | |
| 139.02931 | C9H4ON2 | [M-H2O+H]+ | + | 1.509 | | x | |
| 139.03439 | NA | NA | + | NA | | x | |
| 139.07292 | C6H12O2 | [M+Na]+ | + | -0.263 | | x | |
| 140.01398 | C4H7ONS | [M+Na]+ | + | -0.646 | | x | |
| 140.0304 | NA | NA | + | NA | | x | |
| 141.0143 | NA | NA | + | NA | | x | |
| 141.01742 | C5H2N4 | [M+Na]+ | + | 2.141 | | x | |
| 141.08855 | C6H14O2 | [M+Na]+ | + | -0.428 | | x | |
| 141.97974 | NA | NA | + | NA | | x | x |
| 142.00977 | C10H7P | [M-NH3+H]+ | + | 3.201 | | x | |
| 143.01014 | C5ON6 | [M-H2O+H]+ | + | 0.434 | | x | |
| 143.04667 | C8H8O | [M+Na]+ | + | -0.549 | | x | |
| 143.05015 | C10H9ON | [M+NH3-H]- | - | -0.553 | | x | x |
| 143.97794 | NA | NA | + | NA | | x | x |

| | | | | | | |
|---|---|---|---|---|---|---|
| 144.96034 | C5H6S3 | [M-H2O+H]+ | + | 2.877 | x | x |
| 144.992 | C8H2O4 | [M-H2O+H]+ | + | -0.126 | x | |
| 147.02314 | C12H4O | [M-H2O+H]+ | + | 1.301 | x | |
| 147.99166 | C8H4O4 | [M-NH3+H]+ | + | -0.159 | x | |
| 149.02102 | C6H6O3 | [M+Na]+ | + | 0.832 | x | |
| 149.03886 | C12H6O | [M-H2O+H]+ | + | 1.707 | x | |
| 149.9633 | C7H3O3P | [M-NH3+H]+ | + | 3.567 | x | x |
| 149.99769 | C10HO2N | [M-H2O+H]+ | + | 1.496 | x | |
| 150.03411 | C11H5ON | [M-H2O+H]+ | + | 1.703 | x | |
| 150.07048 | C12H9N | [M-H2O+H]+ | + | 1.610 | x | |
| 150.98168 | C10O3 | [M-H2O+H]+ | + | 1.335 | x | |
| 150.98278 | C10HO2N | [M+NH3-H]- | - | 1.362 | | x |
| 151.0181 | C11H4O2 | [M-H2O+H]+ | + | 1.540 | x | |
| 151.05432 | C12H8O | [M-H2O+H]+ | + | 0.556 | x | |
| 152.01338 | C10H3O2N | [M-H2O+H]+ | + | 1.715 | x | |
| 152.04976 | C11H7ON | [M-H2O+H]+ | + | 1.682 | x | |
| 152.99735 | C10H2O3 | [M-H2O+H]+ | + | 1.436 | x | |
| 153.03135 | C4H12O4NP | [M-NH3+H]+ | + | 1.352 | x | |
| 153.0337 | C11H6O2 | [M-H2O+H]+ | + | 1.228 | x | |
| 153.05021 | NA | NA | + | NA | x | |
| 153.08861 | C7H14O2 | [M+Na]+ | + | 0.073 | x | |
| 153.98399 | C9H2ONP | [M-H2O+H]+ | + | -0.714 | x | x |
| 154.04551 | NA | NA | + | NA | x | |
| 155.10429 | C7H16O2 | [M+Na]+ | + | 0.299 | x | |
| 155.99109 | C4H7NS2 | [M+Na]+ | + | -0.915 | x | |
| 156.97433 | C9H2O2S | [M-H2O+H]+ | + | 0.391 | x | |
| 156.99469 | C8H3ON2P | [M-H2O+H]+ | + | -1.845 | x | |
| 157.98689 | C8HO4N | [M-H2O+H]+ | + | -2.168 | x | |
| 159.04161 | C8H8O2 | [M+Na]+ | + | -0.298 | x | |
| 161.01202 | C5H3N6P | [M-H2O+H]+ | + | -2.093 | x | |
| 161.0573 | C8H10O2 | [M+Na]+ | + | -0.004 | x | |
| 162.01539 | C4H9ON3S2 | [M-H2O+H]+ | + | -0.140 | x | |
| 162.97042 | C6H2O2N3P | [M+NH3-H]- | - | 0.743 | x | x |
| 163.01806 | C12H4O2 | [M-H2O+H]+ | + | 1.215 | x | |
| 163.07307 | C8H12O2 | [M+Na]+ | + | 0.853 | x | |
| 164.01334 | C11H3O2N | [M-H2O+H]+ | + | 1.380 | x | |
| 164.04582 | C7H7O3N3 | [M-H2O+H]+ | + | 2.028 | x | x |
| 164.99738 | C4H10N2S3 | [M-H2O+H]+ | + | 0.318 | x | |
| 165.03374 | C12H6O2 | [M-H2O+H]+ | + | 1.367 | x | |
| 165.04264 | C4H11O4N2P | [M-H2O+H]+ | + | 1.566 | x | x |
| 165.08867 | C8H14O2 | [M+Na]+ | + | 0.489 | x | |
| 166.04828 | C4H14N4S2 | [M+NH3-H]- | - | 2.572 | | x |
| 167.02428 | C10H4O2N2 | [M-H2O+H]+ | + | 1.580 | x | |
| 167.02941 | C7H10N2P2 | [M-H2O+H]+ | + | 4.222 | x | |
| 167.04941 | C12H8O2 | [M-H2O+H]+ | + | 1.460 | x | |
| 167.0603 | C11H8ON2 | [M-H2O+H]+ | + | -0.405 | x | |
| 168.00823 | C10H3O3N | [M-H2O+H]+ | + | 1.217 | x | |
| 168.95853 | C8ON3P | [M-NH3+H]+ | + | -0.517 | x | |
| 169.05831 | C5H10O3N2 | [M+Na]+ | + | -0.364 | x | x |
| 173.01683 | C9H6ON2S | [M-H2O+H]+ | + | 0.183 | x | |
| 173.97478 | C7H10S3 | [M-NH3+H]+ | + | -2.156 | x | |
| 174.00087 | C9H5O2NS | [M-H2O+H]+ | + | 0.309 | x | |
| 174.95884 | C7O3N2S | [M-H2O+H]+ | + | -4.347 | x | |
| 174.98488 | C9H4O3S | [M-H2O+H]+ | + | 0.279 | x | |

| | | | | | | |
|---|---|---|---|---|---|---|
| 175.00081 | C13H4S | [M-H2O+H]+ | + | 3.788 | x | |
| 175.98442 | C13H4S | [M-NH3+H]+ | + | 1.675 | x | |
| 175.98996 | C6H8O5S | [M-NH3+H]+ | + | -0.059 | x | |
| 176.99599 | C6H3O2N4P | [M-H2O+H]+ | + | -0.431 | x | x |
| 177.03378 | C13H6O2 | [M-H2O+H]+ | + | 1.488 | x | |
| 177.07013 | C14H10O | [M-H2O+H]+ | + | 1.306 | x | |
| 177.08867 | C9H14O2 | [M+Na]+ | + | 0.451 | x | |
| 177.99269 | C4H9N3S3 | [M-H2O+H]+ | + | 0.610 | x | |
| 179.01307 | C5H12N2S3 | [M-H2O+H]+ | + | 0.499 | x | |
| 179.04935 | C13H8O2 | [M-H2O+H]+ | + | 1.065 | x | |
| 179.10422 | C9H16O2 | [M+Na]+ | + | -0.195 | x | |
| 180.06395 | C5H2ON6 | [M-NH4]- | - | 0.117 | | x |
| 180.99984 | C6H6N4S2 | [M-H2O+H]+ | + | -1.310 | x | |
| 181.0287 | C5H14N2S3 | [M-H2O+H]+ | + | 0.393 | x | |
| 181.12001 | C9H18O2 | [M+Na]+ | + | 0.692 | x | |
| 182.024 | C4H13N3S3 | [M-H2O+H]+ | + | 0.648 | x | |
| 182.95573 | C6HO7N | [M-NH3+H]+ | + | -1.502 | x | |
| 183.97914 | C6H4O5NP | [M-H2O+H]+ | + | -1.396 | x | x |
| 184.96929 | C10H2O3S | [M-H2O+H]+ | + | 0.562 | x | |
| 185.05728 | C10H10O2 | [M+Na]+ | + | -0.126 | x | |
| 185.11504 | C8H18O3 | [M+Na]+ | + | 1.387 | x | |
| 186.11819 | C7H17N5S | [M-H2O+H]+ | + | 4.984 | x | |
| 189.98088 | C13H2O3 | [M-NH3+H]+ | + | -1.172 | x | |
| 191.08539 | C15H12O | [M-H2O+H]+ | + | -0.656 | x | |
| 192.97936 | C4H2O2N4S | [M+Na]+ | + | 1.722 | x | |
| 192.99238 | C5H10ON2S3 | [M-H2O+H]+ | + | 0.682 | x | |
| 193.02317 | C7H2N6 | [M+Na]+ | + | -0.855 | x | x |
| 193.02868 | C6H14N2S3 | [M-H2O+H]+ | + | 0.276 | x | |
| 193.06498 | C14H10O2 | [M-H2O+H]+ | + | 0.899 | x | |
| 193.11992 | C10H18O2 | [M+Na]+ | + | 0.115 | x | |
| 193.13104 | C9H18ON2 | [M+Na]+ | + | -0.552 | x | |
| 193.94016 | C4H5O3NS3 | [M-H2O+H]+ | + | 1.389 | x | x |
| 194.96037 | C6H2O4N3P | [M+NH3-H]- | - | 1.202 | x | x |
| 194.96611 | C10H2N2P2 | [M-H2O+H]+ | + | 0.363 | x | |
| 195.03778 | C6H8O4N2 | [M+Na]+ | + | 0.885 | x | |
| 196.00313 | C4H11ON3S3 | [M-H2O+H]+ | + | -0.027 | x | |
| 196.02168 | C6H7O5N | [M+Na]+ | + | 0.212 | x | |
| 196.9544 | C8H6OS3 | [M-H2O+H]+ | + | -1.817 | x | |
| 197.03332 | C5H7O2N6P | [M-H2O+H]+ | + | -0.942 | x | |
| 197.11498 | C9H18O3 | [M+Na]+ | + | 0.947 | x | |
| 198.01746 | C5H6O3N5P | [M-H2O+H]+ | + | -0.360 | x | |
| 201.98434 | C11H6O3S | [M-NH3+H]+ | + | -0.698 | x | |
| 204.9861 | C7H6N2S2 | [M+Na]+ | + | -1.982 | x | x |
| 205.1012 | C16H14O | [M-H2O+H]+ | + | 0.083 | x | |
| 206.96037 | C5O6S | [M-H2O-H]- | - | -0.672 | x | x |
| 206.98477 | C9H5O5P | [M-H2O+H]+ | + | 2.672 | x | x |
| 207.04434 | C7H16N2S3 | [M-H2O+H]+ | + | 0.303 | x | |
| 207.08052 | C15H12O2 | [M-H2O+H]+ | + | 0.352 | x | |
| 207.09914 | C10H16O3 | [M+Na]+ | + | -0.136 | x | |
| 211.13041 | C10H20O3 | [M+Na]+ | + | -0.293 | x | |
| 213.10966 | C15H18S | [M-H2O+H]+ | + | 0.119 | x | |
| 215.0161 | C12H8O3S | [M-H2O+H]+ | + | -0.114 | x | |
| 216.95346 | C4H6ON2S3 | [M+Na]+ | + | 0.072 | x | |
| 216.99531 | C11H6O4S | [M-H2O+H]+ | + | -0.347 | x | |

| | | | | | | |
|---|---|---|---|---|---|---|
| 217.10461 | C8H18O5 | [M+Na]+ | + | -0.177 | x | |
| 217.96158 | C11H6O2S2 | [M-NH3+H]+ | + | -0.291 | x | |
| 218.10781 | C15H13N3 | [M-H2O+H]+ | + | 0.642 | x | |
| 219.08061 | C16H12O2 | [M-H2O+H]+ | + | 0.715 | x | |
| 219.09033 | C9H13N6P | [M-H2O+H]+ | + | -1.324 | x | |
| 219.99488 | C11H8O4S | [M-NH3+H]+ | + | -0.749 | x | |
| 222.12776 | C16H17ON | [M-H2O+H]+ | + | 0.144 | x | |
| 222.96128 | C14S | [M+Na]+ | + | -0.060 | x | |
| 223.02773 | C8H16O2S3 | [M-H2O+H]+ | + | -0.930 | x | |
| 223.07538 | C15H12O3 | [M-H2O+H]+ | + | 0.101 | x | |
| 223.0938 | C8H12O3N6 | [M-H2O+H]+ | + | 0.001 | x | |
| 225.95099 | C5H2O6NP | [M+Na]+ | + | -1.007 | x | |
| 227.9479 | C10O5NP | [M-H2O+H]+ | + | -0.900 | x | |
| 228.14946 | C14H19ON3 | [M-H2O+H]+ | + | -0.259 | x | x |
| 228.99542 | C12H6O4S | [M-H2O+H]+ | + | 0.117 | x | |
| 229.03178 | C13H10O3S | [M-H2O+H]+ | + | 0.014 | x | |
| 229.10455 | C15H18OS | [M-H2O+H]+ | + | 0.011 | x | |
| 231.0111 | C12H8O4S | [M-H2O+H]+ | + | 0.237 | x | |
| 231.93742 | C14O3S | [M-NH3+H]+ | + | -0.492 | x | |
| 233.06957 | C9H11ON6P | [M-H2O+H]+ | + | -1.348 | x | |
| 233.10583 | C10H15N6P | [M-H2O+H]+ | + | -1.849 | x | |
| 233.93388 | C6H5O4NS3 | [M-H2O+H]+ | + | -3.593 | x | |
| 233.96759 | C6H2O4N3P | [M+Na]+ | + | 0.364 | x | |
| 234.00108 | C8H5O6N | [M+Na]+ | + | 0.816 | x | |
| 234.14637 | C12H21O2N | [M+Na]+ | + | -0.376 | x | |
| 234.91819 | C5N4S3 | [M+Na]+ | + | 2.173 | x | |
| 234.9516 | C6HO5N2P | [M+Na]+ | + | 0.336 | x | |
| 235.07567 | C9H20N2S3 | [M-H2O+H]+ | + | 0.388 | x | |
| 235.11201 | C17H16O2 | [M-H2O+H]+ | + | 1.066 | x | |
| 235.97215 | C11H8O3S2 | [M-NH3+H]+ | + | -0.249 | x | |
| 237.10096 | C9H15ON6P | [M-H2O+H]+ | + | -0.972 | x | |
| 237.12738 | C17H18O2 | [M-H2O+H]+ | + | -0.044 | x | |
| 238.00553 | C11H10O5S | [M-NH3+H]+ | + | -0.359 | x | |
| 239.08883 | C16H16OS | [M-H2O+H]+ | + | -0.263 | x | |
| 239.16153 | C10H20O2N6 | [M-H2O+H]+ | + | 0.175 | x | |
| 240.09227 | C17H11N3 | [M-H2O+H]+ | + | 1.015 | x | |
| 241.92815 | C8H2O8S | [M-NH3+H]+ | + | 1.492 | x | |
| 242.9776 | C5H5O6N2P | [M+Na]+ | + | -0.652 | x | |
| 243.92503 | C8H4O6S2 | [M-NH3+H]+ | + | -2.411 | x | |
| 243.96153 | C13HN3P2 | [M-H2O+H]+ | + | 0.950 | x | |
| 244.97443 | C10H3O5N2P | [M-H2O+H]+ | + | -0.914 | x | |
| 245.95848 | C10H2O6NP | [M-H2O+H]+ | + | -0.780 | x | |
| 246.97332 | C8O7N4 | [M-H2O+H]+ | + | -0.342 | x | |
| 248.10724 | C17H15O2N | [M-H2O+H]+ | + | 0.942 | x | |
| 249.0912 | C10H22N2S3 | [M-H2O+H]+ | + | -0.083 | x | |
| 251.21163 | C15H28O2N2 | [M-H2O+H]+ | + | -0.594 | | x |
| 251.97809 | C6H4O5N3P | [M+Na]+ | + | 0.053 | x | |
| 252.05722 | C15H12O2NP | [M-H2O+H]+ | + | -0.211 | x | x |
| 252.94469 | C10H6O3S3 | [M-H2O+H]+ | + | 0.267 | x | |
| 252.96211 | C6H3O6N2P | [M+Na]+ | + | 0.072 | x | |
| 256.08866 | C15H16O2NP | [M-H2O+H]+ | + | 0.305 | x | |
| 256.97436 | C11H3O5N2P | [M-H2O+H]+ | + | -1.130 | x | x |
| 258.08544 | C6H18O7N3P | [M-H2O+H]+ | + | 1.790 | x | |
| 258.92479 | C8O3N2S2 | [M+Na]+ | + | 2.270 | x | |

| | | | | | | | | |
|---|---|---|---|---|---|---|---|---|
| 258.93738 | C9H8O2S4 | [M-H2O+H]+ | + | -0.159 | | | x | |
| 258.95478 | C7H2O2N4P2 | [M+Na]+ | + | 1.104 | | | x | |
| 259.11532 | C10H20O6 | [M+Na]+ | + | 0.471 | | | x | |
| 260.95158 | C12ON4P2 | [M-H2O+H]+ | + | 0.484 | | | x | |
| 261.1009 | C11H15ON6P | [M-H2O+H]+ | + | -1.104 | | | x | |
| 261.1204 | C9H19O5N5 | [M+NH3-H]- | - | -0.153 | | | x | |
| 261.13067 | C8H18O5N6 | [M-H2O+H]+ | + | 0.327 | | | x | |
| 262.13417 | C17H17ON3 | [M-H2O+H]+ | + | 1.062 | | | x | |
| 262.95051 | C7H6O8P2 | [M-H2O+H]+ | + | 0.029 | | | x | |
| 263.12522 | C19H20S | [M-H2O+H]+ | + | -0.224 | | | x | |
| 266.95083 | C12H2O3N2P2 | [M-H2O+H]+ | + | 0.187 | | | x | |
| 267.15661 | C19H24S | [M-H2O+H]+ | + | 0.096 | | | x | |
| 268.07147 | C14H11O4N3 | [M-H2O+H]+ | + | -0.692 | | | x | x |
| 269.13577 | C18H22OS | [M-H2O+H]+ | + | -0.253 | | | x | |
| 269.13943 | C14H23O5N | [M+NH3-H]- | - | -0.058 | x | x | | |
| 269.17556 | C15H27O4N | [M+NH3-H]- | - | -0.953 | x | x | | |
| 269.19743 | C7H28ON9P | [M+NH3-H]- | - | 0.573 | | | x | x |
| 271.10413 | C10H25O4NS2 | [M+NH3-H]- | - | -0.673 | | | x | x |
| 271.11867 | C13H21O6N | [M+NH3-H]- | - | -0.143 | x | x | | |
| 271.21306 | C7H24ON7P | [M-NH4]- | - | 0.567 | | | x | x |
| 271.22782 | C16H33O3N | [M+NH3-H]- | - | -0.164 | x | x | | |
| 271.9231 | C5H3O3N3S3 | [M+Na]+ | + | 0.907 | | | x | x |
| 272.11691 | C17H20O4 | [M-NH3+H]+ | + | 0.083 | | | x | x |
| 272.94338 | C6H6O3N2S3 | [M+Na]+ | + | 0.419 | | | x | |
| 273.05788 | C7H10O7N6 | [M-H2O+H]+ | + | 0.248 | | | x | |
| 273.10088 | C10H24O2S3 | [M+H]+ | + | -0.874 | | | x | x |
| 273.1672 | C12H26O5 | [M+Na]+ | + | -0.177 | | | x | |
| 273.19016 | C8H25N5S2 | [M-NH4]- | - | 0.397 | | | x | x |
| 274.09617 | C16H18O5 | [M-NH3+H]+ | + | 0.067 | | | x | x |
| 274.93202 | C9H8O3S4 | [M-H2O+H]+ | + | -1.091 | | | x | |
| 275.16149 | C13H20O2N6 | [M-H2O+H]+ | + | 0.016 | | | x | |
| 278.92178 | C8H2O6P2 | [M+Na]+ | + | -0.398 | | | x | |
| 280.05217 | C16H12O3NP | [M-H2O+H]+ | + | -0.072 | | | x | x |
| 280.93962 | C11H6O4S3 | [M-H2O+H]+ | + | 0.310 | | | x | |
| 281.17226 | C20H26S | [M-H2O+H]+ | + | 0.092 | | | x | |
| 282.0491 | C5H17ON5S3 | [M+Na]+ | + | 1.378 | | | x | x |
| 291.11146 | C12H17O2N6P | [M-H2O+H]+ | + | -1.012 | | | x | x |
| 296.90374 | C11HO3P3 | [M+Na]+ | + | 2.411 | | | x | x |
| 299.16167 | C17H24O3 | [M+Na]+ | + | -0.344 | | | x | |
| 299.18286 | C14H28O5 | [M+Na]+ | + | -0.124 | | | x | |
| 301.14095 | C7H26N8S3 | [M-H2O+H]+ | + | -0.051 | | | x | |
| 305.15699 | C18H26O3S | [M-H2O+H]+ | + | 0.042 | | | x | |
| 313.16517 | C16H27O6N | [M+NH3-H]- | - | -1.492 | x | x | | |
| 315.25361 | C18H37O4N | [M+NH3-H]- | - | -1.424 | x | x | | |
| 317.17215 | C23H26S | [M-H2O+H]+ | + | -0.247 | | | x | |
| 317.19647 | C16H31O6N | [M+NH3-H]- | - | -1.474 | x | x | | |
| 326.9398 | C8H2O9N5P | [M-NH3+H]+ | + | 0.173 | | | x | |
| 327.06867 | C12H16O9 | [M+Na]+ | + | 0.073 | | | x | x |
| 327.24979 | C14H31O6N | [M-NH4]- | - | -0.870 | | | x | x |
| 328.94565 | C8H2N8S3 | [M+Na]+ | + | -0.083 | | | x | |
| 328.95577 | C15H5N2P3 | [M+Na]+ | + | -0.029 | | | x | |
| 329.26522 | C14H33O6N | [M-NH4]- | - | -1.571 | | | x | x |
| 329.95501 | C10H5O11NS | [M-H2O+H]+ | + | -0.093 | | | x | |
| 330.26871 | C23H38O2 | [M+NH3-H]- | - | -0.848 | | | x | x |

| | | | | | | | | |
|---|---|---|---|---|---|---|---|---|
| 330.93395 | C10O5N6S2 | [M-H2O+H]+ | + | 0.229 | | | x | |
| 331.15154 | C8H28ON8S3 | [M-H2O+H]+ | + | 0.027 | | | x | |
| 333.15195 | C13H26O8 | [M+Na]+ | + | -0.123 | | | x | |
| 333.16712 | C23H26OS | [M-H2O+H]+ | + | -0.092 | | | x | |
| 335.05042 | C11H13ON9S2 | [M+NH3-H]- | - | 0.423 | x | x | | |
| 337.2012 | C13H32O4N5P | [M+NH3-H]- | - | 0.526 | x | x | | |
| 338.26652 | C18H37O3N | [M+Na]+ | + | -0.140 | | | x | |
| 345.94094 | C13H5O4N3S3 | [M-H2O+H]+ | + | 0.029 | | | x | |
| 347.17016 | C10H30O7N5P | [M+NH3-H]- | - | 0.143 | x | x | | |
| 347.95362 | C9H11ON5S5 | [M-H2O+H]+ | + | 0.548 | | | x | x |
| 349.05994 | C18H15O4P | [M+Na]+ | + | -0.234 | | | x | |
| 349.18324 | C14H30O8 | [M+Na]+ | + | -0.132 | | | x | |
| 350.23017 | C18H33O4N | [M+Na]+ | + | -0.027 | | | x | |
| 367.20243 | C22H29O3N3 | [M+NH3-H]- | - | -0.731 | | | x | x |
| 367.21121 | C17H25N11 | [M+NH3-H]- | - | -0.139 | x | x | | |
| 369.21799 | C14H35O5N5S | [M+NH3-H]- | - | 0.706 | | | x | x |
| 369.22682 | C17H27N11 | [M+NH3-H]- | - | -0.242 | x | x | | |
| 371.94745 | C13H11O8P3 | [M-NH3+H]+ | + | 0.119 | | | x | |
| 374.92236 | C15HON7S3 | [M+NH3-H]- | - | 0.169 | | | | x |
| 383.20577 | C17H33N7S2 | [M+NH3-H]- | - | 0.153 | x | x | | |
| 385.2124 | C16H32O3N7P | [M+NH3-H]- | - | 0.380 | | | x | x |
| 385.22141 | C17H35N7S2 | [M+NH3-H]- | - | 0.128 | x | x | | |
| 393.2976 | C22H42O4 | [M+Na]+ | + | 0.190 | | | x | |
| 407.18874 | C14H20N14 | [M+Na]+ | + | -0.032 | | | x | |

[a] Molecular formula based on accurate mass only (no isotopic pattern matching performed).

x = Presence of a particular *m/z* in the respective bacterial serotype.

NA= Not available

### Table S2

| | | | | | *S. pneumoniae* | | *S. aureus* | |
|---|---|---|---|---|---|---|---|---|
| *m/z* | Molecular formula | Adduct | Ion mode | Mass error (ppm) | D39 | TIGR4 | JE2 | Cowan1 |
| 55.95689 | NA | NA | + | NA | x | | | |
| 56.00194 | C3H4O2 | [M-NH3+H]+ | + | 1.155 | x | | | |
| 56.96425 | NA | NA | + | NA | x | | | |
| 56.96483 | NA | NA | + | NA | x | | | |
| 56.99994 | NA | NA | + | NA | x | | | |
| 57.00405 | NA | NA | + | NA | x | | | |
| 57.01843 | NA | NA | + | NA | x | | | |
| 57.97248 | NA | NA | + | NA | x | | | |
| 57.97568 | NA | NA | - | NA | x | x | | |
| 57.99834 | NA | NA | + | NA | x | | | |
| 58.96082 | NA | NA | + | NA | x | | | |
| 58.9978 | NA | NA | + | NA | x | | | |
| 59.03599 | NA | NA | + | NA | x | | | |
| 59.99688 | NA | NA | + | NA | x | | | |
| 59.9986 | NA | NA | + | NA | x | | | |
| 60.95766 | NA | NA | + | NA | x | | | |
| 60.98584 | H2ON2S | [M-H2O+H]+ | + | 4.420 | x | | | |
| 60.98633 | NA | NA | + | NA | x | | | |
| 60.99534 | H3ON2P | [M-H2O+H]+ | + | 4.217 | x | | | |
| 61.93556 | NA | NA | - | NA | x | | | |
| 61.99417 | NA | NA | + | NA | x | | | |
| 62.04138 | NA | NA | - | NA | x | | | |
| 63.92845 | NA | NA | + | NA | x | | | |
| 64.93628 | NA | NA | + | NA | x | | | |
| 65.00714 | NA | NA | + | NA | x | | | |
| 65.92535 | NA | NA | + | NA | x | | | |
| 66.00519 | NA | NA | + | NA | x | | | |
| 66.0093 | NA | NA | + | NA | x | | | |
| 66.93318 | NA | NA | + | NA | x | | | |
| 67.00359 | NA | NA | + | NA | x | | | |
| 68.93199 | NA | NA | + | NA | x | | | |
| 68.98206 | NA | NA | + | NA | x | | | |
| 68.99574 | NA | NA | - | NA | x | | | |
| 69.00212 | NA | NA | + | NA | x | | | |
| 69.99937 | C4H6S | [M-NH3+H]+ | + | -4.398 | x | | | |
| 70.01752 | C4H6O2 | [M-NH3+H]+ | + | 0.153 | x | | | |
| 71.00716 | NA | NA | + | NA | x | | | |
| 71.05022 | C4H9ON | [M+NH3-H]- | - | -0.207 | | | x | |
| 72.0536 | NA | NA | - | NA | | | x | |
| 73.02771 | NA | NA | + | NA | x | | | |
| 74.01242 | C3H6O3 | [M-NH3+H]+ | + | -0.016 | x | | | |
| 74.01998 | NA | NA | + | NA | x | | | |
| 74.95278 | NA | NA | + | NA | x | | | |
| 74.97466 | HO2N2P | [M-H2O+H]+ | + | 4.178 | x | | | |
| 75.0006 | NA | NA | + | NA | x | | | |
| 75.01046 | NA | NA | + | NA | x | | | |
| 75.01457 | NA | NA | + | NA | x | | | |
| 75.09167 | C3H12ON2 | [M-H2O+H]+ | + | -0.050 | x | | | |

| | | | | | | | | |
|---|---|---|---|---|---|---|---|---|
| 75.98305 | NA | NA | + | NA | x | | | |
| 76.00888 | NA | NA | + | NA | x | | | |
| 76.91496 | NA | NA | - | NA | | | x | x |
| 77.01276 | NA | NA | + | NA | x | | | |
| 77.95737 | NA | NA | + | NA | x | | | |
| 78.01297 | NA | NA | + | NA | x | | | |
| 78.01497 | NA | NA | + | NA | x | | | |
| 78.99698 | C8 | [M-H2O+H]+ | + | 2.792 | x | | | |
| 79.0046 | NA | NA | + | NA | x | | | |
| 79.02277 | NA | NA | + | NA | x | | | |
| 79.9472 | NA | NA | + | NA | x | | | |
| 79.95553 | NA | NA | + | NA | x | | | |
| 79.97336 | NA | NA | + | NA | x | | | |
| 80.00853 | NA | NA | + | NA | x | | | |
| 80.93122 | NA | NA | + | NA | x | | | |
| 80.93958 | NA | NA | + | NA | x | | | |
| 80.95504 | NA | NA | + | NA | x | | | |
| 80.99261 | NA | NA | + | NA | x | | | |
| 81.93902 | NA | NA | + | NA | x | | | |
| 81.95376 | NA | NA | + | NA | x | | | |
| 81.96292 | C4H2OS | [M-NH3+H]+ | + | -4.520 | x | | | |
| 82.02631 | C2H5ON | [M+Na]+ | + | -0.422 | x | | | |
| 82.02984 | C4H6ON2 | [M+NH3-H]- | - | 0.031 | x | x | | |
| 82.92808 | NA | NA | + | NA | x | | | |
| 82.93774 | NA | NA | + | NA | x | | | |
| 82.94693 | NA | NA | + | NA | x | | | |
| 82.95189 | NA | NA | + | NA | x | | | |
| 83.01769 | NA | NA | + | NA | x | | | |
| 83.02155 | NA | NA | + | NA | x | | | |
| 83.02531 | NA | NA | + | NA | x | | | |
| 83.93592 | NA | NA | + | NA | x | | | |
| 83.95984 | NA | NA | + | NA | x | | | |
| 84.00592 | NA | NA | + | NA | x | | | |
| 84.00909 | C3H4O2N2 | [M+NH3-H]- | - | -0.116 | x | | | |
| 84.04184 | C2H7ON | [M+Na]+ | + | -2.373 | x | | | |
| 84.92693 | NA | NA | + | NA | x | | | |
| 84.93828 | NA | NA | + | NA | x | | | |
| 84.94384 | NA | NA | + | NA | x | | | |
| 84.95073 | NA | NA | + | NA | x | | | |
| 84.95969 | NA | NA | + | NA | x | | | |
| 84.98117 | NA | NA | + | NA | x | | | |
| 85.9347 | NA | NA | + | NA | x | | | |
| 85.9586 | C3H2O2S | [M-NH3+H]+ | + | 3.162 | x | | | |
| 85.963 | NA | NA | + | NA | x | | | |
| 86.01803 | NA | NA | + | NA | x | | | |
| 86.03735 | H5O2P | [M-NH4]- | - | -4.237 | | | x | |
| 86.94267 | NA | NA | + | NA | x | | | |
| 86.95572 | NA | NA | + | NA | x | | | |
| 86.96266 | NA | NA | + | NA | x | | | |
| 86.96394 | NA | NA | + | NA | x | | | |
| 86.99268 | NA | NA | + | NA | x | | | |
| 87.02027 | C5H4 | [M+Na]+ | + | -3.925 | x | | | |
| 87.99347 | C4N4 | [M-NH3+H]+ | + | 4.292 | x | | | |
| 88.0505 | C2H2ON2 | [M+NH4]+ | + | -0.546 | x | | | |

| | | | | | | |
|---|---|---|---|---|---|---|
| 88.95252 | NA | NA | + | NA | x | |
| 88.97025 | C2H3O2NS | [M+NH3-H]- | - | -0.222 | x | |
| 88.97866 | NA | NA | + | NA | x | |
| 88.98124 | NA | NA | + | NA | x | |
| 88.99021 | NA | NA | + | NA | x | |
| 89.00741 | NA | NA | - | NA | x | |
| 89.01304 | NA | NA | - | NA | x | |
| 89.05873 | NA | NA | + | NA | | x |
| 89.10728 | C4H14ON2 | [M-H2O+H]+ | + | -0.420 | x | |
| 89.96368 | NA | NA | + | NA | x | |
| 89.99134 | NA | NA | - | NA | x | |
| 90.94767 | NA | NA | + | NA | x | |
| 91.06391 | NA | NA | + | NA | | x |
| 91.09805 | NA | NA | - | NA | x | |
| 92.0099 | H3O3N3 | [M-H]- | - | -2.840 | x | |
| 92.03061 | NA | NA | + | NA | x | |
| 92.53226 | NA | NA | + | NA | x | |
| 92.94145 | NA | NA | + | NA | x | |
| 92.94964 | NA | NA | + | NA | x | |
| 93.01128 | NA | NA | + | NA | x | |
| 93.51106 | NA | NA | + | NA | x | |
| 94.00972 | C6H7P | [M-NH3+H]+ | + | 4.143 | x | |
| 94.02712 | NA | NA | + | NA | x | |
| 94.02809 | NA | NA | + | NA | x | |
| 94.99052 | H2O2N2S | [M+H]+ | + | -4.836 | x | |
| 95.03954 | NA | NA | + | NA | x | |
| 95.96791 | NA | NA | + | NA | x | |
| 95.97429 | NA | NA | + | NA | x | |
| 96.02355 | C8H3N | [M-H2O+H]+ | + | 2.557 | x | |
| 96.04193 | C3H7ON | [M+Na]+ | + | -0.752 | x | |
| 96.95 | NA | NA | + | NA | x | |
| 96.98214 | NA | NA | + | NA | x | |
| 97.00761 | C8H2O | [M-H2O+H]+ | + | 2.924 | x | |
| 97.02065 | C5H7OP | [M-H2O+H]+ | + | 4.270 | x | |
| 97.03521 | NA | NA | + | NA | x | |
| 97.96615 | NA | NA | + | NA | x | |
| 97.97251 | H3ONS2 | [M+H]+ | + | -3.832 | x | |
| 98.01922 | NA | NA | + | NA | x | |
| 98.02783 | C9H6 | [M-NH3+H]+ | + | 1.336 | x | |
| 98.94184 | NA | NA | + | NA | x | |
| 98.95014 | NA | NA | + | NA | x | |
| 98.9753 | NA | NA | + | NA | x | |
| 98.98029 | O3N2 | [M+Na]+ | + | 2.328 | x | |
| 99.00324 | NA | NA | + | NA | x | |
| 99.01183 | NA | NA | + | NA | x | |
| 99.02026 | C6H4 | [M+Na]+ | + | -3.437 | x | |
| 99.04516 | C5H9O2N | [M+NH3-H]- | - | 0.064 | | x |
| 99.94964 | NA | NA | + | NA | x | |
| 99.96433 | NA | NA | + | NA | x | |
| 100.01485 | NA | NA | + | NA | x | |
| 100.04854 | NA | NA | - | NA | | x |
| 100.05298 | NA | NA | - | NA | | x |
| 100.93202 | OP2 | [M+Na]+ | + | 4.633 | x | |
| 100.93877 | NA | NA | + | NA | x | |

| | | | | | | | |
|---|---|---|---|---|---|---|---|
| 100.94834 | NA | NA | + | NA | x | | |
| 100.98803 | NA | NA | - | NA | x | | |
| 100.99892 | NA | NA | + | NA | x | | |
| 101.00751 | NA | NA | + | NA | x | | |
| 101.03295 | NA | NA | - | NA | | x | x |
| 101.03592 | C6H6 | [M+Na]+ | + | -3.220 | x | | |
| 101.04939 | NA | NA | - | NA | | x | |
| 101.06079 | C5H11O2N | [M+NH3-H]- | - | -0.108 | | x | x |
| 101.08864 | NA | NA | - | NA | | x | x |
| 101.93987 | NA | NA | + | NA | x | | |
| 101.98628 | C7H2O2 | [M-NH3+H]+ | + | 0.620 | x | | |
| 102.01656 | NA | NA | + | NA | x | | |
| 102.06415 | NA | NA | - | NA | | x | x |
| 102.06702 | C3H9ON3 | [M-H]- | - | -2.571 | | x | x |
| 102.51634 | NA | NA | + | NA | x | | |
| 102.51825 | NA | NA | + | NA | x | | |
| 102.93755 | NA | NA | + | NA | x | | |
| 102.94162 | NA | NA | + | NA | x | | |
| 102.97025 | NA | NA | + | NA | x | | |
| 102.99891 | NA | NA | + | NA | x | | |
| 103.0149 | NA | NA | + | NA | x | | |
| 103.06503 | NA | NA | - | NA | | x | x |
| 103.06748 | NA | NA | - | NA | | x | x |
| 103.97353 | NA | NA | + | NA | x | | |
| 103.99578 | C6H3NS | [M-H2O+H]+ | + | 3.705 | x | | |
| 104.01925 | NA | NA | + | NA | x | | |
| 104.10693 | C5H15O2N | [M-H2O+H]+ | + | -0.497 | | x | |
| 104.96628 | NA | NA | + | NA | x | | |
| 104.9733 | NA | NA | + | NA | x | | |
| 104.97451 | NA | NA | + | NA | x | | |
| 105.00329 | NA | NA | + | NA | x | | |
| 105.50907 | NA | NA | + | NA | x | | |
| 105.96648 | C3H6OS2 | [M-NH3+H]+ | + | -2.079 | x | | |
| 106.00408 | C4HO2N3 | [M-H2O+H]+ | + | 3.998 | x | | |
| 106.0066 | NA | NA | + | NA | x | | |
| 106.00794 | C9HN | [M-H2O+H]+ | + | 2.675 | x | | |
| 106.04634 | NA | NA | + | NA | x | | |
| 106.04988 | C3H10O3N2 | [M-NH3+H]+ | + | 0.087 | x | | |
| 106.08625 | C4H13O3N | [M-H2O+H]+ | + | -0.039 | | x | |
| 106.96317 | NA | NA | + | NA | x | | |
| 106.96641 | NA | NA | + | NA | x | | |
| 106.99196 | C9O | [M-H2O+H]+ | + | 2.689 | x | | |
| 107.00084 | NA | NA | + | NA | x | | |
| 107.00749 | N6 | [M+Na]+ | + | -2.086 | x | | |
| 107.08332 | C6H12 | [M+Na]+ | + | 2.363 | | x | |
| 107.08962 | NA | NA | + | NA | | x | |
| 107.09243 | NA | NA | + | NA | | x | |
| 107.95045 | NA | NA | + | NA | x | | |
| 107.96677 | NA | NA | + | NA | x | | |
| 108.042 | C4H7ON | [M+Na]+ | + | 0.178 | x | | |
| 108.09051 | NA | NA | + | NA | | x | |
| 108.92478 | NA | NA | + | NA | x | | |
| 108.92619 | NA | NA | + | NA | x | | |
| 108.9583 | NA | NA | + | NA | x | | |

| | | | | | | | | |
|---|---|---|---|---|---|---|---|---|
| 108.98763 | NA | NA | + | NA | x | | | |
| 109.02287 | NA | NA | + | NA | x | | | |
| 110.02126 | C3H5O2N | [M+Na]+ | + | 0.121 | x | | | |
| 110.92307 | NA | NA | + | NA | x | | | |
| 110.95201 | NA | NA | + | NA | x | | | |
| 110.96022 | NA | NA | + | NA | x | | | |
| 111.04169 | C4H8O2 | [M+Na]+ | + | 0.449 | | | x | x |
| 111.05289 | C3H8ON2 | [M+Na]+ | + | 0.069 | x | | | |
| 111.92422 | NA | NA | + | NA | x | | | |
| 112.03687 | C3H7O2N | [M+Na]+ | + | -0.331 | x | | | |
| 112.04503 | NA | NA | + | NA | | | x | x |
| 112.92187 | NA | NA | + | NA | x | | | |
| 113.02092 | C3H6O3 | [M+Na]+ | + | 0.054 | x | | | |
| 113.03457 | C4H6O3N2 | [M-H2O+H]+ | + | 0.125 | x | | | |
| 113.05012 | C8H6N2 | [M-H2O+H]+ | + | 2.384 | x | | | |
| 113.97852 | H3P3 | [M+NH4]+ | + | -0.679 | x | | | |
| 114.0342 | H6N2P2 | [M+NH4]+ | + | -2.569 | x | | | |
| 114.06615 | C4H9O2N3 | [M-H2O+H]+ | + | -0.291 | x | | | |
| 114.51439 | NA | NA | + | NA | x | | | |
| 114.97029 | NA | NA | + | NA | x | | | |
| 114.98662 | NA | NA | - | NA | x | | | |
| 115.0458 | NA | NA | + | NA | x | | | |
| 115.08667 | C5H12O2N2 | [M-H2O+H]+ | + | 0.612 | x | | | |
| 115.92391 | NA | NA | + | NA | x | | | |
| 115.96338 | NA | NA | + | NA | x | | | |
| 116.01084 | H3O4P | [M+NH4]+ | + | 1.220 | x | | | |
| 116.01417 | C2H7ONS | [M+Na]+ | + | 1.230 | x | | | |
| 116.02982 | NA | NA | + | NA | x | | | |
| 116.04789 | NA | NA | - | NA | | | x | |
| 116.08193 | C4H11O2N3 | [M-H2O+H]+ | + | 0.690 | x | | | |
| 116.94738 | NA | NA | + | NA | x | | | |
| 116.95238 | C6ONP | [M-NH3+H]+ | + | -0.735 | x | | | |
| 116.97617 | NA | NA | + | NA | x | | | |
| 116.98296 | NA | NA | - | NA | x | | | |
| 116.98585 | NA | NA | + | NA | x | | | |
| 117.0139 | NA | NA | + | NA | x | | | |
| 117.05572 | C5H11O3N | [M+NH3-H]- | - | 0.021 | | | x | |
| 117.09212 | C6H15O2N | [M+NH3-H]- | - | 0.131 | x | x | | |
| 117.9209 | NA | NA | + | NA | x | | | |
| 117.94577 | NA | NA | + | NA | x | | | |
| 117.95856 | H3ONP2 | [M+Na]+ | + | 3.708 | x | | | |
| 118.05909 | NA | NA | - | NA | | | x | |
| 118.94264 | H2O2P2 | [M+Na]+ | + | 4.341 | x | | | |
| 118.94942 | NA | NA | + | NA | x | | | |
| 119.00941 | C3H8N2S2 | [M-H2O+H]+ | + | -1.407 | x | | | |
| 119.01907 | NA | NA | + | NA | x | | | |
| 119.0542 | H8O2N4 | [M+Na]+ | + | 2.639 | x | | | |
| 119.05996 | NA | NA | - | NA | | | x | |
| 119.91963 | NA | NA | + | NA | x | | | |
| 119.94322 | C6O2S | [M-NH3+H]+ | + | 4.358 | x | | | |
| 119.95062 | NA | NA | + | NA | x | | | |
| 119.95894 | NA | NA | + | NA | x | | | |
| 119.96055 | C6O4 | [M-NH3+H]+ | + | 1.206 | x | | | |
| 119.99686 | C7H4O3 | [M-NH3+H]+ | + | 0.651 | x | | | |

| | | | | | | | |
|---|---|---|---|---|---|---|---|
| 120.02364 | C10H3N | [M-H2O+H]+ | + | 2.766 | x | | |
| 120.94305 | NA | NA | + | NA | x | | |
| 120.94448 | NA | NA | + | NA | x | | |
| 120.94815 | NA | NA | + | NA | x | | |
| 120.95314 | NA | NA | + | NA | x | | |
| 120.96621 | H3O4P | [M+Na]+ | + | 0.957 | x | | |
| 120.98088 | NA | NA | + | NA | x | | |
| 121.0075 | C10H2O | [M-H2O+H]+ | + | 1.619 | x | | |
| 121.01538 | NA | NA | + | NA | x | | |
| 121.01736 | C5H7P | [M+Na]+ | + | -4.058 | x | | |
| 121.06239 | C6H10O | [M+Na]+ | + | 0.042 | x | | |
| 121.06984 | NA | NA | + | NA | x | | |
| 121.0858 | C5H14O4 | [M-H2O+H]+ | + | -0.872 | x | | |
| 121.9841 | NA | NA | + | NA | x | | |
| 122.01994 | NA | NA | + | NA | x | | |
| 122.03214 | C3H5ON3 | [M+Na]+ | + | -3.462 | x | | |
| 122.05764 | C5H9ON | [M+Na]+ | + | 0.051 | x | | |
| 122.50284 | NA | NA | + | NA | x | | |
| 122.97675 | NA | NA | + | NA | x | | |
| 122.98504 | NA | NA | + | NA | x | | |
| 122.98797 | C9HON | [M+NH3-H]- | - | 2.388 | | | x |
| 123.00384 | NA | NA | + | NA | x | | |
| 123.01382 | NA | NA | + | NA | x | | |
| 123.02325 | C10H4O | [M-H2O+H]+ | + | 2.310 | x | | |
| 123.03439 | C9H4N2 | [M-H2O+H]+ | + | 1.642 | x | | |
| 123.04158 | C5H8O2 | [M+Na]+ | + | -0.755 | x | | |
| 123.0609 | NA | NA | + | NA | x | | |
| 123.51965 | NA | NA | + | NA | x | | |
| 123.55429 | NA | NA | + | NA | x | | |
| 123.97696 | C3H8O2S2 | [M-NH3+H]+ | + | -2.416 | x | | |
| 123.9887 | NA | NA | - | NA | | x | x |
| 124.0115 | NA | NA | + | NA | x | | |
| 124.01469 | C4H3O3N3 | [M-H2O+H]+ | + | 3.809 | x | | |
| 124.01857 | C9H3ON | [M-H2O+H]+ | + | 2.797 | x | | |
| 124.03698 | C4H7O2N | [M+Na]+ | + | 0.797 | x | | |
| 124.0548 | C10H7N | [M-H2O+H]+ | + | 1.694 | x | | |
| 124.07324 | C5H11ON | [M+Na]+ | + | -0.444 | x | | |
| 124.11217 | C8H15ON | [M-H2O+H]+ | + | 0.669 | x | | |
| 124.18795 | NA | NA | - | NA | | x | |
| 124.95329 | NA | NA | + | NA | x | | |
| 124.97374 | NA | NA | + | NA | x | | |
| 124.97708 | NA | NA | + | NA | x | | |
| 125.00248 | C9H2O2 | [M-H2O+H]+ | + | 2.034 | x | | |
| 125.01144 | C6H2N2 | [M+Na]+ | + | 4.124 | x | | |
| 125.0182 | NA | NA | + | NA | x | | |
| 125.02091 | C4H6O3 | [M+Na]+ | + | -0.050 | x | | |
| 125.93736 | NA | NA | + | NA | x | | |
| 125.95284 | C5H2O3S | [M-NH3+H]+ | + | -2.480 | x | | |
| 126.01408 | NA | NA | + | NA | x | | |
| 126.02734 | NA | NA | + | NA | x | | |
| 126.93684 | NA | NA | + | NA | x | | |
| 126.94983 | NA | NA | + | NA | x | | |
| 126.95146 | NA | NA | + | NA | x | | |
| 126.95612 | NA | NA | + | NA | x | | |

| | | | | | | | |
|---|---|---|---|---|---|---|---|
| 126.99817 | NA | NA | + | NA | x | | |
| 127.00019 | C9H4S | [M-H2O+H]+ | + | 0.745 | x | | |
| 127.00667 | NA | NA | + | NA | | x | |
| 127.01555 | C7H4O | [M+Na]+ | + | 1.097 | | x | x |
| 127.03446 | NA | NA | + | NA | x | | |
| 127.0365 | C4H8O3 | [M+Na]+ | + | -0.626 | x | | |
| 127.07285 | C5H12O2 | [M+Na]+ | + | -0.966 | x | | |
| 127.93543 | NA | NA | + | NA | x | | |
| 127.94016 | NA | NA | + | NA | x | | |
| 127.94971 | NA | NA | + | NA | x | | |
| 127.95699 | C8HOP | [M-NH3+H]+ | + | -1.658 | x | | |
| 127.99469 | C9H5P | [M+NH3-H]- | - | -0.145 | x | | |
| 128.02313 | C6H8O4 | [M-NH3+H]+ | + | 1.000 | x | | |
| 128.03191 | C3H7O3N | [M+Na]+ | + | 0.913 | x | | |
| 128.93369 | NA | NA | + | NA | x | | |
| 128.94104 | NA | NA | + | NA | x | | |
| 128.94671 | C4H2O2S2 | [M-H2O+H]+ | + | 2.586 | x | | |
| 128.95075 | NA | NA | + | NA | x | | |
| 128.953 | C7ONP | [M-NH3+H]+ | + | 3.603 | x | | |
| 128.96258 | NA | NA | + | NA | x | | |
| 128.97078 | NA | NA | + | NA | x | | |
| 128.98297 | NA | NA | - | NA | x | | |
| 129.05218 | C4H10O3 | [M+Na]+ | + | -0.331 | x | | |
| 129.13854 | C7H18ON2 | [M-H2O+H]+ | + | -0.579 | x | | |
| 129.93479 | NA | NA | + | NA | x | | |
| 129.93696 | NA | NA | + | NA | x | | |
| 129.94853 | C4H2O4S | [M-NH3+H]+ | + | 2.900 | x | | |
| 129.98116 | C8H2O3 | [M-NH3+H]+ | + | 0.264 | x | | |
| 130.0206 | C8H6NP | [M-H2O+H]+ | + | 0.697 | x | | |
| 130.05561 | NA | NA | + | NA | x | | |
| 130.09736 | C5H13O2N3 | [M-H2O+H]+ | + | -0.872 | x | | |
| 130.5439 | NA | NA | + | NA | x | | |
| 130.93248 | NA | NA | + | NA | x | | |
| 130.94557 | NA | NA | + | NA | x | | |
| 130.95179 | NA | NA | + | NA | x | | |
| 130.96507 | NA | NA | + | NA | x | | |
| 131.01906 | NA | NA | + | NA | x | | |
| 131.03145 | C9H8S | [M-H2O+H]+ | + | 0.455 | x | | |
| 131.55176 | NA | NA | + | NA | x | | |
| 131.93579 | NA | NA | + | NA | x | | |
| 131.94897 | NA | NA | + | NA | x | | |
| 131.95274 | C7HO2P | [M-NH3+H]+ | + | 4.033 | x | | |
| 131.99993 | C7H4ONP | [M-H2O+H]+ | + | 1.127 | x | | |
| 132.04281 | NA | NA | - | NA | | x | x |
| 132.55952 | NA | NA | + | NA | x | | |
| 132.90264 | NA | NA | + | NA | x | | |
| 132.93291 | NA | NA | + | NA | x | | |
| 132.98083 | C5H3OP | [M+Na]+ | + | -4.931 | x | | |
| 132.99824 | NA | NA | + | NA | x | | |
| 133.04619 | NA | NA | - | NA | | x | x |
| 133.06083 | C4H10O4N2 | [M-H2O+H]+ | + | 0.410 | x | | |
| 133.09716 | C5H14O3N2 | [M-H2O+H]+ | + | 0.041 | x | | |
| 133.93444 | NA | NA | + | NA | x | | |
| 133.97392 | NA | NA | + | NA | x | | |

| | | | | | | | | |
|---|---|---|---|---|---|---|---|---|
| 133.97399 | C3H6ON2S2 | [M+NH3-H]- | - | 0.077 | x | x | | |
| 133.97917 | C6H2O2NP | [M-H2O+H]+ | + | 0.949 | x | | | |
| 134.03919 | C11H5N | [M-H2O+H]+ | + | 1.847 | x | | | |
| 134.04464 | C4H10O4N2 | [M-NH3+H]+ | + | -0.960 | x | | | |
| 134.05401 | C6H14O2S | [M+NH3-H]- | - | 4.902 | | | x | |
| 134.05846 | NA | NA | - | NA | | | x | x |
| 134.91855 | NA | NA | + | NA | x | | | |
| 134.94193 | NA | NA | + | NA | x | | | |
| 134.95778 | NA | NA | + | NA | x | | | |
| 134.96119 | C4HO2P | [M+Na]+ | + | 4.939 | x | | | |
| 134.98679 | C10O2 | [M-H2O+H]+ | + | 1.637 | x | | | |
| 135.00504 | C5H4O3 | [M+Na]+ | + | -2.010 | x | | | |
| 135.0231 | C11H4O | [M-H2O+H]+ | + | 1.141 | x | | | |
| 135.12096 | NA | NA | + | NA | | | x | x |
| 135.93144 | C5N2S2 | [M-NH3+H]+ | + | 2.784 | x | | | |
| 135.94523 | NA | NA | + | NA | x | | | |
| 135.95623 | C7H4S2 | [M-NH3+H]+ | + | 0.401 | x | | | |
| 135.96907 | C4O2N3P | [M-H2O+H]+ | + | -2.972 | x | | | |
| 135.99013 | NA | NA | + | NA | x | | | |
| 136.03486 | NA | NA | + | NA | x | | | |
| 136.04626 | C9H12S | [M-NH3+H]+ | + | -2.882 | x | | | |
| 136.07326 | C6H11ON | [M+Na]+ | + | -0.220 | x | | | |
| 136.91537 | NA | NA | + | NA | x | | | |
| 136.9531 | C4O2N3P | [M-NH3+H]+ | + | -2.878 | x | | | |
| 136.9854 | NA | NA | + | NA | x | | | |
| 136.99109 | C4H10S3 | [M-H2O+H]+ | + | -0.546 | x | | | |
| 137.05735 | C6H10O2 | [M+Na]+ | + | 0.434 | x | | | |
| 137.9536 | C6H2O3S | [M-NH3+H]+ | + | 2.649 | x | | | |
| 137.9695 | NA | NA | + | NA | x | | | |
| 137.97097 | C6H2O5 | [M-NH3+H]+ | + | 0.094 | x | | | |
| 138.03409 | C10H5ON | [M-H2O+H]+ | + | 1.705 | x | | | |
| 138.04533 | C9H5N3 | [M-H2O+H]+ | + | 1.748 | x | | | |
| 138.05249 | C5H9O2N | [M+Na]+ | + | -0.517 | x | | | |
| 138.1124 | C5H15O3N | [M+H]+ | + | -0.506 | x | | | |
| 138.54119 | NA | NA | + | NA | x | | | |
| 138.95343 | NA | NA | + | NA | x | | | |
| 138.96367 | C9OS | [M-H2O+H]+ | + | -0.175 | x | | | |
| 138.99134 | NA | NA | + | NA | x | | | |
| 139.01804 | C10H4O2 | [M-H2O+H]+ | + | 1.274 | x | | | |
| 139.02931 | C9H4ON2 | [M-H2O+H]+ | + | 1.509 | x | | | |
| 139.03649 | C5H8O3 | [M+Na]+ | + | -0.647 | x | | | |
| 139.04218 | C4H12O4S | [M-H2O+H]+ | + | -1.032 | x | | | |
| 139.09472 | C9H16S | [M-H2O+H]+ | + | 4.723 | | | x | |
| 139.96486 | C9O3 | [M-NH3+H]+ | + | -3.920 | x | | | |
| 139.96834 | C8ONP | [M-H2O+H]+ | + | -0.778 | x | | | |
| 139.99461 | C9H3NS | [M-H2O+H]+ | + | -4.597 | x | | | |
| 140.0304 | NA | NA | + | NA | x | | | |
| 140.55694 | NA | NA | + | NA | x | | | |
| 140.94896 | NA | NA | + | NA | x | | | |
| 140.95238 | C8ONP | [M-NH3+H]+ | + | -0.622 | x | | | |
| 140.98296 | C5H3O5N | [M+NH3-H]- | - | 0.087 | x | | | |
| 140.98731 | NA | NA | + | NA | x | | | |
| 140.99555 | NA | NA | + | NA | x | | | |
| 141.00252 | NA | NA | - | NA | x | | | |

| | | | | | | |
|---|---|---|---|---|---|---|
| 141.0143 | NA | NA | + | NA | x | |
| 141.01572 | C10H6S | [M-H2O+H]+ | + | -0.080 | x | |
| 141.05215 | C5H10O3 | [M+Na]+ | + | -0.551 | x | |
| 141.08855 | C6H14O2 | [M+Na]+ | + | -0.428 | x | |
| 141.96529 | C5H2O6 | [M-NH3+H]+ | + | -3.641 | x | |
| 141.97974 | NA | NA | + | NA | x | |
| 142.96374 | NA | NA | + | NA | x | |
| 142.97932 | C7HON2P | [M-H2O+H]+ | + | -0.257 | x | |
| 142.9841 | NA | NA | + | NA | x | |
| 143.00143 | NA | NA | + | NA | x | |
| 143.01014 | C5ON6 | [M-H2O+H]+ | + | 0.434 | x | |
| 143.01359 | C4H8O2S | [M+Na]+ | + | -1.093 | x | |
| 143.03136 | C10H8S | [M-H2O+H]+ | + | -0.142 | x | |
| 143.06776 | C5H12O3 | [M+Na]+ | + | -0.875 | x | |
| 143.12595 | C9H20S | [M-H2O+H]+ | + | 4.167 | x | |
| 143.94777 | NA | NA | + | NA | x | |
| 143.96338 | C7O2NP | [M-H2O+H]+ | + | 0.021 | x | |
| 143.97794 | NA | NA | + | NA | x | |
| 143.98261 | C6H8OS2 | [M-NH3+H]+ | + | 1.415 | x | |
| 143.99384 | NA | NA | - | NA | x | |
| 144.94741 | C7O2NP | [M-NH3+H]+ | + | 0.110 | x | |
| 144.96034 | C5H6S3 | [M-H2O+H]+ | + | 2.877 | x | |
| 144.962 | NA | NA | + | NA | x | |
| 144.96659 | NA | NA | + | NA | x | |
| 144.97618 | C8H4P2 | [M-H2O+H]+ | + | 3.982 | x | |
| 144.992 | C8H2O4 | [M-H2O+H]+ | + | -0.126 | x | |
| 145.00871 | C7H6S | [M+Na]+ | + | 3.835 | x | |
| 145.02607 | C7H6O2 | [M+Na]+ | + | 0.570 | x | |
| 145.04705 | C10H10S | [M-H2O+H]+ | + | 0.107 | x | |
| 145.09706 | C6H14O3N2 | [M-H2O+H]+ | + | -0.579 | x | |
| 145.10829 | C5H14O2N4 | [M-H2O+H]+ | + | -0.600 | x | |
| 145.94594 | NA | NA | + | NA | x | |
| 145.95064 | NA | NA | + | NA | x | |
| 145.96027 | C8H4P2 | [M-NH3+H]+ | + | 4.441 | x | |
| 145.97725 | NA | NA | + | NA | x | |
| 146.09226 | C5H13O3N3 | [M-H2O+H]+ | + | -0.876 | x | |
| 146.94429 | NA | NA | + | NA | x | |
| 146.95154 | C5O2S | [M+Na]+ | + | 3.378 | x | |
| 146.96126 | NA | NA | + | NA | x | |
| 146.96388 | NA | NA | + | NA | x | |
| 146.97496 | C4H5O4NS | [M-NH3+H]+ | + | 1.867 | x | |
| 146.99637 | NA | NA | + | NA | x | |
| 147.07628 | C5H12O4N2 | [M-H2O+H]+ | + | -0.843 | x | |
| 147.9453 | NA | NA | + | NA | x | |
| 147.95904 | C5H8S3 | [M-NH3+H]+ | + | -3.047 | x | |
| 147.99166 | C8H4O4 | [M-NH3+H]+ | + | -0.159 | x | |
| 148.94311 | C6O3NP | [M-NH3+H]+ | + | 4.868 | x | |
| 148.9624 | C6H2N2S2 | [M-H2O+H]+ | + | -1.515 | x | |
| 148.97565 | NA | NA | + | NA | x | |
| 149.02102 | C6H6O3 | [M+Na]+ | + | 0.832 | x | |
| 149.04553 | C5H11O5N | [M+NH3-H]- | - | -0.099 | | x |
| 149.9464 | C6HONS2 | [M-H2O+H]+ | + | -1.599 | x | |
| 149.95945 | NA | NA | + | NA | x | |
| 149.9633 | C7H3O3P | [M-NH3+H]+ | + | 3.567 | x | |

| | | | | | | | | | |
|---|---|---|---|---|---|---|---|---|---|
| 149.99769 | C10HO2N | [M-H2O+H]+ | + | 1.496 | x | | | | |
| 150.03411 | C11H5ON | [M-H2O+H]+ | + | 1.703 | x | | | | |
| 150.04888 | C6H14O3S | [M+NH3-H]- | - | 4.162 | | | | x | |
| 150.06415 | C10H14S | [M+NH3-H]- | - | 4.244 | x | x | | | |
| 150.07048 | C12H9N | [M-H2O+H]+ | + | 1.610 | x | | | | |
| 150.91323 | C6S3 | [M-H2O+H]+ | + | 1.822 | x | | | | |
| 150.94344 | NA | NA | + | NA | x | | | | |
| 150.94723 | NA | NA | + | NA | x | | | | |
| 150.98168 | C10O3 | [M-H2O+H]+ | + | 1.335 | x | | | | |
| 150.98278 | C10HO2N | [M+NH3-H]- | - | 1.362 | | | | x | x |
| 150.9913 | C5H5O2P | [M+Na]+ | + | -4.976 | x | | | | |
| 151.00859 | NA | NA | + | NA | x | | | | |
| 151.0181 | C11H4O2 | [M-H2O+H]+ | + | 1.540 | x | | | | |
| 151.07296 | C7H12O2 | [M+Na]+ | + | 0.074 | x | | | | |
| 152.01338 | C10H3O2N | [M-H2O+H]+ | + | 1.715 | x | | | | |
| 152.03173 | C5H7O3N | [M+Na]+ | + | -0.652 | x | | | | |
| 152.04976 | C11H7ON | [M-H2O+H]+ | + | 1.682 | x | | | | |
| 152.0682 | C6H11O2N | [M+Na]+ | + | 0.004 | x | | | | |
| 152.95257 | C9ONP | [M-NH3+H]+ | + | 0.546 | x | | | | |
| 152.97182 | C4H3O3P | [M+Na]+ | + | 4.758 | x | | | | |
| 152.99735 | C10H2O3 | [M-H2O+H]+ | + | 1.436 | x | | | | |
| 153.01382 | NA | NA | + | NA | x | | | | |
| 153.01572 | C11H6S | [M-H2O+H]+ | + | -0.075 | x | | | | |
| 153.05021 | NA | NA | + | NA | x | | | | |
| 153.05213 | C6H10O3 | [M+Na]+ | + | -0.654 | x | | | | |
| 153.08861 | C7H14O2 | [M+Na]+ | + | 0.073 | x | | | | |
| 153.9558 | NA | NA | + | NA | x | | | | |
| 153.98399 | C9H2ONP | [M-H2O+H]+ | + | -0.714 | x | | | | |
| 154.02732 | C4H6ON5P | [M-H2O+H]+ | + | -2.269 | x | | | | |
| 154.04551 | NA | NA | + | NA | x | | | | |
| 154.04741 | C5H9O3N | [M+Na]+ | + | -0.413 | x | | | | |
| 154.05856 | C4H9O2N3 | [M+Na]+ | + | -1.049 | x | | | | |
| 154.96363 | C4HO4N2P | [M-H2O+H]+ | + | -2.762 | x | | | | |
| 154.96801 | C9HO2P | [M-H2O+H]+ | + | -0.684 | x | | | | |
| 154.98558 | C6O2N2 | [M+Na]+ | + | 2.890 | x | | | | |
| 155.01132 | C4H5O2N4P | [M-H2O+H]+ | + | -2.347 | x | | | | |
| 155.03142 | C11H8S | [M-H2O+H]+ | + | 0.217 | x | | | | |
| 155.15415 | C9H20ON2 | [M-H2O+H]+ | + | -0.724 | x | | | | |
| 155.9811 | C6H4O6 | [M-NH3+H]+ | + | -2.414 | x | | | | |
| 156.07666 | C6H11O3N3 | [M-H2O+H]+ | + | -0.536 | x | | | | |
| 156.97433 | C9H2O2S | [M-H2O+H]+ | + | 0.391 | x | | | | |
| 156.98073 | C7H3OP | [M+Na]+ | + | -4.794 | x | | | | |
| 156.99469 | C8H3ON2P | [M-H2O+H]+ | + | -1.845 | x | | | | |
| 157.01711 | C4H12S3 | [M+H]+ | + | -1.786 | x | | | | |
| 157.04709 | C5H10O4 | [M+Na]+ | + | -0.296 | x | | | | |
| 157.10518 | C9H18OS | [M-H2O+H]+ | + | 3.634 | x | | | | |
| 157.97913 | C8H2O2NP | [M-H2O+H]+ | + | 0.590 | x | | | | |
| 158.00951 | C5H6O5N2 | [M+NH3-H]- | - | 0.084 | x | | | | |
| 158.96306 | C8HO3P | [M-H2O+H]+ | + | 0.101 | x | | | | |
| 158.96991 | NA | NA | + | NA | x | | | | |
| 158.99637 | C5HN6P | [M-H2O+H]+ | + | -2.116 | x | | | | |
| 159.02591 | C10H8OS | [M-H2O+H]+ | + | -2.200 | x | | | | |
| 159.04161 | C8H8O2 | [M+Na]+ | + | -0.298 | x | | | | |
| 159.05011 | NA | NA | + | NA | x | | | | |

| | | | | | |
|---|---|---|---|---|---|
| 159.12387 | C6H16O2N4 | [M-H2O+H]+ | + | -0.949 | x |
| 159.97585 | C9H6P2 | [M-NH3+H]+ | + | 3.690 | x |
| 160.10794 | C6H15O3N3 | [M-H2O+H]+ | + | -0.637 | x |
| 160.95987 | NA | NA | + | NA | x |
| 160.97563 | C6H3O2P | [M+Na]+ | + | -4.761 | x |
| 161.02417 | C4H10O3S | [M+Na]+ | + | -0.839 | x |
| 161.0573 | C8H10O2 | [M+Na]+ | + | -0.004 | x |
| 161.06463 | C7H16OP2 | [M-H2O+H]+ | + | 1.575 | x |
| 161.97414 | C5H6O5S | [M-NH3+H]+ | + | -1.019 | x |
| 162.0302 | C7H5O3N3 | [M-H2O+H]+ | + | 2.219 | x |
| 162.95782 | C7HO4P | [M-H2O+H]+ | + | -0.761 | x |
| 162.97042 | C6H2O2N3P | [M+NH3-H]- | - | 0.743 | x |
| 162.99127 | C6H5O2P | [M+Na]+ | + | -4.764 | x |
| 163.00266 | C8H4O5 | [M-H2O+H]+ | + | 0.417 | x |
| 163.97074 | C8H6OP2 | [M-NH3+H]+ | + | 3.471 | x |
| 164.01334 | C11H3O2N | [M-H2O+H]+ | + | 1.380 | x |
| 164.95475 | C4H2N2S2 | [M+Na]+ | + | -2.893 | x |
| 164.99738 | C4H10N2S3 | [M-H2O+H]+ | + | 0.318 | x |
| 165.00697 | C6H7O2P | [M+Na]+ | + | -4.345 | x |
| 165.07565 | C6H14O6 | [M-H2O+H]+ | + | -0.547 | x |
| 165.08867 | C8H14O2 | [M+Na]+ | + | 0.489 | x |
| 165.95586 | NA | NA | + | NA | x |
| 166.08393 | C7H13O2N | [M+Na]+ | + | 0.563 | x |
| 166.95354 | C6HO5P | [M-H2O+H]+ | + | 3.634 | x |
| 166.9862 | C5H5O3P | [M+Na]+ | + | -4.525 | x |
| 167.02428 | C10H4O2N2 | [M-H2O+H]+ | + | 1.580 | x |
| 167.04 | C12H9P | [M-H2O+H]+ | + | -4.882 | x |
| 167.06782 | C7H12O3 | [M+Na]+ | + | -0.313 | x |
| 167.10423 | C8H16O2 | [M+Na]+ | + | -0.142 | x |
| 168.00823 | C10H3O3N | [M-H2O+H]+ | + | 1.217 | x |
| 168.06306 | C6H11O3N | [M+Na]+ | + | -0.373 | x |
| 168.95853 | C8ON3P | [M-NH3+H]+ | + | -0.517 | x |
| 168.99224 | C10H2O4 | [M-H2O+H]+ | + | 1.181 | x |
| 169.00184 | C5H7O3P | [M+Na]+ | + | -4.531 | x |
| 169.04707 | C12H10S | [M-H2O+H]+ | + | 0.200 | x |
| 169.05831 | C5H10O3N2 | [M+Na]+ | + | -0.364 | x |
| 169.11988 | C8H18O2 | [M+Na]+ | + | -0.140 | x |
| 169.97895 | C9H2O2NP | [M-H2O+H]+ | + | -0.410 | x |
| 170.00389 | C11H7OP | [M-NH3+H]+ | + | -1.552 | x |
| 170.02121 | C8H5O2N | [M+Na]+ | + | -0.269 | x |
| 170.04226 | C11H9NS | [M-H2O+H]+ | + | -0.116 | x |
| 170.0504 | C5H9ON5S | [M-H2O+H]+ | + | 4.853 | x |
| 170.11741 | C9H17O3N | [M-H2O+H]+ | + | -0.774 | x |
| 170.9631 | C9HO3P | [M-H2O+H]+ | + | 0.307 | x |
| 171.02626 | C11H8OS | [M-H2O+H]+ | + | -0.198 | x |
| 171.04181 | C9H8O2 | [M+Na]+ | + | 1.077 | x |
| 171.04561 | C6H12O2S | [M+Na]+ | + | 3.977 | x |
| 171.05391 | C4H4N5P | [M+NH4]+ | + | -2.268 | x |
| 171.11261 | C8H16O3N2 | [M-H2O+H]+ | + | -1.030 | x |
| 171.12079 | C10H20OS | [M-H2O+H]+ | + | 3.151 | x |
| 171.95845 | C8O3NP | [M-H2O+H]+ | + | 0.840 | x |
| 171.96269 | C13HP | [M-NH3+H]+ | + | 2.000 | x |
| 171.97619 | C6H4O7 | [M-NH3+H]+ | + | -1.276 | x |
| 172.02586 | C7H3O2N5 | [M-H2O+H]+ | + | 2.506 | x |

| | | | | | |
|---|---|---|---|---|---|
| 172.03786 | C4H8O2N5P | [M-H2O+H]+ | + | -2.183 | x |
| 172.13304 | C9H19O3N | [M-H2O+H]+ | + | -0.871 | x |
| 172.94238 | C8O3NP | [M-NH3+H]+ | + | 0.387 | x |
| 172.95219 | C4H6S3 | [M+Na]+ | + | -1.290 | x |
| 172.96014 | NA | NA | + | NA | x |
| 172.9962 | C6H2O3N2 | [M+Na]+ | + | 2.913 | x |
| 173.01683 | C9H6ON2S | [M-H2O+H]+ | + | 0.183 | x |
| 173.02194 | C4H7O3N4P | [M-H2O+H]+ | + | -1.833 | x |
| 173.03762 | C8H7N4P | [M-H2O+H]+ | + | 0.424 | x |
| 173.0785 | C6H14O4 | [M+Na]+ | + | 0.468 | x |
| 173.12838 | C8H18O3N2 | [M-H2O+H]+ | + | -0.388 | x |
| 173.13651 | C10H22OS | [M-H2O+H]+ | + | 3.485 | x |
| 173.95521 | C9H4OP2 | [M-NH3+H]+ | + | 3.921 | x |
| 173.97478 | C7H10S3 | [M-NH3+H]+ | + | -2.156 | x |
| 174.00087 | C9H5O2NS | [M-H2O+H]+ | + | 0.309 | x |
| 174.02159 | C8H6ON3P | [M-H2O+H]+ | + | 0.157 | x |
| 174.03487 | C8H9NS | [M+Na]+ | + | 0.523 | x |
| 174.08182 | NA | NA | + | NA | x |
| 174.93917 | C5N2S2 | [M+Na]+ | + | -2.242 | x |
| 174.95884 | C7O3N2S | [M-H2O+H]+ | + | -4.347 | x |
| 174.98488 | C9H4O3S | [M-H2O+H]+ | + | 0.279 | x |
| 174.99139 | C5HON6P | [M-H2O+H]+ | + | -1.391 | x |
| 175.00081 | C13H4S | [M-H2O+H]+ | + | 3.788 | x |
| 175.01873 | C8H8OS | [M+Na]+ | + | -0.504 | x |
| 175.02772 | C6H5N6P | [M-H2O+H]+ | + | -1.679 | x |
| 175.05771 | C5H12O5 | [M+Na]+ | + | 0.103 | x |
| 175.10764 | C7H16O4N2 | [M-H2O+H]+ | + | -0.408 | x |
| 175.11573 | C9H20O2S | [M-H2O+H]+ | + | 3.217 | x |
| 175.11889 | C6H16O3N4 | [M-H2O+H]+ | + | -0.322 | x |
| 175.93514 | C5H4O4S2 | [M-NH3+H]+ | + | -3.581 | x |
| 175.98442 | C13H4S | [M-NH3+H]+ | + | 1.675 | x |
| 175.98996 | C6H8O5S | [M-NH3+H]+ | + | -0.059 | x |
| 175.99646 | NA | NA | + | NA | x |
| 176.02299 | C10H8O4 | [M-NH3+H]+ | + | 0.021 | x |
| 176.0998 | C9H22P2 | [M-NH3+H]+ | + | -3.126 | x |
| 176.10286 | C6H15O4N3 | [M-H2O+H]+ | + | -0.556 | x |
| 176.91913 | NA | NA | + | NA | x |
| 176.93799 | C7O4NP | [M-NH3+H]+ | + | 3.983 | x |
| 176.97407 | C8H3O4P | [M-H2O+H]+ | + | 2.387 | x |
| 176.98049 | C7H6S2 | [M+Na]+ | + | 1.151 | x |
| 176.9844 | C4H2ON4S | [M+Na]+ | + | 1.607 | x |
| 176.98902 | C12H3OP | [M-H2O+H]+ | + | 0.809 | x |
| 176.99599 | C6H3O2N4P | [M-H2O+H]+ | + | -0.431 | x |
| 176.99821 | C7H6O2S | [M+Na]+ | + | 0.901 | x |
| 177.00697 | C5H3ON6P | [M-H2O+H]+ | + | -1.738 | x |
| 177.01918 | C4H10O4S | [M+Na]+ | + | -0.132 | x |
| 177.02433 | C13H7P | [M-H2O+H]+ | + | -4.734 | x |
| 177.06072 | NA | NA | + | NA | x |
| 177.08867 | C9H14O2 | [M+Na]+ | + | 0.451 | x |
| 177.12329 | C7H18O4N2 | [M-H2O+H]+ | + | -0.404 | x |
| 177.93202 | NA | NA | + | NA | x |
| 177.98012 | C6H2O3N3P | [M-H2O+H]+ | + | 0.157 | x |
| 177.99269 | C4H9N3S3 | [M-H2O+H]+ | + | 0.610 | x |
| 178.01038 | C4H9O2N3S2 | [M-H2O+H]+ | + | 0.258 | x |

| | | | | | | | | | |
|---|---|---|---|---|---|---|---|---|---|
| 178.91603 | NA | NA | + | NA | x | | | | |
| 178.97696 | C8HN2P | [M+Na]+ | + | 0.027 | x | | | | |
| 178.98628 | C6H5O3P | [M+Na]+ | + | -3.664 | x | | | | |
| 179.01307 | C5H12N2S3 | [M-H2O+H]+ | + | 0.499 | x | | | | |
| 179.04935 | C13H8O2 | [M-H2O+H]+ | + | 1.065 | x | | | | |
| 179.09132 | C7H16O6 | [M-H2O+H]+ | + | -0.406 | x | | | | |
| 179.10422 | C9H16O2 | [M+Na]+ | + | -0.195 | | | | | x |
| 179.93085 | C5H8S4 | [M-NH3+H]+ | + | -3.881 | x | | | | |
| 180.09328 | C10H16ONP | [M-H2O+H]+ | + | -1.939 | | | | x | x |
| 180.09458 | C8H20O3S | [M-NH3+H]+ | + | 2.743 | x | | | | |
| 180.09947 | C8H15O2N | [M+Na]+ | + | -0.188 | x | | | | |
| 180.9475 | C10O2NP | [M-NH3+H]+ | + | 0.547 | x | | | | |
| 180.98367 | C11H3O2P | [M-H2O+H]+ | + | -0.544 | x | | | | |
| 180.99984 | C6H6N4S2 | [M-H2O+H]+ | + | -1.310 | x | | | | |
| 181.00192 | C6H7O3P | [M+Na]+ | + | -3.681 | x | | | | |
| 181.00863 | C10H6S | [M+Na]+ | + | 2.455 | x | | | | |
| 181.01083 | C6H6O5 | [M+Na]+ | + | 0.542 | x | | | | |
| 181.03828 | C5H7ON6P | [M-H2O+H]+ | + | -1.652 | x | | | | |
| 181.04714 | C7H10O4 | [M+Na]+ | + | 0.065 | x | | | | |
| 181.12001 | C9H18O2 | [M+Na]+ | + | 0.692 | x | | | | |
| 182.024 | C4H13N3S3 | [M-H2O+H]+ | + | 0.648 | x | | | | |
| 182.08132 | C9H13O4N | [M-H2O+H]+ | + | 0.756 | x | | | | |
| 182.08435 | C8H14ON3P | [M-H2O+H]+ | + | 0.954 | x | | | | |
| 182.11518 | C8H17O2N | [M+Na]+ | + | 0.192 | x | | | | |
| 182.90234 | C6O2S3 | [M-H2O+H]+ | + | -2.067 | x | | | | |
| 182.95573 | C6HO7N | [M-NH3+H]+ | + | -1.502 | x | | | | |
| 182.96304 | C10HO3P | [M-H2O+H]+ | + | -0.012 | x | | | | |
| 182.98071 | C7O3N2 | [M+Na]+ | + | 3.731 | x | | | | |
| 183.01764 | C4ON5P | [M+NH4]+ | + | -1.404 | x | | | | |
| 183.07806 | C6H17O5P | [M-H2O+H]+ | + | -0.057 | x | | | | |
| 183.97914 | C6H4O5NP | [M-H2O+H]+ | + | -1.396 | x | | | | |
| 183.98151 | C13ON2 | [M-NH3+H]+ | + | -1.401 | x | | | | |
| 183.99468 | C10H4O2NP | [M-H2O+H]+ | + | 0.016 | x | | | | |
| 184.95999 | NA | NA | + | NA | x | | | | |
| 184.96929 | C10H2O3S | [M-H2O+H]+ | + | 0.562 | x | | | | |
| 185.02113 | C9H6O3 | [M+Na]+ | + | 1.326 | x | | | | |
| 185.04204 | C6H10O5 | [M+Na]+ | + | -0.027 | x | | | | |
| 185.05728 | C10H10O2 | [M+Na]+ | + | -0.126 | x | | | | |
| 185.98521 | C8H2O2N3P | [M-H2O+H]+ | + | 0.174 | x | | | | |
| 186.01047 | C10H6O2NP | [M-H2O+H]+ | + | 0.706 | x | | | | |
| 186.93562 | NA | NA | + | NA | x | | | | |
| 186.95869 | C9HO4P | [M-H2O+H]+ | + | 3.594 | x | | | | |
| 186.9944 | C10H5O3P | [M-H2O+H]+ | + | 0.283 | x | | | | |
| 187.02782 | C5H14OS3 | [M+H]+ | + | -0.717 | x | | | | |
| 187.05766 | C6H12O5 | [M+Na]+ | + | -0.209 | x | | | | |
| 187.12031 | C10H16N2 | [M+Na]+ | + | -1.580 | x | | | | |
| 187.14408 | C9H20O3N2 | [M-H2O+H]+ | + | -0.117 | x | | | | |
| 187.98205 | C9H5ONP2 | [M-H2O+H]+ | + | 3.422 | x | | | | |
| 187.98997 | C7H8O5S | [M-NH3+H]+ | + | -0.006 | x | | | | |
| 188.12809 | C9H19O4N | [M-H2O+H]+ | + | -0.143 | x | | | | |
| 188.96606 | C9H4O2P2 | [M-H2O+H]+ | + | 3.378 | x | | | | |
| 188.97399 | C9H3O4P | [M-H2O+H]+ | + | 1.860 | x | | | | |
| 189.00701 | C4H12O2S3 | [M+H]+ | + | -1.106 | x | | | | |
| 189.05236 | C9H10O3 | [M+Na]+ | + | 0.903 | x | | | | |

| | | | | | | | | |
|---|---|---|---|---|---|---|---|---|
| 189.07333 | C6H14O5 | [M+Na]+ | + | -0.086 | x | | | |
| 189.11212 | C9H18O5 | [M-H2O+H]+ | + | -0.073 | x | | | |
| 189.12325 | C8H18O4N2 | [M-H2O+H]+ | + | -0.574 | x | | | |
| 189.13143 | C10H24P2 | [M-H2O+H]+ | + | -2.935 | x | | | |
| 189.15975 | C9H22O3N2 | [M-H2O+H]+ | + | -0.018 | x | | | |
| 189.97341 | C13H3OP | [M-NH3+H]+ | + | 2.579 | x | | | |
| 189.98088 | C13H2O3 | [M-NH3+H]+ | + | -1.172 | x | | | |
| 190.11546 | C10H24P2 | [M-NH3+H]+ | + | -2.865 | x | | | |
| 190.13241 | C10H22O4 | [M-NH3+H]+ | + | -0.611 | x | | | |
| 190.13497 | C9H22O2NP | [M-H2O+H]+ | + | -2.688 | x | | | |
| 190.97069 | C6H4N2S2 | [M+Na]+ | + | -0.719 | x | | | |
| 191.99899 | C12H4ONP | [M-H2O+H]+ | + | -3.694 | x | | | |
| 192.96165 | C4H2N4S2 | [M+Na]+ | + | 2.008 | x | | | |
| 192.97936 | C4H2O2N4S | [M+Na]+ | + | 1.722 | x | | | |
| 192.99238 | C5H10ON2S3 | [M-H2O+H]+ | + | 0.682 | x | | | |
| 193.00187 | C5H3O2N6P | [M-H2O+H]+ | + | -1.675 | x | | | |
| 193.02317 | C7H2N6 | [M+Na]+ | + | -0.855 | x | | | |
| 193.03825 | C6H7ON6P | [M-H2O+H]+ | + | -1.700 | x | | | |
| 193.04713 | C8H10O4 | [M+Na]+ | + | -0.028 | x | | | |
| 193.10706 | C8H18O6 | [M-H2O+H]+ | + | 0.049 | x | | | |
| 193.10826 | C8H19O5N | [M+NH3-H]- | - | 0.543 | | | x | x |
| 193.11992 | C10H18O2 | [M+Na]+ | + | 0.115 | x | | | |
| 193.12636 | C9H24OP2 | [M-H2O+H]+ | + | -2.806 | x | | | |
| 193.13104 | C9H18ON2 | [M+Na]+ | + | -0.552 | x | | | |
| 193.15459 | C8H22O4N2 | [M-H2O+H]+ | + | -0.373 | x | | | |
| 193.9457 | C4HON3S2 | [M+Na]+ | + | 2.198 | x | | | |
| 193.9951 | C5H10O3N2S2 | [M+NH3-H]- | - | -0.036 | x | | | |
| 194.0071 | C7HON5 | [M+Na]+ | + | -1.350 | x | | | |
| 194.07871 | C8H13O3N | [M+Na]+ | + | -0.316 | x | | | |
| 194.11509 | C9H17O2N | [M+Na]+ | + | -0.348 | x | | | |
| 194.92974 | C4O2N2S2 | [M+Na]+ | + | 2.327 | x | | | |
| 194.95855 | C6HO5N2P | [M-H2O+H]+ | + | -2.215 | x | | | |
| 194.96037 | C6H2O4N3P | [M+NH3-H]- | - | 1.202 | x | | | |
| 194.96611 | C10H2N2P2 | [M-H2O+H]+ | + | 0.363 | x | | | |
| 194.99105 | C12H6P2 | [M-H2O+H]+ | + | -0.637 | x | | | |
| 194.99954 | C12H5O2P | [M-H2O+H]+ | + | 0.530 | x | | | |
| 195.03778 | C6H8O4N2 | [M+Na]+ | + | 0.885 | x | | | |
| 195.06274 | C8H12O4 | [M+Na]+ | + | -0.231 | x | | | |
| 195.11402 | C10H17O2N3 | [M+NH3-H]- | - | 0.566 | x | x | | |
| 195.11851 | C10H20S | [M+Na]+ | + | 4.171 | x | | | |
| 195.9426 | C6O6NP | [M-H2O+H]+ | + | -2.043 | x | | | |
| 196.00313 | C4H11ON3S3 | [M-H2O+H]+ | + | -0.027 | x | | | |
| 196.02168 | C6H7O5N | [M+Na]+ | + | 0.212 | x | | | |
| 196.07336 | C6H16O5NP | [M-H2O+H]+ | + | 0.185 | x | | | |
| 196.09454 | C8H15O3N | [M+Na]+ | + | 0.727 | x | | | |
| 196.11783 | C7H19O6N | [M-H2O+H]+ | + | -0.556 | x | | | |
| 196.126 | C10H19N3S | [M-H2O+H]+ | + | -3.189 | x | | | |
| 196.92655 | C6O6NP | [M-NH3+H]+ | + | -2.351 | x | | | |
| 196.9396 | C5O5N3P | [M+NH3-H]- | - | 1.029 | x | | | |
| 196.9544 | C8H6OS3 | [M-H2O+H]+ | + | -1.817 | x | | | |
| 196.98726 | C4H10O2N2S3 | [M-H2O+H]+ | + | 0.508 | x | | | |
| 197.01206 | C6H14O2S3 | [M-H2O+H]+ | + | -1.137 | x | | | |
| 197.03332 | C5H7O2N6P | [M-H2O+H]+ | + | -0.942 | x | | | |
| 197.05735 | C6H15O6P | [M-H2O+H]+ | + | 0.065 | x | | | |

| | | | | | | | |
|---|---|---|---|---|---|---|---|
| 197.09769 | C9H18OS | [M+Na]+ | + | 3.638 | x | | |
| 197.12127 | C8H22OP2 | [M+H]+ | + | -3.030 | x | | |
| 197.94143 | C5H10OS4 | [M-NH3+H]+ | + | -3.482 | x | | |
| 197.97386 | C10H2O3NP | [M-H2O+H]+ | + | -0.378 | x | | |
| 198.01026 | C11H6O2NP | [M-H2O+H]+ | + | -0.310 | x | | |
| 198.01746 | C5H6O3N5P | [M-H2O+H]+ | + | -0.360 | x | | |
| 198.95799 | C10HO4P | [M-H2O+H]+ | + | 0.153 | x | | |
| 198.99427 | C11H5O3P | [M-H2O+H]+ | + | -0.335 | x | | |
| 199.15213 | C12H24OS | [M-H2O+H]+ | + | 2.927 | x | | |
| 199.97111 | C7H4O8 | [M-NH3+H]+ | + | -1.085 | x | | |
| 199.9896 | C10H4O3NP | [M-H2O+H]+ | + | 0.040 | x | | |
| 200.00712 | C7H3O3N3 | [M+Na]+ | + | 2.587 | x | | |
| 200.1141 | C7H15O3N5 | [M-H2O+H]+ | + | -0.464 | x | | |
| 200.9129 | C6H2O3S3 | [M-H2O+H]+ | + | -1.918 | x | | |
| 200.95523 | C7H2N2S2 | [M+Na]+ | + | 0.389 | x | | |
| 200.97351 | C10H3O4P | [M-H2O+H]+ | + | -0.445 | x | | |
| 200.99113 | C13H2N2S | [M-H2O+H]+ | + | 2.520 | x | | |
| 201.04333 | C6H16OS3 | [M+H]+ | + | -1.366 | x | | |
| 201.07323 | C13H14OS | [M-H2O+H]+ | + | -0.079 | x | | |
| 201.16779 | C12H26OS | [M-H2O+H]+ | + | 2.946 | x | | |
| 201.97698 | C9HO6N | [M-H2O+H]+ | + | -0.542 | x | | |
| 201.98434 | C11H6O3S | [M-NH3+H]+ | + | -0.698 | x | | |
| 202.00526 | C10H6O3NP | [M-H2O+H]+ | + | 0.085 | x | | |
| 202.97039 | C7H4N2S2 | [M+Na]+ | + | -2.337 | x | | |
| 202.98952 | C10H5O4P | [M-H2O+H]+ | + | 1.196 | x | | |
| 203.02262 | C5H14O2S3 | [M+H]+ | + | -1.227 | x | | |
| 203.0525 | C12H12O2S | [M-H2O+H]+ | + | -0.054 | x | | |
| 203.05902 | C8H9N6P | [M-H2O+H]+ | + | -1.465 | x | | |
| 203.13897 | C9H20O4N2 | [M-H2O+H]+ | + | -0.220 | x | | |
| 203.14708 | C11H26P2 | [M-H2O+H]+ | + | -2.748 | x | | |
| 203.97152 | C12O3N2 | [M-NH3+H]+ | + | -0.452 | x | | |
| 204.00205 | C12H3N3S | [M-H2O+H]+ | + | 2.580 | x | | |
| 204.0176 | C11H8O5 | [M-NH3+H]+ | + | -1.366 | x | | |
| 204.96918 | C9H3O5P | [M-H2O+H]+ | + | 2.967 | x | | |
| 204.9861 | C7H6N2S2 | [M+Na]+ | + | -1.982 | x | | |
| 205.00211 | C4H12O3S3 | [M+H]+ | + | -0.110 | x | | |
| 205.03827 | C7H7ON6P | [M-H2O+H]+ | + | -1.518 | x | | |
| 205.06821 | C12H14O2S | [M-H2O+H]+ | + | 0.217 | x | | |
| 205.11822 | C8H18O5N2 | [M-H2O+H]+ | + | -0.284 | x | | |
| 205.15464 | C9H22O4N2 | [M-H2O+H]+ | + | -0.128 | x | | |
| 206.00087 | C7H10O6S | [M-NH3+H]+ | + | 1.505 | x | | |
| 206.13859 | C9H21O5N | [M-H2O+H]+ | + | -0.421 | x | | |
| 206.96037 | C5O6S | [M-H2O-H]- | - | -0.672 | x | | |
| 206.98477 | C9H5O5P | [M-H2O+H]+ | + | 2.672 | x | | |
| 207.01752 | C6H5O2N6P | [M-H2O+H]+ | + | -1.570 | x | | |
| 207.04434 | C7H16N2S3 | [M-H2O+H]+ | + | 0.303 | x | | |
| 207.09914 | C10H16O3 | [M+Na]+ | + | -0.136 | x | | |
| 207.142 | C10H26OP2 | [M-H2O+H]+ | + | -2.675 | x | | |
| 208.09428 | C7H11O2N7 | [M-H2O+H]+ | + | 0.648 | x | | |
| 208.13076 | C10H19O2N | [M+Na]+ | + | -0.213 | x | | |
| 209.03321 | C6H7O2N6P | [M-H2O+H]+ | + | -1.379 | x | | |
| 209.07829 | C15H14S | [M-H2O+H]+ | + | -0.189 | x | | |
| 209.10187 | C8H18O7 | [M-H2O+H]+ | + | -0.417 | x | | |
| 209.10321 | C8H19O6N | [M+NH3-H]- | - | 0.662 | | | x |

| | | | | | |
|---|---|---|---|---|---|
| 209.12064 | C9H22O4S | [M-H2O+H]+ | + | 0.216 | x |
| 209.12594 | C9H18O2N2 | [M+Na]+ | + | -0.583 | x |
| 210.08895 | C12H13ON | [M+Na]+ | + | 0.081 | x |
| 210.11002 | C9H17O3N | [M+Na]+ | + | -0.236 | x |
| 211.073 | C7H17O6P | [M-H2O+H]+ | + | 0.061 | x |
| 211.09407 | C9H16O4 | [M+Na]+ | + | -0.052 | x |
| 211.13041 | C10H20O3 | [M+Na]+ | + | -0.293 | x |
| 211.95627 | C9HON3P2 | [M-H2O+H]+ | + | 0.321 | x |
| 211.98959 | C11H4O3NP | [M-H2O+H]+ | + | -0.006 | x |
| 212.06829 | C6H16O6NP | [M-H2O+H]+ | + | 0.240 | x |
| 212.08928 | C14H15NS | [M-H2O+H]+ | + | 0.211 | x |
| 212.09737 | C8H15ON5S | [M-H2O+H]+ | + | 4.049 | x |
| 212.12564 | C9H19O3N | [M+Na]+ | + | -0.392 | x |
| 212.94028 | C9O2N2P2 | [M-H2O+H]+ | + | 0.295 | x |
| 212.97358 | C11H3O4P | [M-H2O+H]+ | + | -0.117 | x |
| 213.01004 | C12H7O3P | [M-H2O+H]+ | + | 0.207 | x |
| 213.15258 | C10H22N4S | [M-H2O+H]+ | + | -2.819 | x |
| 213.95316 | C9HON3S2 | [M-H2O+H]+ | + | 1.494 | x |
| 214.12849 | C7H19O6N | [M+H]+ | + | -0.109 | x |
| 214.93712 | C9O2N2S2 | [M-H2O+H]+ | + | 1.248 | x |
| 214.98924 | C11H5O4P | [M-H2O+H]+ | + | -0.073 | x |
| 215.05903 | C7H18OS3 | [M+H]+ | + | -1.043 | x |
| 215.1754 | C11H24O3N2 | [M-H2O+H]+ | + | -0.016 | x |
| 215.98454 | C10H4O4NP | [M-H2O+H]+ | + | 0.146 | x |
| 216.15936 | C11H23O4N | [M-H2O+H]+ | + | -0.255 | x |
| 216.93602 | C11P2 | [M+Na]+ | + | -3.735 | x |
| 216.96854 | C10H3O5P | [M-H2O+H]+ | + | 0.079 | x |
| 216.98613 | C8H6N2S2 | [M+Na]+ | + | -1.705 | x |
| 216.99531 | C11H6O4S | [M-H2O+H]+ | + | -0.347 | x |
| 217.0054 | C8H8N2P2 | [M+Na]+ | + | -0.476 | x |
| 217.03831 | C6H16O2S3 | [M+H]+ | + | -0.962 | x |
| 217.04723 | C10H10O4 | [M+Na]+ | + | 0.517 | x |
| 217.06818 | C13H14O2S | [M-H2O+H]+ | + | 0.077 | x |
| 217.10461 | C8H18O5 | [M+Na]+ | + | -0.177 | x |
| 217.15619 | C13H22O | [M+Na]+ | + | -0.494 | x |
| 217.16271 | C12H28P2 | [M-H2O+H]+ | + | -2.669 | x |
| 217.96158 | C11H6O2S2 | [M-NH3+H]+ | + | -0.291 | x |
| 217.9816 | C7H5N3S2 | [M+Na]+ | + | -0.563 | x |
| 218.10781 | C15H13N3 | [M-H2O+H]+ | + | 0.642 | x |
| 218.12462 | C7H17O4N5 | [M-H2O+H]+ | + | -0.618 | x |
| 218.96569 | C7H4ON2S2 | [M+Na]+ | + | -0.180 | x |
| 218.98495 | C7H6ON2P2 | [M+Na]+ | + | 0.985 | x |
| 219.01758 | C5H14O3S3 | [M+H]+ | + | -0.929 | x |
| 219.02653 | C9H8O5 | [M+Na]+ | + | 0.692 | x |
| 219.05388 | C8H9ON6P | [M-H2O+H]+ | + | -1.597 | x |
| 219.08383 | C13H16O2S | [M-H2O+H]+ | + | 0.077 | x |
| 219.09033 | C9H13N6P | [M-H2O+H]+ | + | -1.324 | x |
| 219.12261 | C10H20O6 | [M-H2O+H]+ | + | -0.379 | x |
| 219.12816 | C16H16N2 | [M-H2O+H]+ | + | 0.423 | x |
| 219.98001 | C7H8O7S | [M-NH3+H]+ | + | 0.888 | x |
| 219.99488 | C11H8O4S | [M-NH3+H]+ | + | -0.749 | x |
| 220.9635 | C9H3O6P | [M-H2O+H]+ | + | 0.269 | x |
| 221.03317 | C7H7O2N6P | [M-H2O+H]+ | + | -1.478 | x |
| 221.0696 | C8H11ON6P | [M-H2O+H]+ | + | -1.290 | x |

| | | | | | | | |
|---|---|---|---|---|---|---|---|
| 221.14975 | C9H22O5N2 | [M-H2O+H]+ | + | 0.701 | x | | |
| 222.14634 | C11H21O2N | [M+Na]+ | + | -0.550 | x | | |
| 222.96128 | C14S | [M+Na]+ | + | -0.060 | x | | |
| 222.99464 | C13H5O3P | [M-H2O+H]+ | + | 1.240 | x | | |
| 223.02773 | C8H16O2S3 | [M-H2O+H]+ | + | -0.930 | x | | |
| 223.0938 | C8H12O3N6 | [M-H2O+H]+ | + | 0.001 | x | | |
| 223.10504 | C7H12O2N8 | [M-H2O+H]+ | + | 0.028 | x | | |
| 223.11748 | C9H20O7 | [M-H2O+H]+ | + | -0.559 | x | | |
| 223.13037 | C11H20O3 | [M+Na]+ | + | -0.475 | x | | |
| 224.02589 | C13H8O2NP | [M-H2O+H]+ | + | -0.360 | x | | |
| 224.12557 | C8H15O2N7 | [M-H2O+H]+ | + | 0.563 | x | | |
| 225.00049 | C13H6O3S | [M-H2O+H]+ | + | 0.056 | x | | |
| 225.00997 | C13H7O3P | [M-H2O+H]+ | + | -0.092 | x | | |
| 225.05232 | C7H15O7P | [M-H2O+H]+ | + | 0.286 | x | | |
| 225.10954 | C8H14O3N6 | [M-H2O+H]+ | + | 0.373 | x | | |
| 225.95099 | C5H2O6NP | [M+Na]+ | + | -1.007 | x | | |
| 226.00522 | C12H6O3NP | [M-H2O+H]+ | + | -0.088 | x | | |
| 226.02283 | C10H10O7 | [M-NH3+H]+ | + | -2.272 | x | | |
| 226.0416 | C13H10O2NP | [M-H2O+H]+ | + | -0.110 | x | | |
| 226.05049 | C10H10N3P | [M+Na]+ | + | 0.173 | x | | |
| 226.0749 | C15H14O3 | [M-NH3+H]+ | + | -0.501 | x | | |
| 226.08389 | C7H18O6NP | [M-H2O+H]+ | + | 0.020 | x | | |
| 226.10476 | C7H13O3N7 | [M-H2O+H]+ | + | 0.252 | x | | |
| 226.12835 | C8H21O7N | [M-H2O+H]+ | + | -0.671 | x | | |
| 226.14377 | C12H21O4N | [M-H2O+H]+ | + | 0.002 | x | | |
| 226.98934 | C12H5O4P | [M-H2O+H]+ | + | 0.341 | x | | |
| 227.02563 | C13H9O3P | [M-H2O+H]+ | + | -0.050 | x | | |
| 227.03453 | C6H8O3N6S | [M-H2O+H]+ | + | -0.166 | x | | |
| 227.05896 | C8H20O2S3 | [M-H2O+H]+ | + | -1.202 | x | | |
| 227.06793 | C7H17O7P | [M-H2O+H]+ | + | 0.120 | x | | |
| 227.0889 | C15H16OS | [M-H2O+H]+ | + | 0.011 | x | | |
| 227.10821 | C10H20O2S | [M+Na]+ | + | 2.885 | x | | |
| 227.9479 | C10O5NP | [M-H2O+H]+ | + | -0.900 | x | | |
| 228.00208 | C9H8O8 | [M-NH3+H]+ | + | -2.313 | x | | |
| 228.07805 | C10H16O4NP | [M-H2O+H]+ | + | -1.452 | | | x |
| 228.08194 | C12H15NS | [M+Na]+ | + | 0.946 | x | | |
| 228.14417 | C8H21O6N | [M+H]+ | + | 0.030 | x | | |
| 228.14946 | C14H19ON3 | [M-H2O+H]+ | + | -0.259 | x | | |
| 228.99542 | C12H6O4S | [M-H2O+H]+ | + | 0.117 | x | | |
| 229.02248 | C15H6N2S | [M-H2O+H]+ | + | 2.436 | x | | |
| 229.03833 | C7H18O3S3 | [M-H2O+H]+ | + | -0.764 | x | | |
| 229.07464 | C8H20OS3 | [M+H]+ | + | -1.154 | x | | |
| 229.10455 | C15H18OS | [M-H2O+H]+ | + | 0.011 | x | | |
| 229.15458 | C11H22O4N2 | [M-H2O+H]+ | + | -0.359 | x | | |
| 230.00901 | C8H6O2N3P | [M+Na]+ | + | 0.126 | x | | |
| 230.14989 | C10H21O4N3 | [M-H2O+H]+ | + | -0.111 | x | | |
| 230.99304 | C8H5O3N2P | [M+Na]+ | + | 0.194 | x | | |
| 231.01761 | C6H16O4S3 | [M-H2O+H]+ | + | -0.696 | x | | |
| 231.05396 | C7H18O2S3 | [M+H]+ | + | -0.904 | x | | |
| 231.08374 | C14H16O2S | [M-H2O+H]+ | + | -0.290 | x | | |
| 231.17031 | C11H24O4N2 | [M-H2O+H]+ | + | -0.034 | x | | |
| 231.93742 | C14O3S | [M-NH3+H]+ | + | -0.492 | x | | |
| 233.03322 | C6H16O3S3 | [M+H]+ | + | -0.916 | x | | |
| 233.06087 | C16H11O2N | [M+NH3-H]- | - | 0.271 | x | x | |

| | | | | | | |
|---|---|---|---|---|---|---|
| 233.06957 | C9H11ON6P | [M-H2O+H]+ | + | -1.348 | x | |
| 233.07855 | C6H17O7P | [M+H]+ | + | 0.365 | x | |
| 233.09949 | C14H18O2S | [M-H2O+H]+ | + | 0.112 | x | |
| 233.10583 | C10H15N6P | [M-H2O+H]+ | + | -1.849 | x | |
| 233.93388 | C6H5O4NS3 | [M-H2O+H]+ | + | -3.593 | x | |
| 233.96759 | C6H2O4N3P | [M+Na]+ | + | 0.364 | x | |
| 234.10838 | C8H17O6N3 | [M-H2O+H]+ | + | -0.245 | x | |
| 234.14637 | C12H21O2N | [M+Na]+ | + | -0.376 | x | |
| 234.91819 | C5N4S3 | [M+Na]+ | + | 2.173 | x | |
| 234.9516 | C6HO5N2P | [M+Na]+ | + | 0.336 | x | |
| 235.04882 | C8H9O2N6P | [M-H2O+H]+ | + | -1.395 | x | |
| 235.07567 | C9H20N2S3 | [M-H2O+H]+ | + | 0.388 | x | x |
| 235.1176 | C10H20O7 | [M-H2O+H]+ | + | -0.056 | x | |
| 235.93003 | C8HO5NP2 | [M-H2O+H]+ | + | 1.277 | x | |
| 235.97215 | C11H8O3S2 | [M-NH3+H]+ | + | -0.249 | x | |
| 236.12566 | C11H19O3N | [M+Na]+ | + | -0.254 | x | |
| 236.16203 | C12H23O2N | [M+Na]+ | + | -0.326 | x | |
| 237.02808 | C7H7O3N6P | [M-H2O+H]+ | + | -1.383 | x | |
| 237.10096 | C9H15ON6P | [M-H2O+H]+ | + | -0.972 | x | |
| 237.12063 | C8H14O2N8 | [M-H2O+H]+ | + | -0.209 | x | |
| 237.1456 | C10H26ON2S2 | [M-H2O+H]+ | + | 0.921 | x | |
| 237.18113 | C10H26O5N2 | [M-H2O+H]+ | + | 0.991 | x | |
| 238.00553 | C11H10O5S | [M-NH3+H]+ | + | -0.359 | x | |
| 238.10487 | C16H17NS | [M-H2O+H]+ | + | -0.046 | x | |
| 238.14119 | C9H17O2N7 | [M-H2O+H]+ | + | 0.415 | x | |
| 239.12509 | C9H16O3N6 | [M-H2O+H]+ | + | -0.038 | x | |
| 239.1488 | C10H24O7 | [M-H2O+H]+ | + | -0.446 | x | |
| 239.16153 | C10H20O2N6 | [M-H2O+H]+ | + | 0.175 | x | |
| 240.0209 | C13H8O3NP | [M-H2O+H]+ | + | 0.034 | x | |
| 240.05425 | C15H12O4 | [M-NH3+H]+ | + | -0.141 | x | |
| 240.09227 | C17H11N3 | [M-H2O+H]+ | + | 1.015 | x | |
| 240.12049 | C16H19NS | [M-H2O+H]+ | + | -0.162 | x | |
| 240.14401 | C9H23O7N | [M-H2O+H]+ | + | -0.596 | x | |
| 240.15215 | C12H23ON3S | [M-H2O+H]+ | + | -2.894 | x | |
| 241.03827 | C8H18O3S3 | [M-H2O+H]+ | + | -0.961 | x | |
| 241.06817 | C15H14O2S | [M-H2O+H]+ | + | 0.031 | x | |
| 241.09204 | C8H10N8 | [M+Na]+ | + | -0.107 | x | |
| 241.10446 | C16H18OS | [M-H2O+H]+ | + | -0.338 | x | |
| 241.14744 | C10H24O4S | [M+H]+ | + | 2.641 | x | |
| 241.92815 | C8H2O8S | [M-NH3+H]+ | + | 1.492 | x | |
| 242.02701 | C13H9O3NS | [M-H2O+H]+ | + | -0.060 | x | |
| 242.07883 | C7H18O7NP | [M-H2O+H]+ | + | 0.117 | x | |
| 242.09972 | C15H17ONS | [M-H2O+H]+ | + | -0.294 | x | |
| 242.9776 | C5H5O6N2P | [M+Na]+ | + | -0.652 | x | |
| 242.9842 | C12H5O5P | [M-H2O+H]+ | + | 0.110 | x | |
| 243.05392 | C8H20O3S3 | [M-H2O+H]+ | + | -0.953 | x | |
| 243.08375 | C15H16O2S | [M-H2O+H]+ | + | -0.238 | x | |
| 243.10312 | C16H12N4 | [M-H2O+H]+ | + | 0.815 | x | |
| 243.92503 | C8H4O6S2 | [M-NH3+H]+ | + | -2.411 | x | |
| 243.96153 | C13HN3P2 | [M-H2O+H]+ | + | 0.950 | x | |
| 244.06109 | C6H11O3N7S | [M-H2O+H]+ | + | -0.113 | x | |
| 244.08547 | C15H16O4 | [M-NH3+H]+ | + | -0.446 | x | |
| 244.97443 | C10H3O5N2P | [M-H2O+H]+ | + | -0.914 | x | |
| 245.03319 | C7H18O4S3 | [M-H2O+H]+ | + | -0.925 | x | |

| | | | | | |
|---|---|---|---|---|---|
| 245.04509 | C6H10O4N6S | [M-H2O+H]+ | + | -0.172 | x |
| 245.06948 | C8H20O2S3 | [M+H]+ | + | -1.384 | x |
| 245.0783 | C18H14S | [M-H2O+H]+ | + | -0.125 | x |
| 245.09937 | C7H14O5N6 | [M-H2O+H]+ | + | 0.347 | x |
| 245.95848 | C10H2O6NP | [M-H2O+H]+ | + | -0.780 | x |
| 245.98555 | C11H5O5NS | [M-H2O+H]+ | + | -0.018 | x |
| 246.11856 | C7H13N9 | [M+Na]+ | + | -0.234 | x |
| 246.96974 | C11H4O6S | [M-H2O+H]+ | + | 0.643 | x |
| 246.97332 | C8O7N4 | [M-H2O+H]+ | + | -0.342 | x |
| 247.04884 | C7H18O3S3 | [M+H]+ | + | -0.986 | x |
| 247.08516 | C10H13ON6P | [M-H2O+H]+ | + | -1.504 | x |
| 247.12885 | C10H20O6N2 | [M-H2O+H]+ | + | 0.009 | x |
| 248.01958 | C8H8O3N3P | [M+Na]+ | + | 0.140 | x |
| 248.10724 | C17H15O2N | [M-H2O+H]+ | + | 0.942 | x |
| 248.94045 | C12O2N2P2 | [M-H2O+H]+ | + | 0.894 | x |
| 249.00358 | C8H7O4N2P | [M+Na]+ | + | 0.070 | x |
| 249.02807 | C6H16O4S3 | [M+H]+ | + | -1.117 | x |
| 249.0645 | C9H11O2N6P | [M-H2O+H]+ | + | -1.209 | x |
| 249.0912 | C10H22N2S3 | [M-H2O+H]+ | + | -0.083 | x |
| 249.09438 | C14H18O3S | [M-H2O+H]+ | + | 0.013 | x |
| 249.15248 | C12H26O4S | [M-H2O+H]+ | + | 2.213 | x |
| 251.12504 | C10H16O3N6 | [M-H2O+H]+ | + | -0.223 | x |
| 251.97809 | C6H4O5N3P | [M+Na]+ | + | 0.053 | x |
| 252.05722 | C15H12O2NP | [M-H2O+H]+ | + | -0.211 | x |
| 252.12042 | C9H15O3N7 | [M-H2O+H]+ | + | 0.265 | x |
| 252.13176 | C15H26P2 | [M-NH3+H]+ | + | 0.222 | x |
| 252.14402 | C10H23O7N | [M-H2O+H]+ | + | -0.532 | x |
| 252.94469 | C10H6O3S3 | [M-H2O+H]+ | + | 0.267 | x |
| 252.96211 | C6H3O6N2P | [M+Na]+ | + | 0.072 | x |
| 252.9737 | C9H6O4N2S2 | [M-H2O+H]+ | + | 0.334 | x |
| 253.04121 | C15H11O3P | [M-H2O+H]+ | + | -0.286 | x |
| 253.10445 | C9H14O4N6 | [M-H2O+H]+ | + | 0.317 | x |
| 253.12806 | C10H22O8 | [M-H2O+H]+ | + | -0.440 | x |
| 253.14081 | C10H18O3N6 | [M-H2O+H]+ | + | 0.223 | x |
| 253.26376 | C16H34ON2 | [M-H2O+H]+ | + | -0.239 | x |
| 254.03657 | C14H10O3NP | [M-H2O+H]+ | + | 0.106 | x |
| 254.13611 | C17H21NS | [M-H2O+H]+ | + | -0.264 | x |
| 254.15974 | C10H25O7N | [M-H2O+H]+ | + | -0.270 | x |
| 255.12007 | C9H16O4N6 | [M-H2O+H]+ | + | 0.204 | x |
| 255.91053 | C5H3N3S4 | [M+Na]+ | + | 1.412 | x |
| 256.03337 | C11H12O8 | [M-NH3+H]+ | + | -2.111 | x |
| 256.08547 | C16H16O4 | [M-NH3+H]+ | + | -0.426 | x |
| 256.08866 | C15H16O2NP | [M-H2O+H]+ | + | 0.305 | x |
| 256.11533 | C8H15O4N7 | [M-H2O+H]+ | + | 0.244 | x |
| 256.13894 | C9H23O8N | [M-H2O+H]+ | + | -0.504 | x |
| 256.1753 | C10H25O6N | [M+H]+ | + | -0.640 | x |
| 256.89458 | C5H2ON2S4 | [M+Na]+ | + | 1.553 | x |
| 256.97436 | C11H3O5N2P | [M-H2O+H]+ | + | -1.130 | x |
| 256.99982 | C13H7O5P | [M-H2O+H]+ | + | -0.005 | x |
| 258.06473 | C15H14O5 | [M-NH3+H]+ | + | -0.440 | x |
| 258.07365 | C12H13O4N | [M+Na]+ | + | -0.122 | x |
| 258.08544 | C6H18O7N3P | [M-H2O+H]+ | + | 1.790 | x |
| 258.10994 | C13H17O3N | [M+Na]+ | + | -0.528 | x |
| 258.15463 | C9H23O7N | [M+H]+ | + | -0.380 | x |

| | | | | | |
|---|---|---|---|---|---|
| 258.89149 | C4O4N2S3 | [M+Na]+ | + | 1.061 | x |
| 258.92479 | C8O3N2S2 | [M+Na]+ | + | 2.270 | x |
| 258.92771 | C5H4O3N2S3 | [M+Na]+ | + | 0.359 | x |
| 258.95478 | C7H2O2N4P2 | [M+Na]+ | + | 1.104 | x |
| 259.0156 | C13H9O5P | [M-H2O+H]+ | + | 0.466 | x |
| 259.04877 | C8H20O4S3 | [M-H2O+H]+ | + | -1.132 | x |
| 259.0577 | C12H12O5 | [M+Na]+ | + | 0.024 | x |
| 259.06889 | C11H12O4N2 | [M+Na]+ | + | -0.160 | x |
| 259.08527 | C9H22O2S3 | [M+H]+ | + | -0.767 | x |
| 259.11532 | C10H20O6 | [M+Na]+ | + | 0.471 | x |
| 259.15385 | C13H24O6 | [M-H2O+H]+ | + | -0.542 | x |
| 259.16566 | C9H25ON4P | [M+Na]+ | + | -0.671 | x |
| 259.17318 | C7H23N5S2 | [M+NH4]+ | + | -0.527 | x |
| 260.08051 | C15H16O5 | [M-NH3+H]+ | + | 0.034 | x |
| 260.89037 | C6H2O3N2S4 | [M-H2O+H]+ | + | -4.198 | x |
| 260.93296 | C9O6N3P | [M-NH3+H]+ | + | -0.862 | x |
| 260.95158 | C12ON4P2 | [M-H2O+H]+ | + | 0.484 | x |
| 261.07337 | C7H20O8NP | [M-NH3+H]+ | + | -0.036 | x |
| 261.09441 | C15H18O3S | [M-H2O+H]+ | + | 0.120 | x |
| 261.1009 | C11H15ON6P | [M-H2O+H]+ | + | -1.104 | x |
| 261.13067 | C8H18O5N6 | [M-H2O+H]+ | + | 0.327 | x |
| 261.18082 | C12H26O5N2 | [M-H2O+H]+ | + | -0.226 | x |
| 262.09607 | C15H18O5 | [M-NH3+H]+ | + | -0.290 | x |
| 262.13417 | C17H17ON3 | [M-H2O+H]+ | + | 1.062 | x |
| 262.95051 | C7H6O8P2 | [M-H2O+H]+ | + | 0.029 | x |
| 263.04374 | C7H18O4S3 | [M+H]+ | + | -0.981 | x |
| 263.0801 | C7H12N8S | [M+Na]+ | + | 1.316 | x |
| 263.12522 | C19H20S | [M-H2O+H]+ | + | -0.224 | x |
| 263.14881 | C12H24O7 | [M-H2O+H]+ | + | -0.390 | x |
| 264.02074 | C15H8O3NP | [M-H2O+H]+ | + | -0.538 | x |
| 264.08302 | C10H22O2NP3 | [M-H2O+H]+ | + | -0.106 | x |
| 264.16775 | C10H19ON9 | [M-H2O+H]+ | + | -0.774 | x |
| 265.02298 | C6H16O5S3 | [M+H]+ | + | -1.067 | x |
| 265.10423 | C10H14O4N6 | [M-H2O+H]+ | + | -0.476 | x |
| 265.14083 | C11H18O3N6 | [M-H2O+H]+ | + | 0.284 | x |
| 265.17686 | C12H30ON2S2 | [M-H2O+H]+ | + | 0.688 | x |
| 266.03646 | C15H10O3NP | [M-H2O+H]+ | + | -0.287 | x |
| 266.13616 | C18H21NS | [M-H2O+H]+ | + | -0.077 | x |
| 266.17255 | C13H25O3N | [M+Na]+ | + | -0.469 | x |
| 266.23251 | C13H31O4N | [M+H]+ | + | -0.279 | x |
| 266.95083 | C12H2O3N2P2 | [M-H2O+H]+ | + | 0.187 | x |
| 267.06805 | C15H13O2N2P | [M-H2O+H]+ | + | -0.442 | x |
| 267.12005 | C10H16O4N6 | [M-H2O+H]+ | + | 0.125 | x |
| 267.14371 | C11H24O8 | [M-H2O+H]+ | + | -0.418 | x |
| 267.15661 | C19H24S | [M-H2O+H]+ | + | 0.096 | x |
| 268.05205 | C15H12O3NP | [M-H2O+H]+ | + | -0.496 | x |
| 268.07147 | C14H11O4N3 | [M-H2O+H]+ | + | -0.692 | x |
| 268.09443 | C14H15O3N | [M+Na]+ | + | 0.065 | x |
| 268.13892 | C10H23O8N | [M-H2O+H]+ | + | -0.553 | x |
| 269.03616 | C20H6 | [M+Na]+ | + | -0.046 | x |
| 269.06498 | C6H15O7N4P | [M-H2O+H]+ | + | 1.515 | x |
| 269.13577 | C18H22OS | [M-H2O+H]+ | + | -0.253 | x |
| 269.15273 | C12H23O2N4P | [M-H2O+H]+ | + | 0.546 | x |
| 269.97641 | C8HN9S2 | [M-H2O+H]+ | + | 0.229 | x |

| | | | | | | |
|---|---|---|---|---|---|---|
| 270.04897 | C4H17ON5S3 | [M+Na]+ | + | 0.919 | x | |
| 270.13099 | C9H17O4N7 | [M-H2O+H]+ | + | 0.267 | x | |
| 270.15459 | C10H25O8N | [M-H2O+H]+ | + | -0.480 | x | |
| 270.98434 | C19HN2P | [M-H2O+H]+ | + | -0.370 | x | |
| 271.03296 | C4H16O2N4S3 | [M+Na]+ | + | 0.812 | x | |
| 271.05209 | C15H13O4P | [M-H2O+H]+ | + | 0.809 | x | |
| 271.06378 | C12H16O6S | [M-H2O+H]+ | + | 1.075 | x | |
| 271.158 | C11H26O5S | [M+H]+ | + | 2.331 | x | |
| 271.20669 | C19H29ON | [M+NH3-H]- | - | -0.167 | x | x |
| 271.9231 | C5H3O3N3S3 | [M+Na]+ | + | 0.907 | x | |
| 272.04777 | C11H13N3P2 | [M+Na]+ | + | 0.316 | x | |
| 272.08039 | C16H16O5 | [M-NH3+H]+ | + | -0.384 | x | |
| 272.11691 | C17H20O4 | [M-NH3+H]+ | + | 0.083 | x | |
| 272.17028 | C10H25O7N | [M+H]+ | + | -0.361 | x | |
| 273.03175 | C11H12ON2P2 | [M+Na]+ | + | 0.172 | x | |
| 273.06446 | C9H22O4S3 | [M-H2O+H]+ | + | -0.939 | x | |
| 273.09435 | C16H18O3S | [M-H2O+H]+ | + | -0.091 | x | |
| 273.10088 | C10H24O2S3 | [M+H]+ | + | -0.874 | x | |
| 273.1672 | C12H26O5 | [M+Na]+ | + | -0.177 | x | |
| 273.9211 | C5H5ON3S4 | [M+Na]+ | + | 1.332 | x | |
| 274.09617 | C16H18O5 | [M-NH3+H]+ | + | 0.067 | x | |
| 274.90515 | C13S3 | [M+Na]+ | + | -1.125 | x | |
| 274.93202 | C9H8O3S4 | [M-H2O+H]+ | + | -1.091 | x | |
| 275.08012 | C9H22O3S3 | [M+H]+ | + | -0.940 | x | |
| 275.10999 | C16H20O3S | [M-H2O+H]+ | + | -0.125 | x | |
| 275.16149 | C13H20O2N6 | [M-H2O+H]+ | + | 0.016 | x | |
| 275.91816 | C4H3O4N3S3 | [M+Na]+ | + | 1.468 | x | |
| 276.07532 | C15H16O6 | [M-NH3+H]+ | + | -0.326 | x | |
| 276.18043 | C13H27O6N | [M-H2O+H]+ | + | -0.405 | x | |
| 276.90212 | C10H2ON2S4 | [M-H2O+H]+ | + | 1.404 | x | |
| 276.93549 | C13O3N2P2 | [M-H2O+H]+ | + | 1.236 | x | |
| 277.05936 | C8H20O4S3 | [M+H]+ | + | -1.040 | x | |
| 277.09583 | C11H15O2N6P | [M-H2O+H]+ | + | -0.992 | x | |
| 277.1047 | C8H21O8P | [M+H]+ | + | 0.091 | x | |
| 277.1405 | C12H26O2N2S2 | [M-H2O+H]+ | + | 0.746 | x | |
| 277.91695 | C5H10O6S4 | [M-NH3+H]+ | + | 0.687 | x | |
| 278.19612 | C13H29O6N | [M-H2O+H]+ | + | -0.266 | x | |
| 278.90092 | C12OS3 | [M+Na]+ | + | 2.235 | x | |
| 279.07508 | C10H13O3N6P | [M-H2O+H]+ | + | -1.035 | x | |
| 279.1437 | C12H24O8 | [M-H2O+H]+ | + | -0.435 | x | |
| 280.05217 | C16H12O3NP | [M-H2O+H]+ | + | -0.072 | x | |
| 280.15184 | C19H23NS | [M-H2O+H]+ | + | 0.028 | x | |
| 280.93962 | C11H6O4S3 | [M-H2O+H]+ | + | 0.310 | x | |
| 281.04729 | C15H11O3N2P | [M-H2O+H]+ | + | -0.504 | x | |
| 281.13568 | C11H18O4N6 | [M-H2O+H]+ | + | 0.052 | x | |
| 281.17226 | C20H26S | [M-H2O+H]+ | + | 0.092 | x | |
| 282.03135 | C20H5N | [M+Na]+ | + | -0.271 | x | |
| 282.0491 | C5H17ON5S3 | [M+Na]+ | + | 1.378 | x | |
| 282.131 | C10H17O4N7 | [M-H2O+H]+ | + | 0.290 | x | |
| 282.15463 | C11H25O8N | [M-H2O+H]+ | + | -0.327 | x | |
| 283.11496 | C10H16O5N6 | [M-H2O+H]+ | + | 0.103 | x | |
| 283.17502 | C12H28O8 | [M-H2O+H]+ | + | -0.362 | x | |
| 283.19026 | C16H28O5 | [M-H2O+H]+ | + | -0.416 | x | |
| 284.02819 | C4H15O2N5S3 | [M+Na]+ | + | 0.699 | x | |

| | | | | | |
|---|---|---|---|---|---|
| 284.04748 | C13H16O6S | [M-NH3+H]+ | + | -0.020 | x |
| 284.17026 | C11H27O8N | [M-H2O+H]+ | + | -0.391 | x |
| 285.13084 | C12H22O6 | [M+Na]+ | + | -0.072 | x |
| 285.17369 | C12H30O3P2 | [M+H]+ | + | -2.123 | x |
| 285.20591 | C16H30O5 | [M-H2O+H]+ | + | -0.414 | x |
| 286.02684 | C12H14O7S | [M-NH3+H]+ | + | 0.296 | x |
| 286.0961 | C17H18O5 | [M-NH3+H]+ | + | -0.167 | x |
| 286.18596 | C11H27O7N | [M+H]+ | + | -0.238 | x |
| 287.08012 | C10H24O4S3 | [M-H2O+H]+ | + | -0.863 | x |
| 288.07533 | C16H16O6 | [M-NH3+H]+ | + | -0.280 | x |
| 288.16518 | C10H25O8N | [M+H]+ | + | -0.391 | x |
| 289.08928 | C16H18O4S | [M-H2O+H]+ | + | -0.036 | x |
| 289.09576 | C10H24O3S3 | [M+H]+ | + | -0.946 | x |
| 289.12575 | C11H22O7 | [M+Na]+ | + | -0.088 | x |
| 290.09103 | C16H18O6 | [M-NH3+H]+ | + | -0.115 | x |
| 291.07505 | C9H22O4S3 | [M+H]+ | + | -0.852 | x |
| 291.08399 | C8H22O9NP | [M-NH3+H]+ | + | 0.132 | x |
| 291.10489 | C16H20O4S | [M-H2O+H]+ | + | -0.150 | x |
| 291.11146 | C12H17O2N6P | [M-H2O+H]+ | + | -1.012 | x |
| 291.15647 | C13H20O3N6 | [M-H2O+H]+ | + | 0.228 | x |
| 293.05426 | C8H20O5S3 | [M+H]+ | + | -1.033 | x |
| 293.13565 | C12H18O4N6 | [M-H2O+H]+ | + | -0.046 | x |
| 293.20881 | C16H30O3 | [M+Na]+ | + | 0.351 | x |
| 294.22742 | C14H33O6N | [M-H2O+H]+ | + | -0.253 | x |
| 295.19033 | C17H28O5 | [M-H2O+H]+ | + | -0.176 | x |
| 296.17024 | C12H27O8N | [M-H2O+H]+ | + | -0.440 | x |
| 296.2583 | C18H35O3N | [M-H2O+H]+ | + | -0.335 | x |
| 296.90374 | C11HO3P3 | [M+Na]+ | + | 2.411 | x |
| 297.15429 | C12H26O9 | [M-H2O+H]+ | + | -0.329 | x |
| 297.88772 | C11HNS4 | [M+Na]+ | + | -2.485 | x |
| 298.14949 | C11H25O9N | [M-H2O+H]+ | + | -0.484 | x |
| 298.18592 | C12H29O8N | [M-H2O+H]+ | + | -0.342 | x |
| 298.87171 | C11OS4 | [M+Na]+ | + | -2.569 | x |
| 298.90054 | C8H4N4S5 | [M-H2O+H]+ | + | -0.385 | x |
| 299.05791 | C20H8N2 | [M+Na]+ | + | -0.233 | x |
| 299.16167 | C17H24O3 | [M+Na]+ | + | -0.344 | x |
| 299.18286 | C14H28O5 | [M+Na]+ | + | -0.124 | x |
| 299.88461 | C8H3ON3S5 | [M-H2O+H]+ | + | -0.212 | x |
| 300.04203 | C15H12O5NP | [M-H2O+H]+ | + | 0.030 | x |
| 300.16516 | C11H27O9N | [M-H2O+H]+ | + | -0.417 | x |
| 300.20153 | C12H29O7N | [M+H]+ | + | -0.494 | x |
| 300.21681 | C16H31O5N | [M-H2O+H]+ | + | -0.391 | x |
| 300.86861 | C8H2O2N2S5 | [M-H2O+H]+ | + | -0.260 | x |
| 301.0593 | C9H10ON8S | [M+Na]+ | + | 0.904 | x |
| 301.12574 | C12H22O7 | [M+Na]+ | + | -0.120 | x |
| 301.14095 | C7H26N8S3 | [M-H2O+H]+ | + | -0.051 | x |
| 301.20494 | C18H31P | [M+Na]+ | + | -2.221 | x |
| 301.88332 | C10HONS4 | [M+Na]+ | + | 0.008 | x |
| 302.18085 | C11H27O8N | [M+H]+ | + | -0.307 | x |
| 302.2325 | C16H33O5N | [M-H2O+H]+ | + | -0.263 | x |
| 303.10493 | C17H20O4S | [M-H2O+H]+ | + | -0.035 | x |
| 304.07025 | C16H16O7 | [M-NH3+H]+ | + | -0.249 | x |
| 304.2117 | C15H31O6N | [M-H2O+H]+ | + | -0.463 | x |
| 305.15699 | C18H26O3S | [M-H2O+H]+ | + | 0.042 | x |

| | | | | | | |
|---|---|---|---|---|---|---|
| 310.23748 | C18H33O4N | [M-H2O+H]+ | + | -0.579 | x | |
| 311.16991 | C13H28O9 | [M-H2O+H]+ | + | -0.406 | x | |
| 311.1852 | C17H28O6 | [M-H2O+H]+ | + | -0.303 | x | |
| 312.16516 | C12H27O9N | [M-H2O+H]+ | + | -0.402 | x | |
| 312.21682 | C17H31O5N | [M-H2O+H]+ | + | -0.331 | x | |
| 312.25322 | C18H35O4N | [M-H2O+H]+ | + | -0.302 | x | |
| 314.18078 | C12H29O9N | [M-H2O+H]+ | + | -0.490 | x | |
| 315.18421 | C16H25N6P | [M-H2O+H]+ | + | -1.001 | x | |
| 316.19644 | C12H29O8N | [M+H]+ | + | -0.484 | x | |
| 316.21168 | C16H31O6N | [M-H2O+H]+ | + | -0.506 | x | |
| 317.17215 | C23H26S | [M-H2O+H]+ | + | -0.247 | x | |
| 317.1926 | C11H25O8N | [M-NH4]- | - | -1.131 | x | |
| 317.19647 | C16H31O6N | [M+NH3-H]- | - | -1.474 | x | x |
| 319.13623 | C18H24O4S | [M-H2O+H]+ | + | -0.033 | x | |
| 321.13083 | C15H22O6 | [M+Na]+ | + | -0.097 | x | |
| 323.90352 | C18N2S3 | [M-NH3+H]+ | + | 1.286 | x | |
| 326.19607 | C17H29O6N | [M-H2O+H]+ | + | -0.375 | x | |
| 326.88444 | C10H4O2N2S5 | [M-H2O+H]+ | + | 0.283 | x | |
| 327.06867 | C12H16O9 | [M+Na]+ | + | 0.073 | x | |
| 327.17988 | C9H24N10S | [M+Na]+ | + | 0.174 | x | |
| 327.2012 | C14H32O9 | [M-H2O+H]+ | + | -0.417 | x | |
| 327.24979 | C14H31O6N | [M-NH4]- | - | -0.870 | x | |
| 328.16008 | C12H27N5S2 | [M+Na]+ | + | 0.237 | x | |
| 328.19643 | C13H31O9N | [M-H2O+H]+ | + | -0.470 | x | |
| 328.2117 | C17H31O6N | [M-H2O+H]+ | + | -0.430 | x | |
| 328.24815 | C18H35O5N | [M-H2O+H]+ | + | -0.243 | x | |
| 329.19988 | C17H27N6P | [M-H2O+H]+ | + | -0.903 | x | |
| 330.17571 | C12H29N5S2 | [M+Na]+ | + | 0.170 | x | |
| 330.21217 | C13H31O8N | [M+H]+ | + | -0.220 | x | |
| 330.22697 | C17H33O6N | [M-H2O+H]+ | + | -1.522 | x | |
| 330.26373 | C18H37O5N | [M-H2O+H]+ | + | -0.443 | x | |
| 330.26871 | C23H38O2 | [M+NH3-H]- | - | -0.848 | x | |
| 331.09975 | C10H16O8N6 | [M-H2O+H]+ | + | 0.222 | x | |
| 331.15154 | C8H28ON8S3 | [M-H2O+H]+ | + | 0.027 | x | |
| 332.1914 | C12H31N5S2 | [M+Na]+ | + | 0.299 | x | |
| 333.15195 | C13H26O8 | [M+Na]+ | + | -0.123 | x | |
| 333.16712 | C23H26OS | [M-H2O+H]+ | + | -0.092 | x | |
| 336.25088 | C18H35O3N | [M+Na]+ | + | -0.109 | x | |
| 337.2012 | C13H32O4N5P | [M+NH3-H]- | - | 0.526 | x | x |
| 338.26652 | C18H37O3N | [M+Na]+ | + | -0.140 | x | |
| 340.19643 | C14H31O9N | [M-H2O+H]+ | + | -0.455 | x | |
| 341.24334 | C18H34O5N2 | [M-H2O+H]+ | + | -0.399 | x | |
| 342.17571 | C13H29N5S2 | [M+Na]+ | + | 0.164 | x | |
| 344.19136 | C13H31N5S2 | [M+Na]+ | + | 0.163 | x | |
| 344.20661 | C17H31O7N | [M-H2O+H]+ | + | -0.424 | x | |
| 344.22773 | C14H33O8N | [M+H]+ | + | -0.473 | x | |
| 344.2429 | C18H35O6N | [M-H2O+H]+ | + | -0.688 | x | |
| 345.15197 | C14H26O8 | [M+Na]+ | + | -0.056 | x | |
| 345.2312 | C18H31N6P | [M-H2O+H]+ | + | -0.807 | x | |
| 347.13118 | C19H24O5S | [M-H2O+H]+ | + | 0.067 | x | |
| 347.17016 | C10H30O7N5P | [M+NH3-H]- | - | 0.143 | x | |
| 347.95362 | C9H11ON5S5 | [M-H2O+H]+ | + | 0.548 | x | |
| 349.05994 | C18H15O4P | [M+Na]+ | + | -0.234 | x | |
| 349.18324 | C14H30O8 | [M+Na]+ | + | -0.132 | x | |

| | | | | | | |
|---|---|---|---|---|---|---|
| 350.23017 | C18H33O4N | [M+Na]+ | + | -0.027 | x | |
| 356.1914 | C12H19N15 | [M-H2O+H]+ | + | -0.260 | x | |
| 358.2071 | C12H21N15 | [M-H2O+H]+ | + | -0.125 | x | |
| 360.18635 | C11H14N14 | [M+NH4]+ | + | -0.180 | x | |
| 360.22273 | C12H23N15 | [M-H2O+H]+ | + | -0.178 | x | |
| 363.16255 | C14H28O9 | [M+Na]+ | + | -0.008 | x | |
| 367.21121 | C17H25N11 | [M+NH3-H]- | - | -0.139 | x | x |
| 369.22682 | C17H27N11 | [M+NH3-H]- | - | -0.242 | x | x |
| 371.2275 | C14H24N14 | [M-H2O+H]+ | + | -0.124 | x | |
| 372.18638 | C12H19ON15 | [M-H2O+H]+ | + | -0.081 | x | |
| 372.22277 | C13H23N15 | [M-H2O+H]+ | + | -0.069 | x | |
| 374.202 | C12H21ON15 | [M-H2O+H]+ | + | -0.157 | x | |
| 375.2489 | C18H36O7N2 | [M-H2O+H]+ | + | -0.159 | x | |
| 383.20577 | C17H33N7S2 | [M+NH3-H]- | - | 0.153 | x | x |
| 384.22281 | C16H35O10N | [M-H2O+H]+ | + | 0.007 | x | |
| 385.22141 | C17H35N7S2 | [M+NH3-H]- | - | 0.128 | x | x |
| 386.20206 | C13H21ON15 | [M-H2O+H]+ | + | -0.004 | x | |
| 388.18126 | C12H19O2N15 | [M-H2O+H]+ | + | -0.163 | x | |
| 388.21772 | C15H35O11N | [M-H2O+H]+ | + | -0.004 | x | |
| 388.25415 | C16H37O9N | [M+H]+ | + | 0.111 | x | |
| 389.25753 | C20H35ON6P | [M-H2O+H]+ | + | -0.436 | x | |
| 391.15752 | C15H28O10 | [M+Na]+ | + | 0.143 | x | |
| 393.2976 | C22H42O4 | [M+Na]+ | + | 0.190 | x | |
| 397.23081 | C24H34O2N2S | [M-H2O+H]+ | + | 0.000 | x | |
| 400.21766 | C14H23ON15 | [M-H2O+H]+ | + | -0.124 | x | |
| 402.23333 | C14H25ON15 | [M-H2O+H]+ | + | -0.075 | x | |
| 407.18874 | C14H20N14 | [M+Na]+ | + | -0.032 | x | |
| 416.21267 | C16H35O12N | [M-H2O+H]+ | + | 0.078 | x | |
| 416.24908 | C22H37N5P2 | [M-H2O+H]+ | + | -0.115 | x | |
| 428.24895 | C16H27ON15 | [M-H2O+H]+ | + | -0.138 | x | |
| 432.28026 | C16H31ON15 | [M-H2O+H]+ | + | -0.115 | x | |
| 446.25966 | C23H39ON5P2 | [M-H2O+H]+ | + | -0.075 | x | |
| 476.30641 | C20H47O2N5S2 | [M+Na]+ | + | 0.150 | x | |

[a] Molecular formula based on accurate mass only (no isotopic pattern matching performed).

x = Presence of a particular *m/z* in the respective bacterial serotype.

NA= Not available
